# Dating in the Dark: Elevated Substitution Rates in Cave Cockroaches (Blattodea: Nocticolidae) Have Negative Impacts on Molecular Date Estimates

**DOI:** 10.1101/2023.01.17.524483

**Authors:** Toby G. L. Kovacs, James Walker, Simon Hellemans, Thomas Bourguignon, Nikolai J. Tatarnic, Jane M. Mcrae, Simon Y. W. Ho, Nathan Lo

## Abstract

Rates of nucleotide substitution vary substantially across the Tree of Life, with potentially confounding effects on phylogenetic and evolutionary analyses. A large acceleration in mitochondrial substitution rate occurs in the cockroach family Nocticolidae, which predominantly inhabit subterranean environments. To evaluate the impacts of this among-lineage rate heterogeneity on estimates of phylogenetic relationships and evolutionary timescales, we analysed nuclear ultraconserved elements (UCEs) and mitochondrial genomes from nocticolids and other cockroaches. Substitution rates were substantially elevated in nocticolid lineages compared with other cockroaches, especially in mitochondrial protein-coding genes. This disparity in evolutionary rates is likely to have led to different evolutionary relationships being supported by phylogenetic analyses of mitochondrial genomes and UCE loci. Furthermore, Bayesian dating analyses using relaxed-clock models inferred much deeper divergence times compared with a flexible local clock. Our phylogenetic analysis of UCEs, which is the first genome-scale study to include all ten major cockroach families, unites Corydiidae and Nocticolidae and places Anaplectidae as the sister lineage to the rest of Blattoidea. We uncover an extraordinary level of genetic divergence in Nocticolidae, including two highly distinct clades that separated ∼115 million years ago despite both containing representatives of the genus *Nocticola*. The results of our study highlight the potential impacts of high among-lineage rate variation on estimates of phylogenetic relationships and evolutionary timescales.

---

Rates of molecular evolution vary by several orders of magnitude across the Tree of Life. This among-lineage rate variation has been linked to differences in life-history traits such as body size, population size, longevity, metabolic rate, and generation time (Bromham 2009). Remarkably, large disparities in evolutionary rates can even be found between closely related lineages, such as in burrowing crayfish (Gan et al. 2018), crabronid wasps (Kaltenpoth et al. 2012), and parasitic plants (Lemaire et al. 2011; Bromham et al. 2013).

When molecular evolutionary rates show extreme or complex variation across lineages, they can pose difficulties for phylogenetic inference and molecular dating (Kolaczkowski and Thornton 2004, 2009; Dornburg et al. 2012; Crisp et al. 2014; Susko 2015; Roch et al. 2019). A range of clock models have been developed in order to account for among-lineage rate variation in Bayesian phylogenetic dating analyses. These include local clocks, in which different rates are allowed in groups of neighbouring branches, and relaxed clocks, in which a different rate is allowed along each branch (Ho and Duchêne 2014). The placement of local clocks can either be selected a priori or jointly estimated with the phylogeny, which is possible using the random local clock (Drummond and Suchard 2010) and the shrinkage-based random local clock (Fisher et al. 2021). The most widely used relaxed clocks either treat the branch rates as autocorrelated (Thorne et al. 1998; Sanderson 2002; Lepage et al. 2006), where they are assumed to be related between adjacent branches, or uncorrelated, where they are drawn independently from a chosen parametric distribution (Drummond et al. 2006; Rannala and Yang 2007). These relaxed clocks typically rely on the branch rates fitting a simple unimodal distribution and might provide a poor fit under conditions of complex rate variation.

Although rate variation is ubiquituous across the tree of life (Ho 2020), few studies have focused on the behaviour and performance of clock models when challenged with extreme among-lineage rate variation. Studies of simulations have demonstrated that misspecification of relaxed-clock models can lead to imprecise estimates of evolutionary rates and timescales (Dornburg et al. 2012; Duchêne et al. 2014; Fourment and Darling 2018). There has also been varying support for uncorrelated and autocorrelated relaxed clocks, with substantial impacts on the inference of evolutionary histories (Lepage et al. 2007; Linder et al. 2011; Ho et al. 2015; dos Reis et al. 2018). When applied to phylogenies with among-lineage rate variation, uncorrelated relaxed clocks tend to infer a distribution of substitution rates centred on the average rate across the tree, leading to poor estimation of divergence times (Dornburg et al. 2012; Crisp et al. 2014). Data sets containing such rate variation produce more intuitive date estimates when analysed using a random local clock (Dornburg et al. 2012; Crisp et al. 2014), although this might depend on the distribution and extent of rate variation across the tree (Gan et al. 2018). Despite its potential utility, the random local clock can be difficult to implement and so other clock models are usually preferred in practice (Drummond and Suchard 2010; Dornburg et al. 2012).

The flexible local clock combines some of the features of local clocks and relaxed clocks by allowing multiple, independent uncorrelated relaxed clocks for different clades across the tree (Fourment and Darling 2018). This model can capture large-scale rate disparities among clades while also accounting for local variation in rates. Although the flexible local clock provides an interesting framework that allows distinct evolutionary rates between clades, it has not been widely employed. This is partly due to the challenge of selecting local clocks a priori, but possibly also because its implementation is non-trivial. To date, there has been no study that has compared the behaviour of a diverse range of clock models, including local clocks and relaxed clocks, when challenged by extreme rate variation among lineages.

One group of organisms that appears to have experienced a dramatic shift in mitochondrial substitution rates is the cockroach family Nocticolidae (Lo et al. 2007; Legendre et al. 2008, 2015; Djernæs et al. 2015; Wang et al. 2017; Bourguignon et al. 2018; Li and Huang 2020). The causes and timing of this rate elevation, and whether such a pattern also occurs in the nuclear genome of nocticolids, are unknown. Dominated by cavernicolous taxa exhibiting troglomorphic (cave adapted) characters, nocticolids are small (<10 mm body length), pale, and delicate, and often have absent or reduced eyes and wings (Fig. 1). There are 38 described species of Nocticolidae across nine genera, found in Africa, Asia, and Oceania (Supplementary Table S1). The family is in desperate need of taxonomic revision, with 26 species placed in a poorly defined genus (*Nocticola*) accompanied by six monotypic genera, a genus (*Alluaudellina*) that is likely to be a synonym of *Nocticola* (Chopard 1932; Roth 1988), and a genus (*Spelaeoblatta*) containing four species only from Thailand and Myanmar (Roth and McGavin 1994; Vidlička et al. 2003). There are even doubts that four of the genera (including *Spelaeoblatta*) should be included in the family at all, based on morphology (Li and Huang 2020). The majority (24) of species are subterranean, including three from non-cave subterranean environments (Trotter et al. 2017) and two that are known to move in and out of caves. Ten species are epigean and three are termitophilous (living within termite nests). Epigean taxa typically have fully developed wings, compound eyes, and dark pigmentation, while subterranean taxa rarely possess these features. Owing to their small size, great speed, and inaccessible habitats, nocticolids remain the most poorly studied of the ten recognized cockroach families (Karny 1924; Fernando 1957; Asahina 1974; Roth 1988).

**FIGURE 1.**
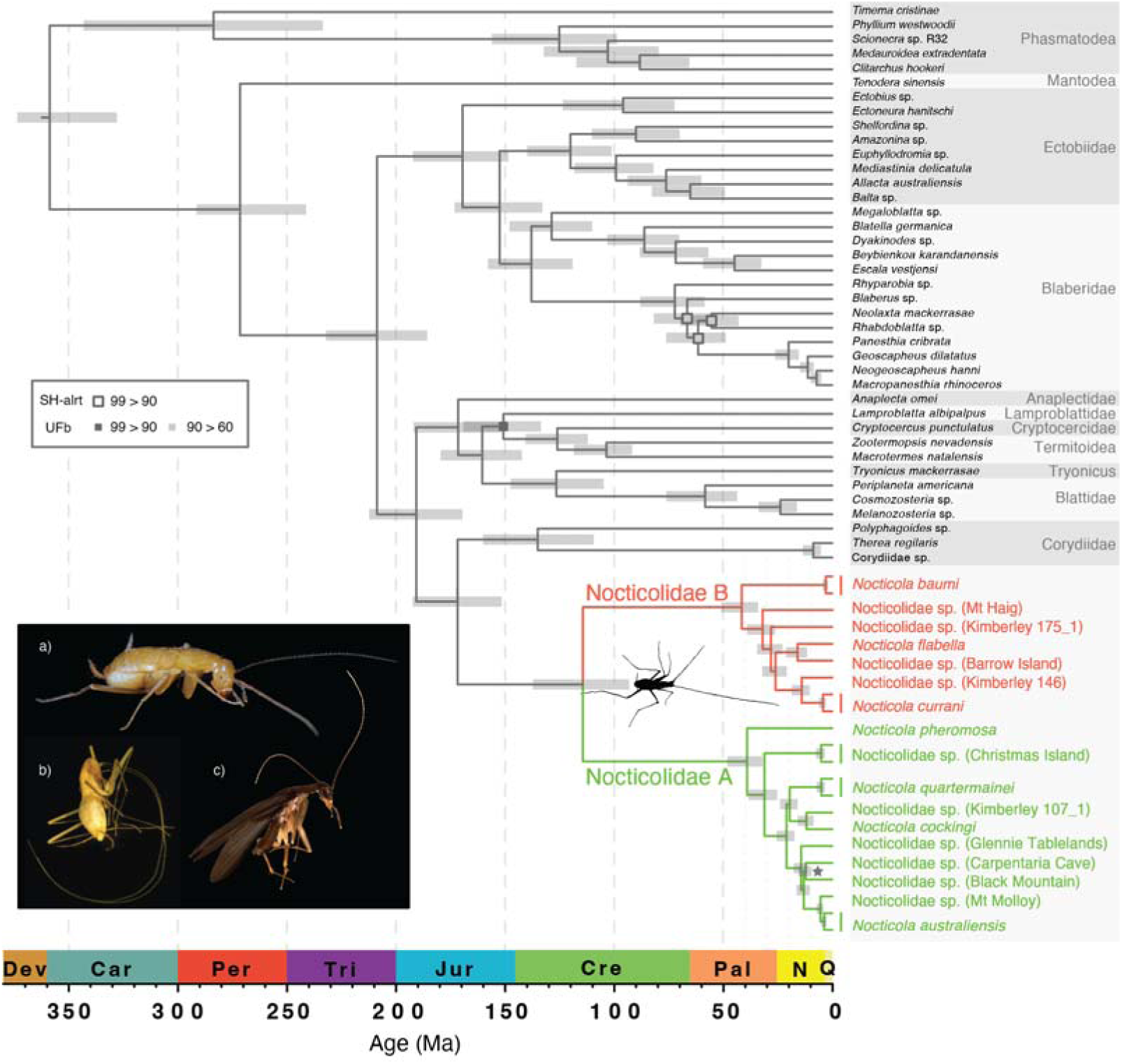
Evolutionary timescale of Blattodea, with Mantodea and Phasmatodea as outgroup taxa. Divergence times were estimated based on UCE loci, using a Bayesian phylogenomic approach with an uncorrelated relaxed clock in MCMCTree. The fixed tree topology used in this analysis was inferred using maximum-likelihood and Bayesian analyses of UCE loci. Node support from these analyses is represented by squares: outline colours indicate values of the SH-like approximate likelihood-ratio test (SH-alrt), fill colours indicate ultrafast bootstrap (UFb) values. Nodes with SH-alrt and UFb > 99 are not labelled. Posterior probabilities were 1.0 for all nodes, with the exception of the node labelled with a star, which had a different topology. Insert shows the diverse morphology present in Nocticolidae: a) female *Nocticola australiensis* (Royal Arch Cave) with highly reduced eyes (photograph by Braxton Jones); b) blind female *Nocticola baumi* with elongated appendages (photograph by Martin Bláha); and c) male *Nocticola pheromosa* with fully developed eyes and wings (photograph by Maimon Hussin).

The phylogenetic position of Nocticolidae remains uncertain because of limited taxon sampling in the family. Several phylogenetic studies have found a close relationship between Nocticolidae and the family Corydiidae, though these analyses have typically included only a small number of nocticolids (Inward et al. 2007; Legendre et al. 2008; Djernæs et al. 2012; Legendre et al. 2015; Djernæs et al. 2015; Wang et al. 2017; Bourguignon et al. 2018; Evangelista et al. 2019; Li and Huang 2020; Djernæs and Murienne 2022). There has so far been no phylogenomic analysis of all ten cockroach families, nor has there been a comprehensive phylogenetic study of nocticolids themselves. Furthermore, previous genome-scale dating analyses of Blattodea have lacked sufficient sampling of Nocticolidae to infer divergence times within the family. A single specimen was included in an analysis of transcriptomes that inferred minimal among-lineage rate variation and a Nocticolidae– Corydiidae divergence in the Jurassic (Evangelista et al. 2019). Contrastingly, a single specimen was included in an analysis of mitochondrial genomes that inferred a long nocticolid branch and an older Nocticolidae–Corydiidae divergence in the Triassic, possibly because the higher substitution rate pushed back the date estimate for this node (Bourguignon et al. 2018).

Studies based on a handful of mitochondrial and nuclear molecular markers have included up to six representatives of Nocticolidae, again highlighting increased substitution rates across the family and deep divergences (100–200 Ma) among representatives of the genus *Nocticola* (Djernæs et al. 2015; Legendre et al. 2015; Wang et al. 2017; Li and Huang 2020). Li and Huang (2020) used an autocorrelated relaxed clock to infer large increases in substitution rates within Nocticolidae and the sister lineage Latindiinae, but their divergence times remain uncertain due to sparse taxon sampling and imprecise calibrations. A population-level study of three Australian species of *Nocticola* found very large mitochondrial COI distances (17.3–25.8 %) between species, indicative of high substitution rates, deep divergences, or both (Trotter et al. 2017). There has not yet been any phylogenomic analysis of a diverse range of nocticolids, so uncertainties about their taxonomy and evolutionary history have persisted.

Here we evaluate the impact of extreme among-lineage rate variation on the inference of phylogenetic relationships and evolutionary timescales in Nocticolidae and Blattodea. We present a detailed phylogenetic analysis of all ten recognized families of cockroaches, using the largest data set so far assembled for Nocticolidae. We design the first baits for ultraconserved elements (UCEs) from cockroaches, and assemble a data set comprising UCE loci and mitochondrial genomes. Our study resolves the major branching order of all ten cockroach families, identifies two highly divergent and rapidly evolving clades within Nocticolidae, and shows that widely used relaxed-clock models have difficulty in accounting for lineage-specific accelerations in evolutionary rates.

## Materials and Methods

We generated three genetic sequence data sets for our phylogenetic analyses of nocticolids and other cockroaches. We began by using a barcoding marker, *16S*, to identify specimens for genetic sequencing, given that the majority of samples either represented undescribed species or could not be identified based on morphology. From the observed diversity in the family, we chose 22 samples for short-read metagenomic ‘genome skim’ sequencing. Using these libraries, we assembled our first data set, comprising sequences from mitochondrial protein-coding genes (mtPCG), and our second data set, comprising sequences from nuclear ultraconserved elements (UCE). We combined sequences of UCE loci, mitochondrial protein-coding genes, and mitochondrial *16S* to form a third data set with maximum taxon coverage (UCE-mtPCG-*16S*).

### Taxon Sampling and Sequencing

We sampled 131 specimens of Nocticolidae from museums and private collections. The majority of samples were from Australia (including Christmas Island), but we also included taxa from New Guinea (Indonesia) and Singapore. For non-type samples and taxa with multiple specimens, genomic DNA was extracted from whole organisms. For type specimens or individuals from unique locations for which no known species have been described, a single hind leg was used. We used PCR to amplify a 440 bp fragment of the mitochondrial gene encoding 16S rRNA (*16S*). These 115 newly generated sequences were combined with six nocticolid sequences available on GenBank, as well as outgroup sequence data from 37 non-nocticolid cockroaches and termites and five other insects (Supplementary Tables S2– S4). We aligned these sequences using MAFFT 7.475 (Katoh and Toh 2008) and manually removed poorly aligned regions. The final *16S* alignment comprised 438 bp from 163 taxa. Using this alignment, we estimated phylogenetic relationships using maximum likelihood in IQ-TREE v2 (Bui et al. 2020) to identify representative taxa for further sequencing.

For our first data set, we selected 23 representative lineages (Supplementary Table S5) for short-read metagenomic ‘genome skim’ sequencing by BGI (Shenzhen, China). Using the read libraries produced, we assembled mitochondrial genomes by mapping reads to an available nocticolid mitochondrial genome (Bourguignon et al. 2018) in Geneious 2020.0.5 (www.geneious.com). Samples with incomplete mapping were assembled *de novo* using NOVOplasty (Dierckxsens et al. 2017) and the resulting assembly was then used as a reference for read mapping. We annotated mitochondrial genes using MITOS2 (Donath et al. 2019). Mitochondrial genomes from nocticolids were combined with those from representatives of the diversity in each of the nine other cockroach families as well as outgroup taxa from related insect lineages (Supplementary Table S6; Bourguignon et al. 2018). Nucleotide sequences of each of the 13 mitochondrial protein-coding genes were aligned at the amino acid level using MUSCLE through the TranslatorX web server (Abascal et al. 2010). Using PhyloMAd (Duchêne et al. 2018, 2022), we found evidence of saturation at first and third codon sites and removed these from the sequences. The resulting data set comprised the second codon sites (3793 bp) of the 13 mitochondrial protein-coding genes (mtPCG) from 23 nocticolid taxa and 87 outgroup taxa.

To construct our second data set, we designed UCE baits using PHYLUCE 1.6.6 (Faircloth 2016). Baits were designed from whole genomes of five cockroaches and one termite, using the same parameters as Hellemans et al. (2022) for the termite-specific set. Our sampling for this approach included our nocticolid taxa, at least one representative from each of the other nine cockroach families, as well as one mantid and five phasmids as outgroup taxa. Reads were assembled using metaSPAdes 3.13 (Nurk et al. 2017), and UCEs were extracted and aligned using the PHYLUCE suite. The concatenated UCE data set comprised 1676 loci (total length 383,325 bp) from 61 samples (Supplementary Table S7). To maximize the taxon coverage in Nocticolidae and the outgroup, we constructed a third data set by combining the UCE data with mtPCG and *16S* sequences (UCE-mtPCG-*16S*). Further details of taxon selection, DNA extraction, amplification, library preparation, museum vouchers, sample locations, and accession numbers are given in the Supplementary Material.

### Phylogenetic Analyses

We inferred phylogenetic relationships using the mtPCG, UCE, and combined UCE-mtPCG-*16S* data sets using maximum likelihood in IQ-TREE. For our data sets containing UCEs (UCE and UCE+mtPCG+*16S*), we selected the partitioning scheme in ModelFinder using a greedy algorithm to merge individual UCEs and mitochondrial genes (-m TESTMERGE) (Lanfear et al. 2012; Chernomor et al. 2016). We allowed branch lengths to be proportionate among subsets of the data (-p), which has been found to result in the highest statistical support in many cases (Duchêne et al. 2020). The ModelFinder option in IQ-TREE was used to select the best-fitting substitution models according to the Bayesian information criterion (Kalyaanamoorthy et al. 2017). We also performed Bayesian phylogenetic analyses using ExaBayes v1.5 (Aberer et al. 2014) for the mtPCG and UCE data sets, checking convergence over four independent runs. In these analyses, we managed computational load by running the analysis with the GTR+G substitution model on an unpartitioned data set.

To account for gene-tree incongruence among loci, we also analysed the UCE data set using the summary-coalescent method in ASTRAL-III (Zhang et al. 2018). The gene tree for each UCE locus was estimated using IQ-TREE with substitution models selected using ModelFinder. In order to compare substitution rates across Blattodea, we calculated the mean root-to-tip distance for each cockroach family based on the maximum-likelihood trees inferred from the mtPCG and UCE data sets. We assessed the adequacy of substitution models using PhyloMAd (Duchêne et al. 2018).

We inferred the evolutionary timescale of cockroaches using Bayesian phylogenetic analyses of the mtPCG data set using BEAST v2.5 (Bouckaert et al. 2019) and the approximate likelihood method in MCMCTree (Yang 2007; dos Reis and Yang 2011). We also analysed our UCE data set in MCMCTree. In BEAST, we ran analyses using five different clock models: strict clock, uncorrelated exponential relaxed clock, uncorrelated lognormal relaxed clock, random local clock, and flexible local clock. We used the flexible local clock to model substitution rates across the tree under two independent lognormal distributions, one within Nocticolidae (including its stem branch) and one for the rest of the tree. The five clock models were compared using marginal likelihoods estimated by nested sampling (Maturana Russel et al. 2019). We used a set of fossil calibrations selected by Evangelista et al. (2019), with the addition of *Crenocticola* (Li and Huang 2020), to inform minimum bounds of uniform distributions (Supplementary Table S8). Both the mtPCG and UCE data sets were analysed in MCMCtree using the two available relaxed clock models: the independent (uncorrelated) lognormal relaxed clock and autocorrelated relaxed clock. Our UCE data set had fewer outgroup taxa, reducing the number of fossil calibrations that could be applied; we added secondary calibrations for the root of the tree and crown Dictyoptera, based on estimates from a recent phylogenomic study (Evangelista et al. 2019).

In order to test the influence of including Nocticolidae on estimates of topology and divergence times, we repeated all of our phylogenetic analyses (IQTREE, ExaBayes, BEAST, and MCMCtree) using the mtPCG and UCE data sets after removing the nocticolid taxa. For our Bayesian dating analyses, we compared all of the clock models except for the flexible local clock (for which a separate relaxed clock had previously been assigned to Nocticolidae). A detailed description of all methods can be found in the Supplementary Material.

## Results

### Phylogenetic Relationships

Phylogenetic relationships among cockroach families could not be confidently resolved using the mtPCG data set alone (Supplementary Fig. S2−S4). In contrast, our analyses of the UCE data set produced well-resolved phylogenetic trees with strong support for relationships among cockroach families (Fig. 1, Supplementary Fig. S5−S7). All of our phylogenetic analyses strongly supported the monophyly of Nocticolidae (Fig. 1, Supplementary Fig. S1-8). However, the inferred relationships between Nocticolidae and the other cockroach families differed between our mtPCG and UCE analyses. In our mtPCG analyses, the relationship between Nocticolidae and the other cockroach families varied between maximum-likelihood and Bayesian analyses and even between replicate Bayesian analyses, with Anaplectidae usually being placed as the sister group. In our analyses of UCE data, Corydiidae was consistently and confidently placed as the sister group to Nocticolidae (Fig. 1; UFb = 100, SH-aLRT = 100, ASTRAL-PP = 0.99, ExaBayes-PP = 1) and Anaplectidae was inferred to be the sister lineage to the rest of Blattoidea, with high support (UFb = 100, SH-aLRT = 100, ASTRAL-PP = 0.99, ExaBayes-PP = 1). After removing nocticolid taxa, we found that the phylogenetic relationships among cockroach families could still not be confidently resolved using the mtPCG data set (Supplementary Fig, S13-15). However, the topologies inferred using the UCE data set were congruent to those from our analyses that included Nocticolidae (Supplementary Fig, S16-17). Our assessment of substitution model adequacy found that the full mtPCG data set had a high risk of biased inferences according to the consistency index. This statistic reported a low risk of biased inferences after removing nocticolid taxa. Unfortunately our UCE data set contained too much missing data to assess substitution model adequacy.

Within Nocticolidae, our analyses found uniformly strong support for the mutual monophyly of two divergent clades. We refer to these clades as Nocticolidae A and B. The inferred relationships within each of these two clades varied slightly between mtPCG and UCE analyses. Here we focus on the results of our analyses of the data sets that included the UCE loci, which yielded the highest phylogenetic support. Within Nocticolidae, consistent relationships were found between our analyses of the UCE data set (Fig. 1) and the UCE+mtPCG+*16S* data set (Fig. 2). A nocticolid sequence obtained from GenBank (“*Nocticola* sp. Unknown”; Legendre et al. 2015), for which collection location details are not available, was found to be the sister lineage to all other nocticolids examined.

**FIGURE 2.**
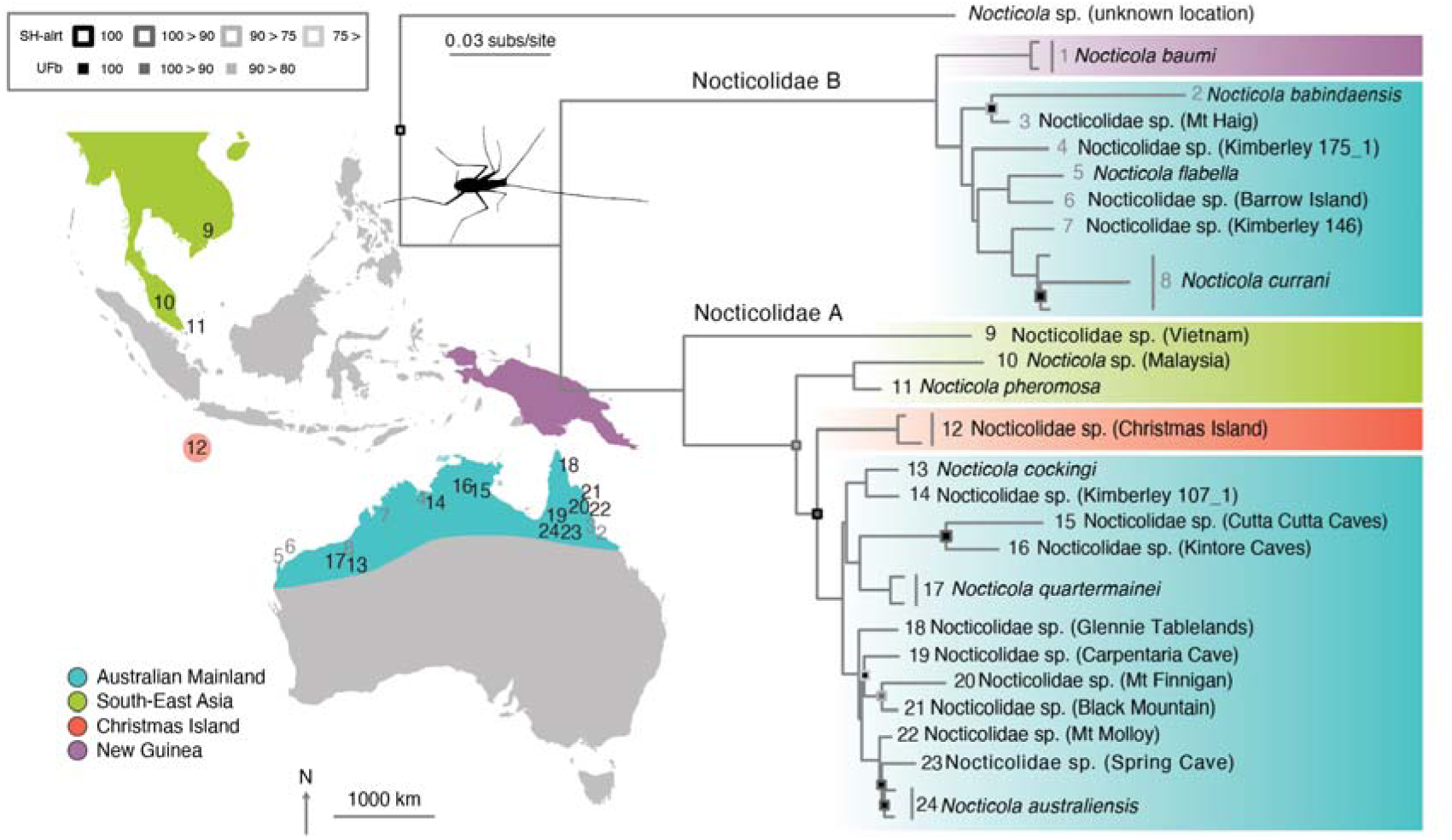
Phylogeny of Nocticolidae. Maximum-likelihood tree estimated using concatenated UCE loci, second codon sites of 13 mitochondrial protein-coding genes, and mitochondrial *16S*. There are two major clades in Nocticolidae, A (numbers in black) and B (numbers in grey). Node support is represented by squares: outline colours indicate values of the SH-like approximate likelihood-ratio test (SH-alrt), fill colours indicate ultrafast bootstrap (UFb) values. Nodes with SH-alrt and UFb = 100 are not labelled. Colours represent the geographical location of taxa from the Australian mainland (blue), South-East Asia (green), Christmas Island (orange), and New Guinea (purple).

Nocticolidae A contains approximately half of the Australian mainland taxa, nested within three lineages containing taxa from Vietnam, Singapore and Malaysia, and Christmas Island, respectively (Fig. 2). Among the Australian taxa in Nocticolidae A, Queensland taxa formed the sister clade to a group comprising taxa from Western Australia and the Northern Territory. The second clade, Nocticolidae B, comprised a sister grouping between the remaining Australian taxa and one taxon from New Guinea (*N. baumi*). The Queensland taxa, *N. babindaensis* and *N*. sp. (Mt Haig), again formed the sister clade to taxa from Western Australia and the Northern Territory.

### Evolutionary Timescales and Substitution Rates

Our phylogenetic analyses suggest that an acceleration in mitochondrial substitution rates occurred early in the evolution of Nocticolidae. In our maximum-likelihood analyses of the mtPCG and UCE data sets, root-to-tip distances for nocticolid clades were considerably greater than those of other cockroach families (Fig. 3). For the mtPCG data set, Nocticolidae A and B had root-to-tip distances that were 3–5 times those of representatives of other cockroach families. In comparison, the UCE sequences showed root-to-tip distances for nocticolids that were ∼1.5 times those of other cockroach families.

**FIGURE 3.**
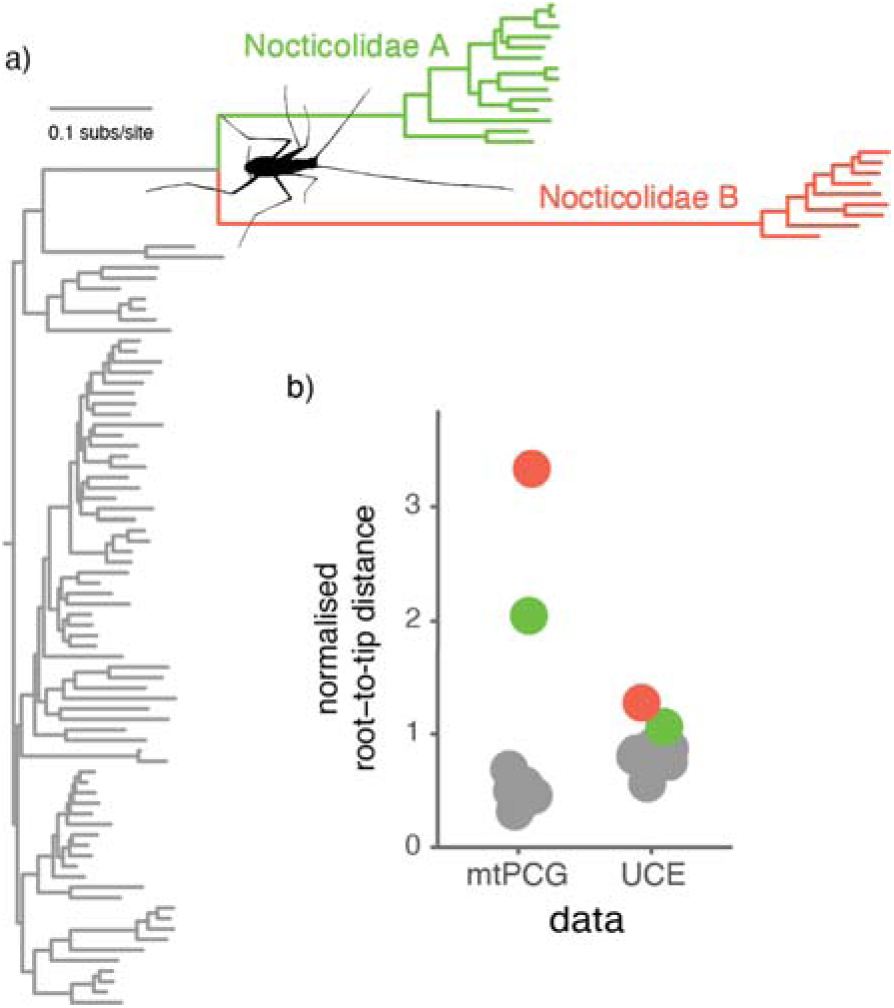
Accelerated mitochondrial substitution rates in Nocticolidae compared with other families in Blattodea (cockroaches). (a) Maximum-likelihood tree of Blattodea inferred using the second codon sites of the 13 mitochondrial protein-coding genes. (b) Phylogenetic root-to-tip distances for the concatenated mitochondrial protein-coding genes and the concatenated UCE loci. Root-to-tip distances were calculated for each of the major cockroach families, with Nocticolidae split into its two major clades, A and B. Root-to-tip distances were normalized by dividing the mean for each family by the mean root-to-tip distance across all taxa.

In the Bayesian phylogenetic analysis of the mtPCG data set using BEAST, the flexible local clock was decisively preferred over other clock models (minimum log Bayes factor = 30.8, SD = 6.0, Supplementary Table S9). The flexible local clock inferred a substantially higher rate in Nocticolidae (posterior mean 3.7×10^-3^ substitutions/site/Myr; 95% CI 2.6–6.3×10^-3^) than across the rest of the tree (posterior mean 8.0×10^-4^ substitutions/site/Myr; 95% CI 6.4–9.4×10^-4^) (Fig. 4). Our analyses using the uncorrelated exponential and lognormal relaxed-clock models inferred very high rates along the stem branches leading to Nocticolidae and to Nocticolidae A and B. Within each of these clades, however, the inferred rates were similar to the background rate across the tree (Fig. 4).

**FIGURE 4.**
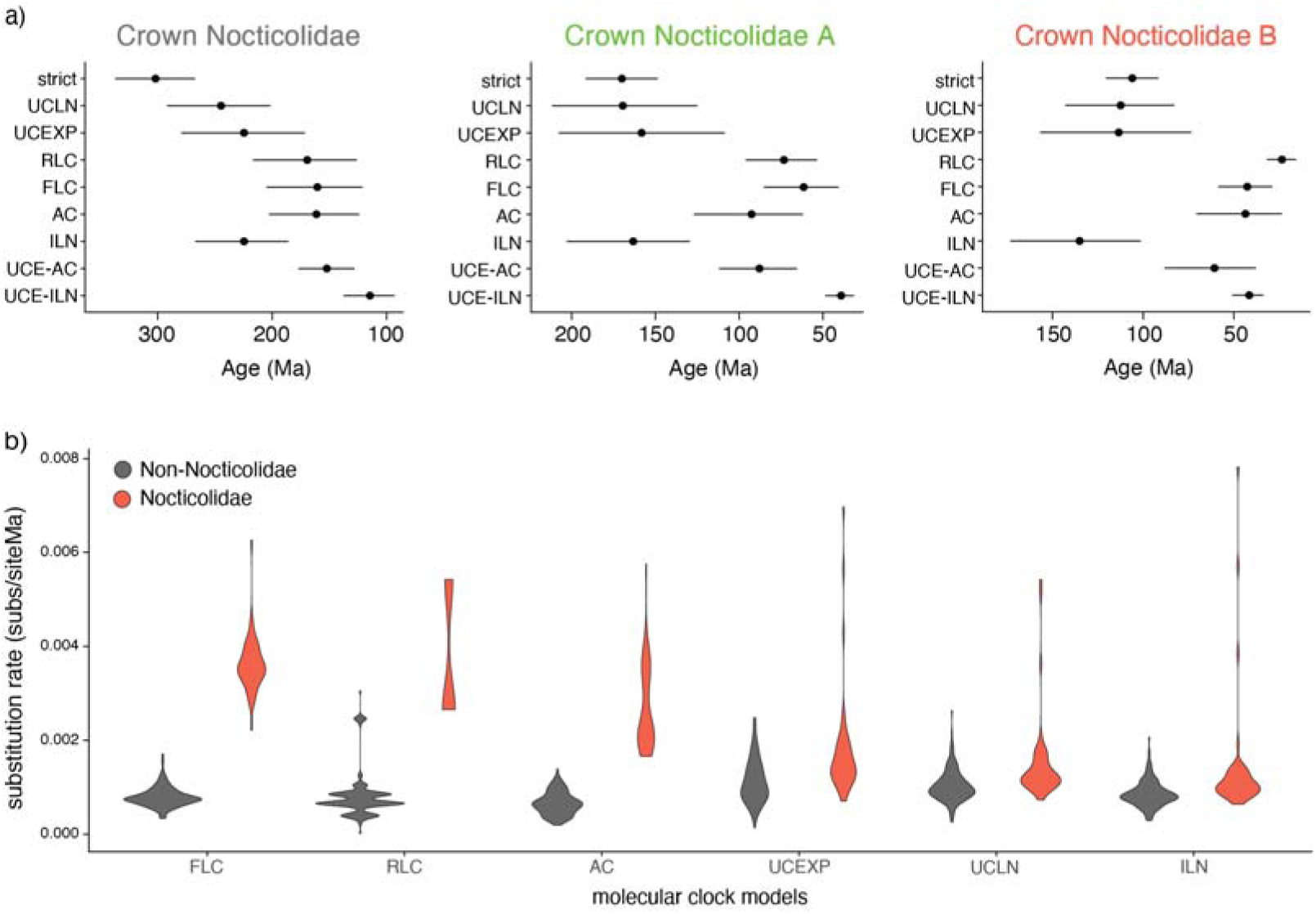
Estimates of divergence times and substitution rates using a range of molecular-clock models. (a) Dates for crown Nocticolidae, crown Nocticolidae A, and crown Nocticolidae B estimated from mitochondrial protein-coding genes using a strict clock (strict), uncorrelated lognormal relaxed clock (UCLN), uncorrelated exponential relaxed clock (UCEXP), random local clock (RLC), flexible local clock (FLC), autocorrelated relaxed clock (AC), and independent lognormal relaxed clock (ILN). Date estimates for the same nodes using our UCE data set are also presented for comparison using an autocorrelated relaxed clock (UCE-AC) and independent lognormal relaxed clock (UCE-ILN). Circles indicate posterior means and bars indicate 95% credibility intervals. b) Estimates of mitochondrial substitution rates in Nocticolidae compared with other families in Blattodea (cockroaches). Violin plots represent the mean substitution rate (substitutions per site per Myr) for all terminal and internal branches within Nocticolidae (including stem) and outside Nocticolidae (non-Nocticolidae). Rates were estimated using concatenated mitochondrial protein-coding genes.

Our Bayesian analyses inferred deeper divergence times when using mtPCG data than UCE data. Among our analyses of the mtPCG data, the flexible local clock produced the youngest estimates of divergence times, particularly for crown Nocticolidae (Fig. 4). These estimates were the most similar to those inferred using the UCE data set. Our analyses using the random local clock estimated the second most similar divergence times to those inferred using the UCE data set, and also identified a clear distinction between substitution rates in Nocticolidae and across the rest of the tree. The strict clock estimated the oldest dates for nodes within Nocticolidae, presumably to accommodate an excess of substitutions in nocticolid lineages.

In the molecular dating analysis of the mtPCG data set using MCMCtree, the autocorrelated relaxed clock gave very similar estimates of substitution rates and divergence times to those inferred using the flexible local clock in BEAST (Fig. 4). The independent lognormal relaxed clock in MCMCtree gave similar estimates of substitution rates and divergence times to those inferred using the uncorrelated relaxed-clock models in BEAST (Fig. 4). For our UCE data set, the independent lognormal and autocorrelated relaxed clocks yielded different estimates of substitution rates and divergence times. The independent lognormal relaxed clock estimated slightly higher substitution rates in Nocticolidae compared with the rest of the tree, aligning with the results from our investigation of root-to-tip distances (Supplementary Fig. S10). The autocorrelated relaxed clock estimated that the highest substitution rates occurred along the branches leading to the phasmid and mantid outgroups, the early branches among the major cockroach families including the stem of Noctioclidae, and the stem branches leading to the two nocticolid clades (Supplementary Fig. S10). Estimated substitution rates within each of the families were generally lower than those along the respective stem branches, with branches within each of the major Nocticolidae clades having the lowest rates in the tree (Supplementary Fig. S10). Consequently, for the UCE data set the autocorrelated relaxed clock yielded older estimates of divergence times within Nocticolidae than did the independent lognormal relaxed clock (Fig. 4).

Our dating analyses after removing nocticolid taxa support our contention that high rates in Nocticolidae are driving older estimates of divergence times using some clock models. For the mtPCG data set, the dates of crown Blattodea and Dictyoptera estimated using the lognormal relaxed clock, exponential relaxed clock, and strict clock were reduced by the removal of nocticolid taxa, and similar to those inferred using the flexible local clock when including Nocticolidae (Supplementary Figures S15,20).

We consider our estimate of evolutionary timescales based on UCE data and the independent lognormal relaxed clock to be the most reliable for various reasons. First, the inferences are based on a large genome-scale data set with representatives from all 10 cockroach families. Second, the UCE data have low among-lineage rate variation, especially compared with the mtPCG data set. Third, the independent lognormal relaxed clock appears to be the most appropriate because the distribution of branch rates estimated by the autocorrelated relaxed clock is inconsistent with the root-to-tip distances; branches within Nocticolidae were inferred to have a lower substitution rate than the rest of the tree, despite its taxa having greater root-to-tip distances. We believe that this led to overestimation of divergence times in Nocticolidae using this model.

Based on our analysis of UCE data with the independent lognormal relaxed clock, the most recent common ancestor of cockroaches and mantids is dated at 272 Ma (95% CI 241– 292 Ma), while that of cockroaches is dated at 209 Ma (95% CI 186–233 Ma). The age of crown Nocticolidae was inferred to be 115 Ma (95% CI 93–138 Ma), with the family diverging from Corydiidae at 172 Ma (95% CI 152–193 Ma). The crown age of Nocticolidae A was estimated at 39 Ma (95% CI 32–48 Ma), with the Australian lineages diverging from each other at 21 Ma (95% CI 17–26 Ma). Nocticolidae B was inferred to have a crown age of 42 Ma (95% CI 34–51 Ma), with the Australian lineages diverging from each other at 32 Ma (95% CI 26–39 Ma).

## Discussion

Our analyses have highlighted a substantial elevation of evolutionary rates in the mitochondrial genomes of nocticolids, particularly along the basal branches of the clade, and that a similar but smaller increase in rates has occurred in the nuclear genome. These results are partly consistent with previous findings of long nocticolid branches in mitochondrial trees (Lo et al. 2007; Legendre et al. 2008, 2015; Djernæs et al. 2015; Wang et al. 2017; Bourguignon et al. 2018; Li and Huang 2020) but not in trees inferred from nuclear protein-coding genes from transcriptomic data (Evangelista et al. 2019). We have also found that some of the widely used relaxed-clock models are unable to accommodate the large disparities in mitochondrial substitution rates across branches. This rate heterogeneity is likely to be at least partly responsible for the differences in evolutionary relationships and divergence times between the phylogenetic trees inferred from mitochondrial genomes and nuclear ultraconserved elements.

### Molecular Dating When Rates Vary

Mitochondrial substitution rates differed by a factor of more than 3 between nocticolids and other cockroaches, representing a marked elevation of rates in a single family compared with its relatives. This disparity in rates was apparent in the root-to-tip distances of the phylograms and was identified in the branch rates inferred under the flexible local clock and random local clock in BEAST, as well as the autocorrelated relaxed clock in MCMCtree. The ability of these models to infer such highly distinct rates across the tree aligns with previous applications of these models, and are expected to provide a better fit under conditons of complex rate variation among lineages (Fourment and Darling 2018; Drummond and Suchard 2010). Among the clock models compared in BEAST, the flexible local clock had the highest marginal likelihood and yielded mitochondrial estimates of divergence times that were most similar to those inferred from UCE data. Although the random local clock was also able to identify an elevation of mitochondrial rates in nocticolids, it had a much lower marginal likelihood that indicated poorer fit. This model produced a date estimate for crown Nocticolidae that was similar to that inferred using the flexible local clock, but appeared to have difficulty in resolving the location of an early shift in rates within the family.

The uncorrelated relaxed clocks in BEAST and MCMCtree were unable to reconstruct the disparity in substitution rates between nocticolids and other cockroaches. Notably, the three relaxed-clock models inferred dates for nocticolids that were broadly similar to those obtained under a strict clock, but widely divergent from those inferred from the UCE data. Furthermore, the inferred topologies varied across BEAST analyses, illustrating that the impacts of clock-model misspecification extend beyond estimates of substitution rates and divergence times. These results call into question the reliability of the divergence times inferred using these widely used relaxed-clock models under conditions of extreme rate variation. Our analysis of UCEs with an independent lognormal relaxed clock produced younger estimates of divergence times within Nocticolidae compared with our other analyses and those inferred previously (Djernæs et al. 2015; Wang et al. 2017; Li and Huang 2020), which were likely to have been pushed back to accommodate the greater numbers of substitutions on nocticolid branches. However, we also used slightly different calibration methods between our analyses of UCE data and mtPCG data.

Although we focus on Nocticolidae as a case study, our results have broad implications for estimating evolutionary timescales for groups with high among-lineage rate variation. We find that the flexible local clock, random local clock, and autocorrelated clock models produce the most concordant estimates of substitution rates and divergence times when analysing data containing high among-lineage rate variation. The unifying feature of the local clock models is their ability to allow a distinct rate, or distribution of rates, in Nocticolidae. The reasons for the performance of the autocorrelated clock model are less clear, but it could be that the model does not substantially penalize a single jump in rates at the base of Nocticolidae. Nevertheless, the different estimates of rate variation within Nocticolidae highlight the problems associated with each model. For example, the two replicate analyses using a random local clock had identified a separate local clock for each clade in Nocticolidae, but had grouped the rate of stem Nocticolidae with different clades of Nocticolidae (A or B). This led to age estimates for crown Nocticolidae that differed between the two analyses. Although we attempted to account for this uncertainty by combining the posterior samples, poor Markov chain convergence appears to be a common issue for this model when applied to large data sets (Dornburg et al. 2012, Drummond and Suchard 2010).

Applying an independent unimodal parametric distribution of branch rates to Nocticolidae through the flexible local clock appeared to model rate variation between Nocticolidae and the rest of the tree effectively. However, this restricted separation of rates between the two clades within Nocticolidae, instead accounting for rate variation in the family by estimating higher substitution rates on a few branches. This resulted in the highest substitution rate on the stem branch of Nocticolidae B while comparable rates were present within Nocticolidae A and B and on the stem branch of Nocticolidae A, likely leading to an overestimation of divergence times in Nocticolidae B.

The random local clock is the only model that maintained a high substitution rate throughout Nocticolidae B, which appears to be more biologically plausible, leading to the youngest age estimate for crown Nocticolidae B. Even our analyses of UCEs, which had only slightly greater root-to-tip distances in Nocticolidae compared with the rest of the tree, inferred a slightly increased rate in stem Nocticolidae and stem Nocticolidae B compared with the rest of the group, potentially leading to slight overestimation of divergence dates in Nocticolidae B. Perplexingly, analysis of UCE data using the autocorrelated clock model estimated substitution rates in Nocticolidae to be lower than across the rest of the tree, highlighting that this model can misbehave when there is minimal among-lineage rate variation. Our results suggest that molecular dating analyses should be carried out using data sets with minimal rate variation among lineages, where posible. This could include data sets such as the UCEs analysed here or, when dealing with whole-genome data, this could involve filtering loci for clocklikeness (Jarvis et al. 2014; Smith et al. 2018). Selecting loci with lower among-lineage rate variation has additional benefits for phylogenetic resolution (Vankan et al. 2021).

### Phylogenetic Inference When Rates Vary

Although not the focus of this study, we briefly address the potential effects of extreme among-lineage rate variation on our phylogenetic inference. Our analysis of UCE data found strong support for Corydiidae as the sister group to Nocticolidae, confirming the results of previous analyses. The relationships among the major cockroach families are also congruent to those inferred using transcriptomes (Evangelista et al. 2019), with the addition of the family Anaplectidae as the sister group to the rest of Blattoidea. This placement conflicts with the results of previous studies that supported a close relationship between Anaplectidae and Lamproblattidae based on mitochondrial genomes (Bourguignon et al. 2018), or that placed Lamproblattidae as the sister group to the rest of Blattoidea with Anaplectidae as the sister group to Cryptocercidae+Isoptera and Tryonicidae based on mitochondrial and nuclear markers (Djernæs and Murienne 2022).

The results of our UCE analysis also conflict with the evolutionary relationships inferred from our mtPCG data set, which united Nocticolidae with Anaplectidae. The instability of the position of Nocticolidae in our analyses of mitochondrial genomes could be due to the long stem branches leading to the two clades of the family. In our analysis, there was also a long stem branch leading to Anaplectidae because of minimal taxon sampling in the family. Consequently, long-branch attraction could have occurred in the maximum-likelihood analysis, even though this phenomenon is less pronounced when using model-based phylogenetic methods (Swofford et al. 2001; Kolaczkowski and Thornton 2009; Kück et al. 2012). Furthermore, our tests of model adequacy suggested a high risk of biased inferences according to the consistency index, suggesting that among-lineage rate variation could be responsible for the incongruence between our data sets. These results highlight that extreme among-lineage rate variation can still have substantial impacts on model-based methods of statistical phylogenetic inference.

### Causes of Increased Substitution Rates in Nocticolidae

The causes of the increased mitochondrial and nuclear rates in Nocticolidae are yet to be determined. Previous studies have suggested that the rate acceleration might be the result of small population sizes and repeated bottlenecks associated with the colonization of caves, or relaxed selection associated with the loss or reduction of characters no longer required in subterranean environments (Bourguignon et al. 2018). Increased mitochondrial substitution rates have been associated with the loss of flight in a number of insect lineages (Mitterboeck and Adamowicz 2013). However, our results suggest that an increase in substitution rate occurred early in the evolution of Nocticolidae when the ancestor had not yet become restricted to the subterranean habitat, and presumably still possessed fully developed wings. Similar patterns have been seen using a smaller number of nocticolids (Legendre et al. 2015; Li and Huang 2020). Previous studies have placed the subfamily Latindiinae as the sister lineage to Nocticolidae (Legendre et al. 2015; Wang et al. 2017; Li and Huang 2020; Liu et al. 2023), which also exhibits increased substitution rates despite not being associated with subterranean habitats, suggesting that habitat type is unlikely to be the sole driver of evolutionary rate acceleration.

One common factor across all studied lineages of Nocticolidae, including epigean and subterranean taxa, is their reduced body size compared with other cockroaches. Small body size is associated with higher substitution rates in a number of animals, presumably because of its covariation with shorter generations, higher metabolic rate, and increased fecundity, none of which has been studied in Nocticolidae (Bromham 2009; Thomas et al. 2010). In cockroaches, there is a positive relationship between body size and clutch size, although this might be offset by increased frequency of clutches in smaller-bodied species (Djernæs et al. 2020). Other groups of cockroaches with small body sizes have also shown greater phylogenetic root-to-tip distances, including the subfamily Latindiinae, and a distantly related genus of minute myrmecophilous cockroaches, *Attaphila* (Djernæs et al. 2020). The effects of life-history traits in elevating substitution rates are more likely to be seen in mitochondrial genomes, which have smaller effective population sizes than nuclear genomes.

We found evidence of increased substitution rates for nocticolids in the UCE data set, although less pronounced than the increases found in mitochondrial genomes. The UCE loci are likely to be under strong selective constraints because many are found in exonic regions in insects (Zhang et al. 2019; Hellemans et al. 2022)(Zhang et al. 2019; Hellemans et al. 2022). Our finding of an increased rate in the nuclear DNA of nocticolids stands in contrast with the results of a previous analysis of a transcriptome data set, which found broadly similar root-to-tip distances among nocticolids and other taxa (Evangelista et al. 2019). However, the transcriptome data set comprised the second codon sites from ∼2000 protein-coding genes conserved across a diverse group of insects, which are likely to be under strong purifying selection.

### Evolutionary History of Nocticolidae

Our results suggest that Nocticolidae is an old family with a crown age of ∼115 Ma and most recent common ancestor with Corydiidae in the Jurassic. This aligns with results from recent molecular dating studies (Evangelista et al. 2019; Li and Huang 2020), and the existence of ∼100 Myr old nocticolid fossils from Asia (Li and Huang 2020; Sendi et al. 2020). The crown age of the family might be even older than inferred in this study, given that we did not include nocticolid representatives from Africa and Madagascar.

The estimated timing of divergence between Nocticolidae A and B suggests that these two lineages separated after the breakup of Pangaea (∼200–150 Ma) (Seton et al. 2012). An Asian origin for Nocticolidae A is suggested by the placement of Asian taxa and the nested position of Australian mainland and Christmas Island taxa. Our date estimates suggest that nocticolids arrived on the Australian mainland some time between 31 and 21 Ma, which aligns with the collision of the Australian and Asian tectonic plates ∼25 Ma. Around this time, there was extensive floral and faunal exchange between Australia and South-East Asia (Maekawa et al. 2003; Crayn et al. 2015). The estimated crown age of 42 Ma for Nocticolidae B suggests its presence on the ancient Australia/New Guinea landmass prior to its collision with the Asian plate to the north. However, because we believe our divergence dates in Nocticolidae B could be slight overestimates, we cannot rule out the possibility of dispersal into Australia around the same time as Nocticolidae A, especially considering the close relationship inferred between *N. babindaensis* and a specimen from China (Wang et al. 2017). Furthermore, any biogeographic hypotheses should be considered tentative due to the absence of African, Madagascan, and other Asian lineages in our data sets. Further analyses involving additional taxa will be needed to test the biogeographic origins of Nocticolidae B and of Nocticolidae.

Australia was largely covered in rainforest ∼25–100 Ma (Byrne et al. 2011), which would have favoured the movement of ancestral epigean nocticolids across the continent. Periods of aridity from the Miocene onwards led to rainforest habitats receding to a few refugia along the east coast of Australia (Byrne et al. 2011). These conditions might have forced various ancestral nocticolid lineages to enter caves or other subterranean habitats, or to develop an inquiline relationship with burrowing insects.

### Nocticolid Systematics and Taxonomy

Our study is the first to include genetic data for a diverse range of Nocticolidae and highlights the need for taxonomic revision in the family (Roth 1988). A recent morphological analysis was unable to resolve relationships amongst the genera of Nocticolidae and Corydiidae (Li and Huang 2020). By combining morphological data for all individuals of a genus, the authors assumed the monophyly of each, despite widespread taxonomical uncertainty. As with most troglomorphic taxa, regressive evolution is likely to have occurred in parallel across Nocticolidae, owing to multiple independent transitions into caves. Therefore, it is likely that the current taxonomy of the group, which is based on morphology, does not accurately reflect true evolutionary relationships. Our analysis of UCE data yielded an estimate of ∼115 Ma for the crown age of *Nocticola*, making this genus very old and potentially containing at least some of the other nocticolid genera. Our samples from Christmas Island are likely to represent *Metanocticola christmasensis*, which would confirm the non-monophyly of *Nocticola*, although we only sampled nymphs which are difficult to identify (Roth 1999) (full discussion in Supplementary Material). Further genetic sampling of other nocticolid genera is required to confirm that *Nocticola* is indeed polyphyletic.

Members of Nocticolidae A and B are likely to represent two distinct genera. Representatives of the genus *Nocticola* have previously been split into two groups: the *uenoi*-species group, in which the male terga are specialized; and the *simoni*-species group, in which the male’s terga are unspecialized (Roth 1988). This grouping does not align with the phylogenetic relationships inferred here. Instead, tergal glands appear to have been derived independently on multiple occasions, potentially during subterranean adaptation, as they are not present in any epigean or inquiline taxa. Tergal specializations vary substantially in *Nocticola*, with some taxa (e.g., *N. australiensis* and *N*. sp. (Carpentaria Cave)) possessing a simple sclerotized indent on the fourth tergum, while others have complex structures involving multiple tergal segments (e.g., *N. currani*, *N. ueoni*, and *N*. sp. (Kimberley 175_1)).

There are two morphological characters that align with the relationships inferred using molecular data and are good candidate synapomorphies. One potential synapomorphy of members of Nocticolidae A constitutes a group of large sclerotized spines on the ventral surface of segments 4–6 of the female cerci (Roth 1988; Trotter et al. 2017). This has been observed in all available females of Nocticolidae A (Supplementary Table S1), including *Nocticola* sp. (Malaysia) and our Christmas Island samples, which suggests that the trait could be ancestral in this clade (Mari Fujita, personal communication). Importantly, this character was not observed in any of the females in Nocticolidae B. This character could provide some guidance for the placement of species without genetic data (see Supplementary Table S1). Unfortunately, no females have been sampled from most of the other nocticolid genera, but females in *Speleoblatta* and *Alluaudellina* do not have this morphology (Chopard 1932; Vidlička et al. 2003, 2017). *Alluaudellina* is likely to be a synonym of *Nocticola* (Chopard 1932; Roth 1988) and the lack of female cerci spinals suggests that African nocticolids are not closely related to Nocticolidae A, although genetic analysis will be required to confirm this.

Previous studies have found striking differences in male genitalia between closely related species in *Nocticola* (Trotter et al. 2017). One genital character that aligns with our molecular results is the shape of L3d (left phallomere 3 dorsal). In three representatives of Nocticolidae B (*N.* sp. (Kimberley 175_1), *N. currani*, and *N. flabella*), L3d is long, curved, and spear-like (Roth 1991; Trotter et al. 2017). Contrastingly, in five representatives of Nocticolidae A (*N. cockingi*, *N. quartermainei*, *N*. sp. (Kimberley 107_1), *N. australiensis*, and *N*. sp. (Glennie Tablelands)), L3d is rounder, complex, and shovel-like (Trotter et al. 2017). The spear-like L3d present in the African species *N. clavata*, *N. scytala*, *N. wliensis*, and *A. cavernicola* (Chopard 1932; Andersen and Kjaerandsen 1995) appears to be similar to the morphology seen in Nocticolidae B, although modern, high quality genitalia drawings (eg. Lucañas and Lit 2016; Trotter et al. 2017; Lucañas and Maosheng 2023) are required to confirm this.

It is also possible that unsampled nocticolids represent sister lineages to the two studied here, potentially with a close relationship to the undescribed sample from an unknown location which we found to be the sister lineage to the rest of Nocticolidae. Unfortunately, the type species for the genus, *Nocticola simoni*, cannot be grouped into either clade because of its basic description (Bolivar 1892). Therefore, re-examination and generation of genomic data for the type taxon is required to identify whether Nocticolidae A or B should be redescribed as *Nocticola*, and which of these should be described as a new genus.

## Conclusions

In performing the largest phylogenomic and molecular dating analysis of nocticolid cockroaches, we have uncovered an extraordinary level of genetic diversity in the family. Nocticolids have experienced remarkably high substitution rates compared with other cockroaches, especially in mitochondrial protein-coding genes. This extreme lineage-specific rate acceleration is likely to have misled phylogenetic inference using maximum-likelihood and Bayesian methods. Our phylogenetic analysis of UCE loci has resolved the deep relationships among the major cockroach families, in uniting Corydiidae with Nocticolidae and placing Anaplectidae as the sister lineage to the rest of Blattoidea. Within Nocticolidae, we identify two highly divergent clades that separated ∼115 Ma despite both containing representatives of the cryptic genus *Nocticola*, suggesting that the genus requires taxonomic attention.

Our results suggest that some of the widely used relaxed-clock models are unable to account for large disparities in substitution rates among lineages, with potentially negative impacts on estimates of divergence times. In this regard, localized accelerations of rates can be taken into account using models such as the flexible local clock. This model has not been widely used, possibly because it requires that the number and phylogenetic placements of local relaxed clocks be chosen a priori. However, unusual patterns of among-lineage rate heterogeneity call for the use of more complex clock models. Modelling the variation in molecular rates more accurately will allow more reliable reconstructions of evolutionary timescales and phylogenetic relationships.

## Supplementary Material

Data available from the Dryad Digital Repository: http://dx.doi.org/10.5061/dryad.[NNNN]

## Author Contributions

TGLK, JW, SYWH, and NL conceived and designed the study. Personal collections were provided by JW and sample collection was undertaken by TGLK, JW, and NL. NJT and JMM provided museum and environmental survey samples. SH and TB designed and generated UCE data. SYWH provided input into phylogenetic and molecular dating analyses. TGLK completed the wet lab work, analysed the molecular data, performed data analyses, and drafted the manuscript with assistance from SYWH and NL. All authors approved the manuscript prior to submission.

## Acknowledgements

We thank Bruce Gray for help with collecting *Nocticola* specimens, Mari Fujita for correspondence regarding the morphology of *Nocticola* sp. (Malaysia), Maosheng Foo (LKCNHMS), Jiří Patoka, and Martin Bláha for providing specimens, and Kyle Ewart and Yi-Kai Tea for guidance on data analyses. We also thank the late Fred Stone for his lifelong contribution to furthering our understanding of subterranean fauna. The authors acknowledge the Sydney Informatics Hub and the use of the University of Sydney’s high-performance computing cluster, Artemis. We thank an anonymous reviewer and the Editors for valuable input on an earlier version of this manuscript.

