## Supplementary Figures for "Dating in the Dark: Elevated Substitution Rates in Cave Cockroaches (Blattodea: Nocticolidae) Have Negative Impacts on Molecular Date Estimates"

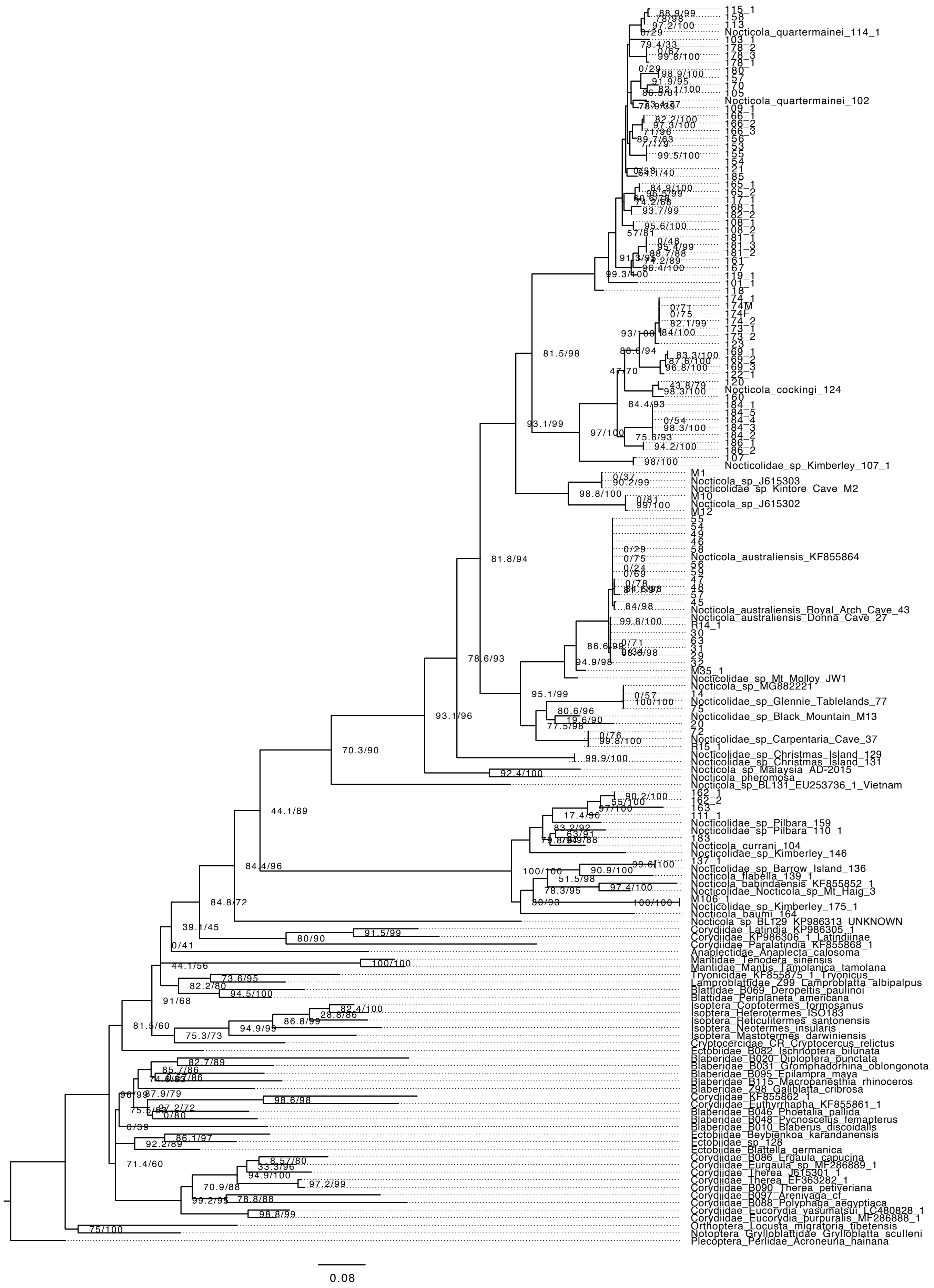

Supplementary Figure S1. Maximum-likelihood tree of Blattodea inferred using a 440 bp fragment of the mitochondrial gene encoding 16S rRNA . Phylogenetic relationships of Nocticolidae estimated using maximum likelihood based on . Node values represent SH-like approximate likelihood-ratio test (SH-alrt) and ultrafast bootstrap (UFb) values. Scale bar indicates 0.08 substitutions per site.

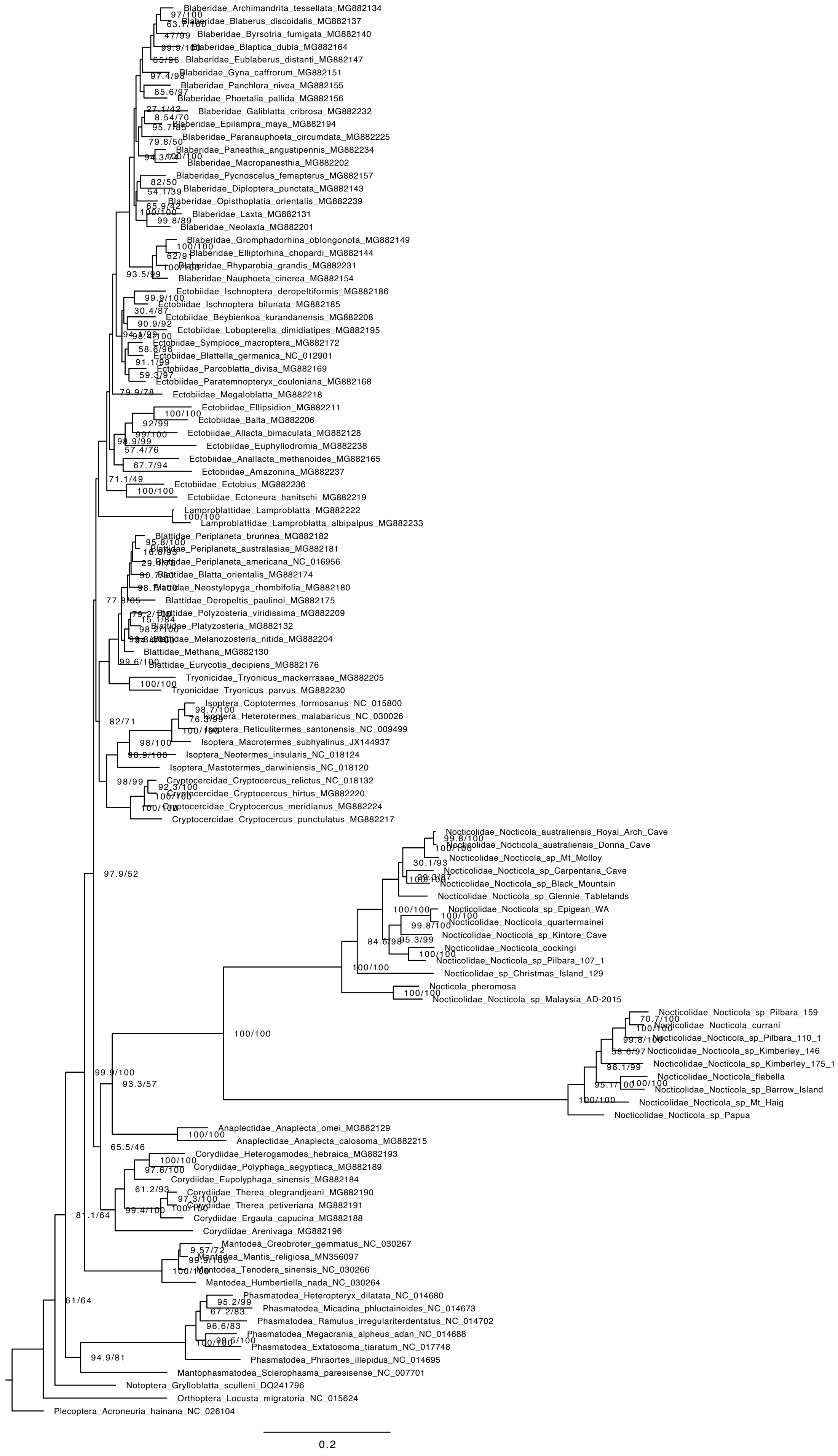

Supplementary Figure S2. Maximum-likelihood tree of Blattodea inferred using the second codon sites of the 13 mitochondrial protein-coding genes. Scale bar indicates 0.2 substitutions per site. Node values represent SH-like approximate likelihood-ratio test (SH-alrt) and ultrafast bootstrap (UFb) values.

a)

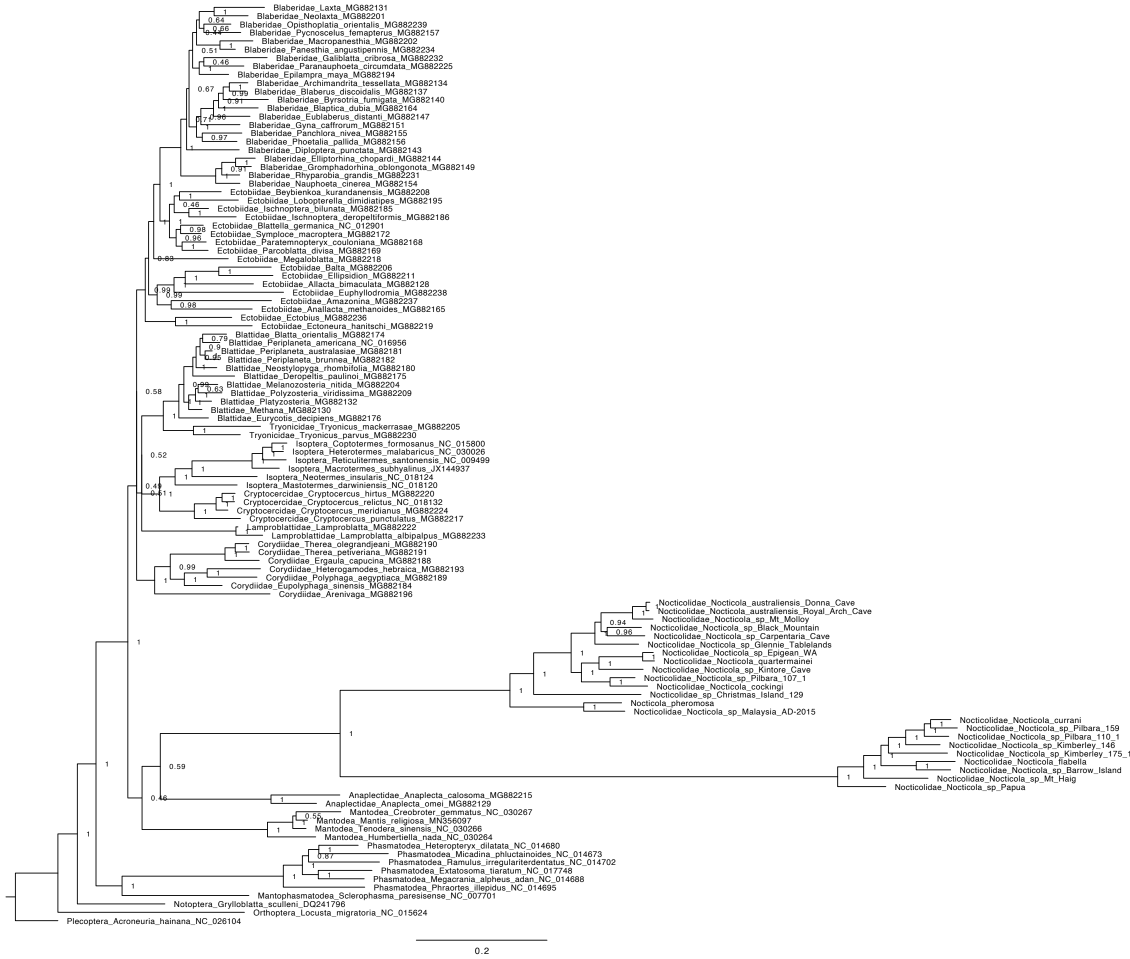


b)

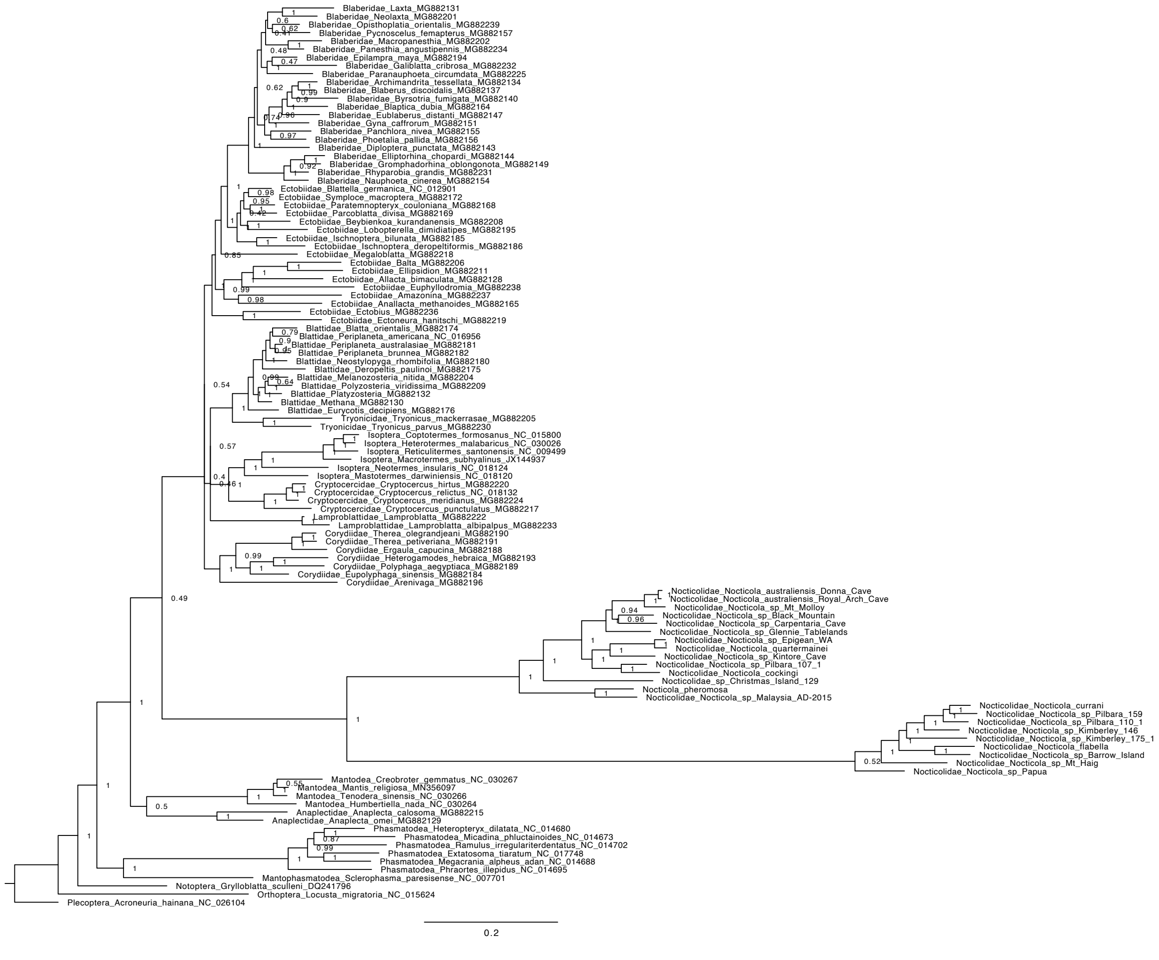


Supplementary Figure S3. Bayesian inference tree of Blattodea inferred using the second codon sites of the 13 mitochondrial protein-coding genes. Scale bar indicates 0.2 substitutions per site. Node values represent posterior probabilities. a-d indicate the differing topologies inferred from the four parallel mcmc simulations.

c)

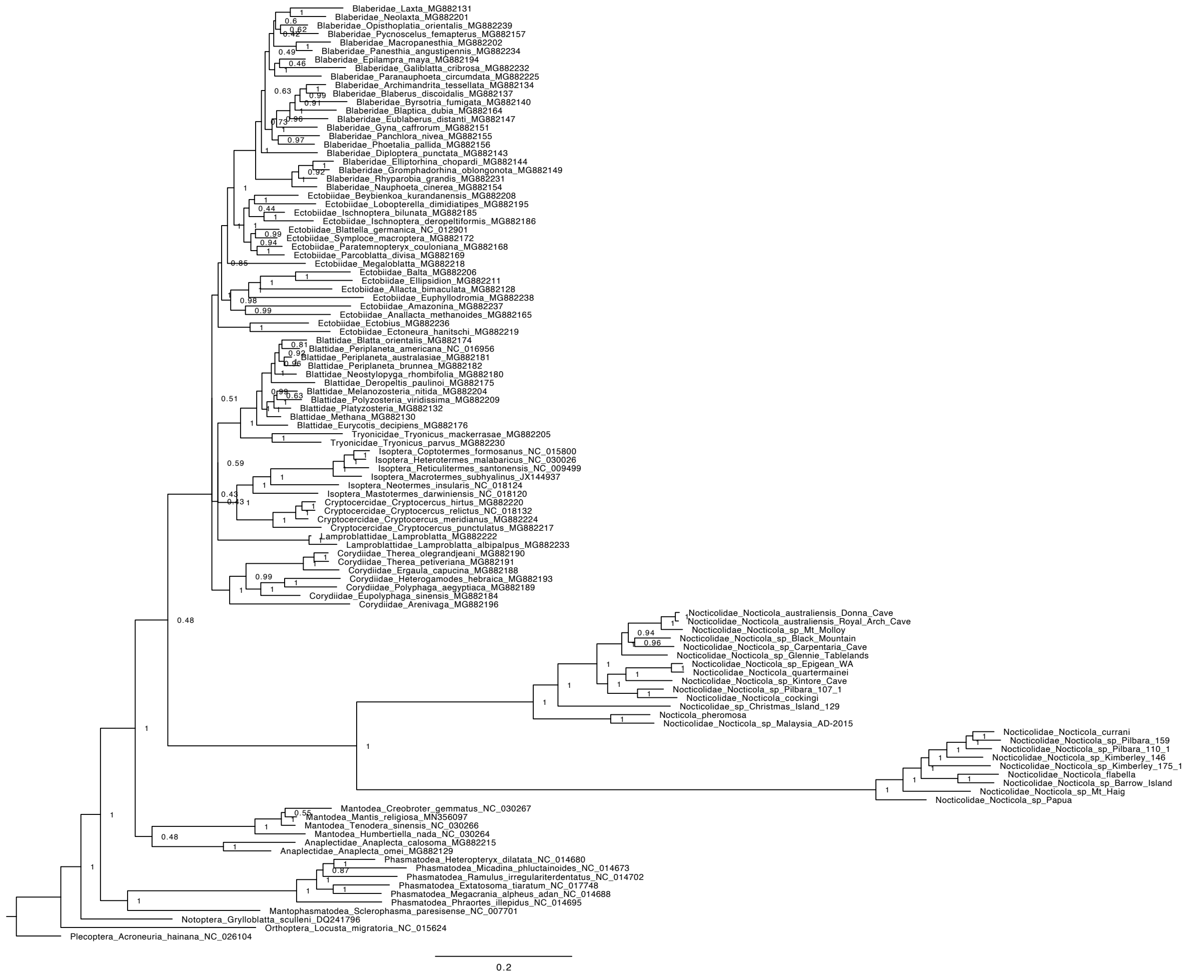


d)

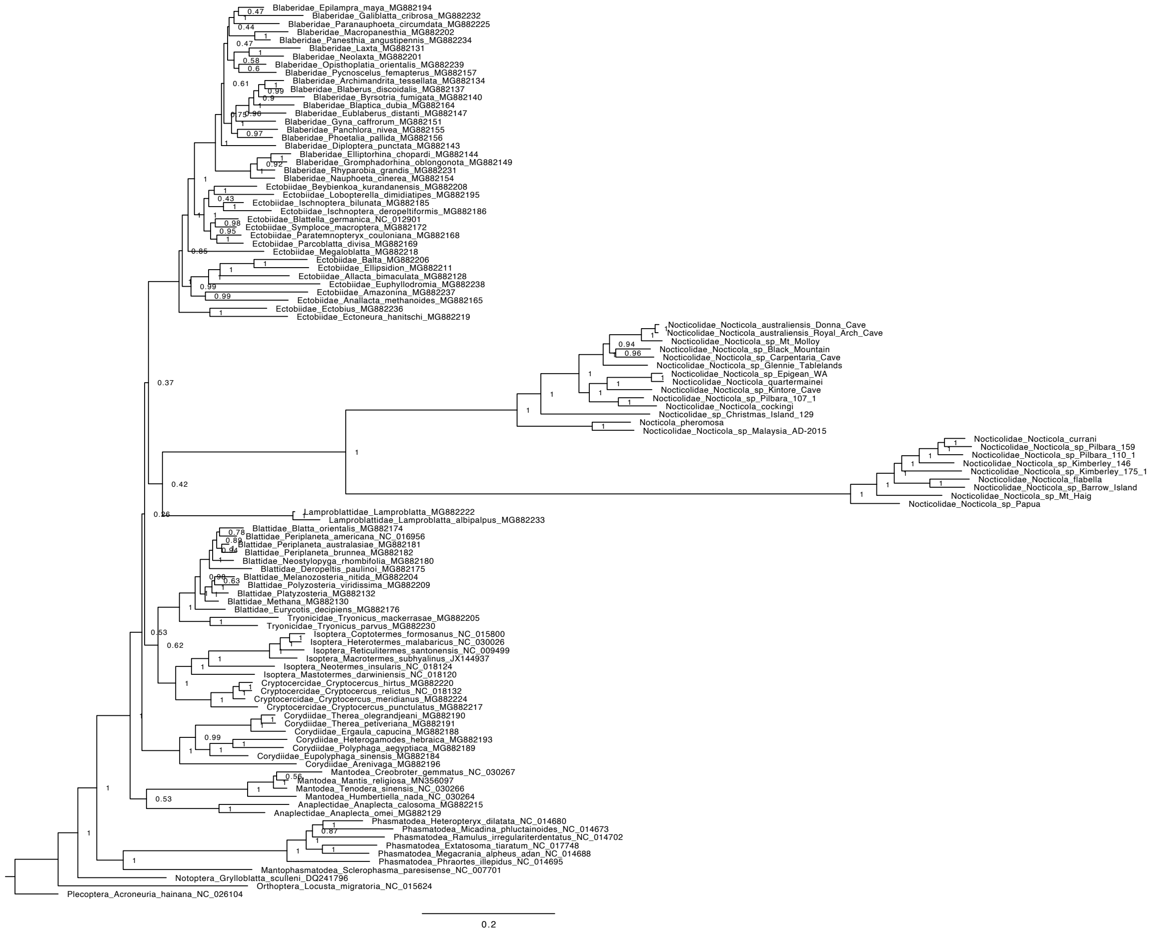


Supplementary Figure S3 continued.

a)

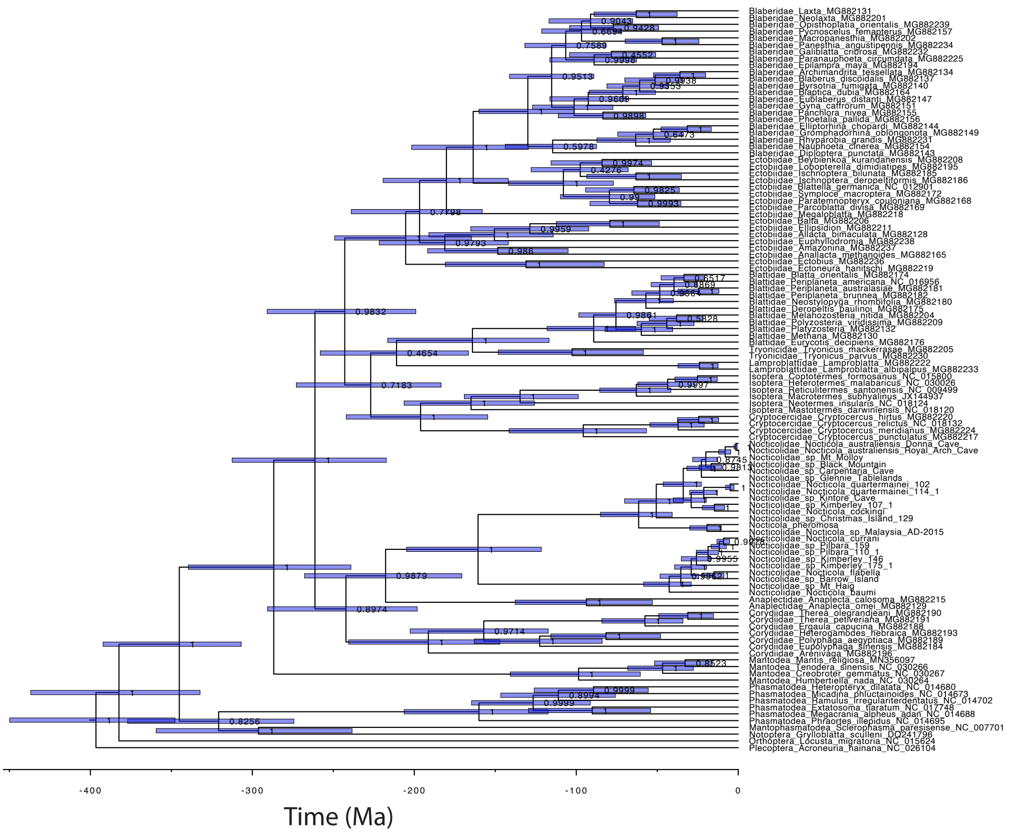


b)

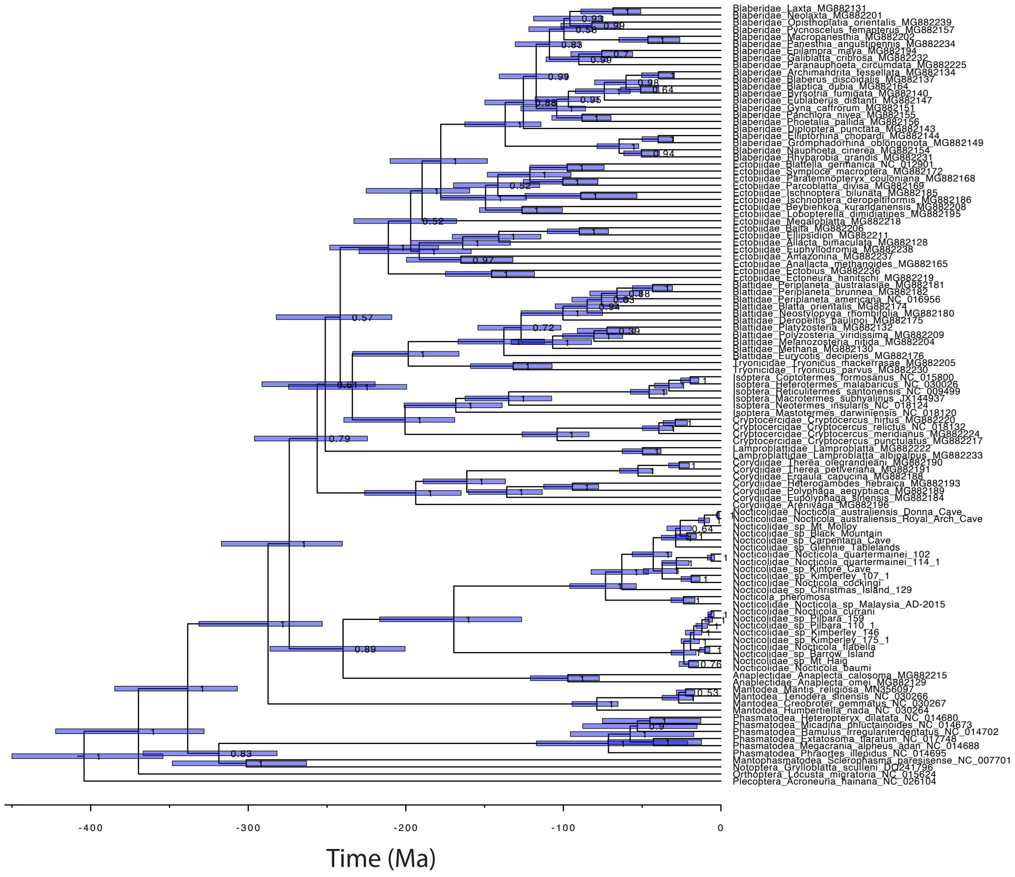


Supplementary Figure S4. Evolutionary timescale of Blattodea inferred in BEAST using the second codon sites of the 13 mitochondrial protein-coding genes. Molecular clock models used include the flexible local clock (a), random local clock (b), uncorrelated exponential relaxed clock (c), uncorrelated lognormal relaxed clock (d) and the strict clock (e). Node values represent posterior probabilities and node bars indicate 95% CI.

c)

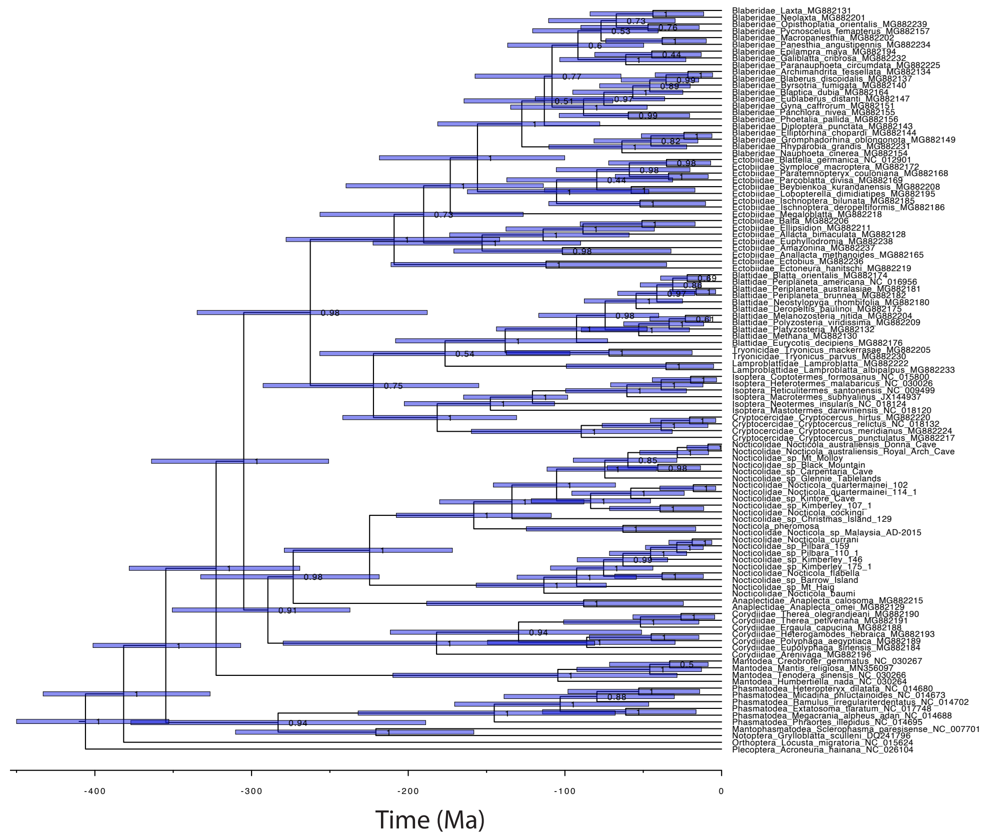


d)

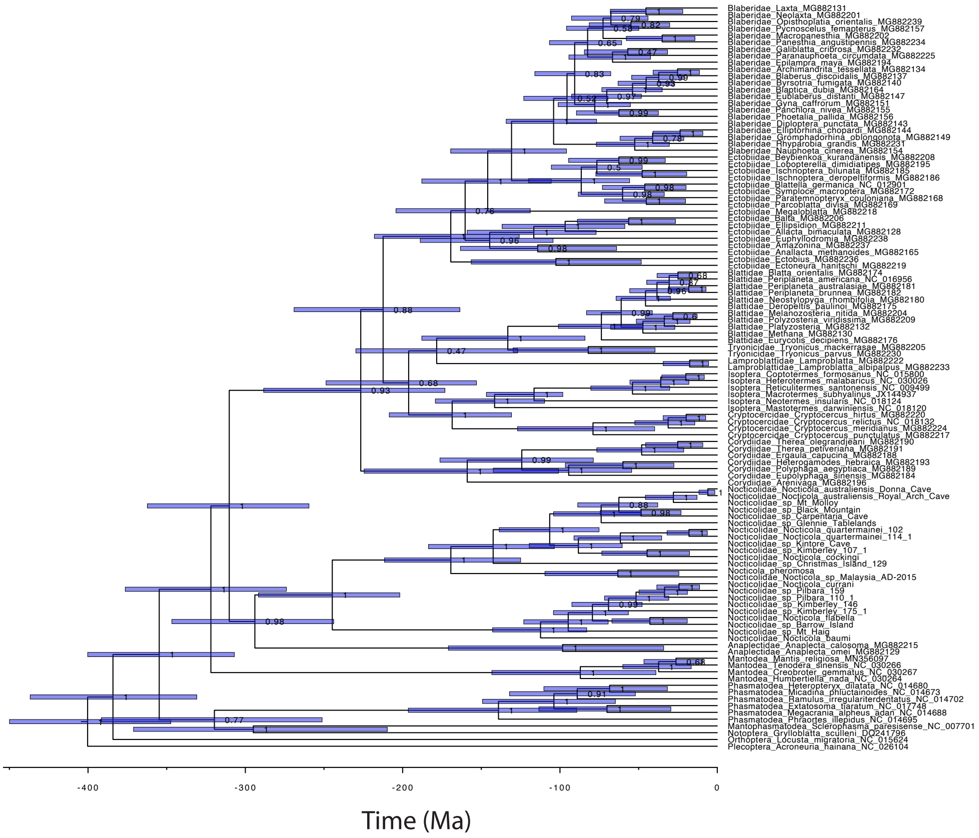


Supplementary Figure S4 continued.

e)

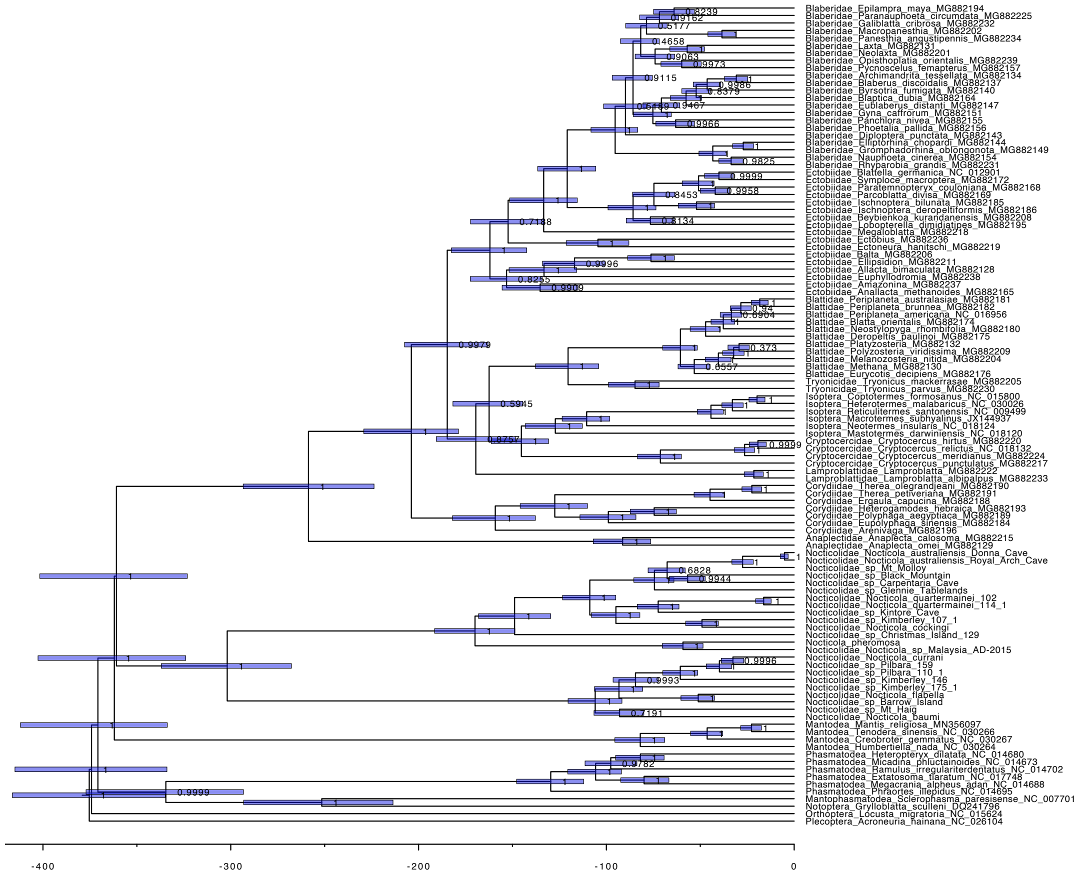


Supplementary Figure S4 continued.

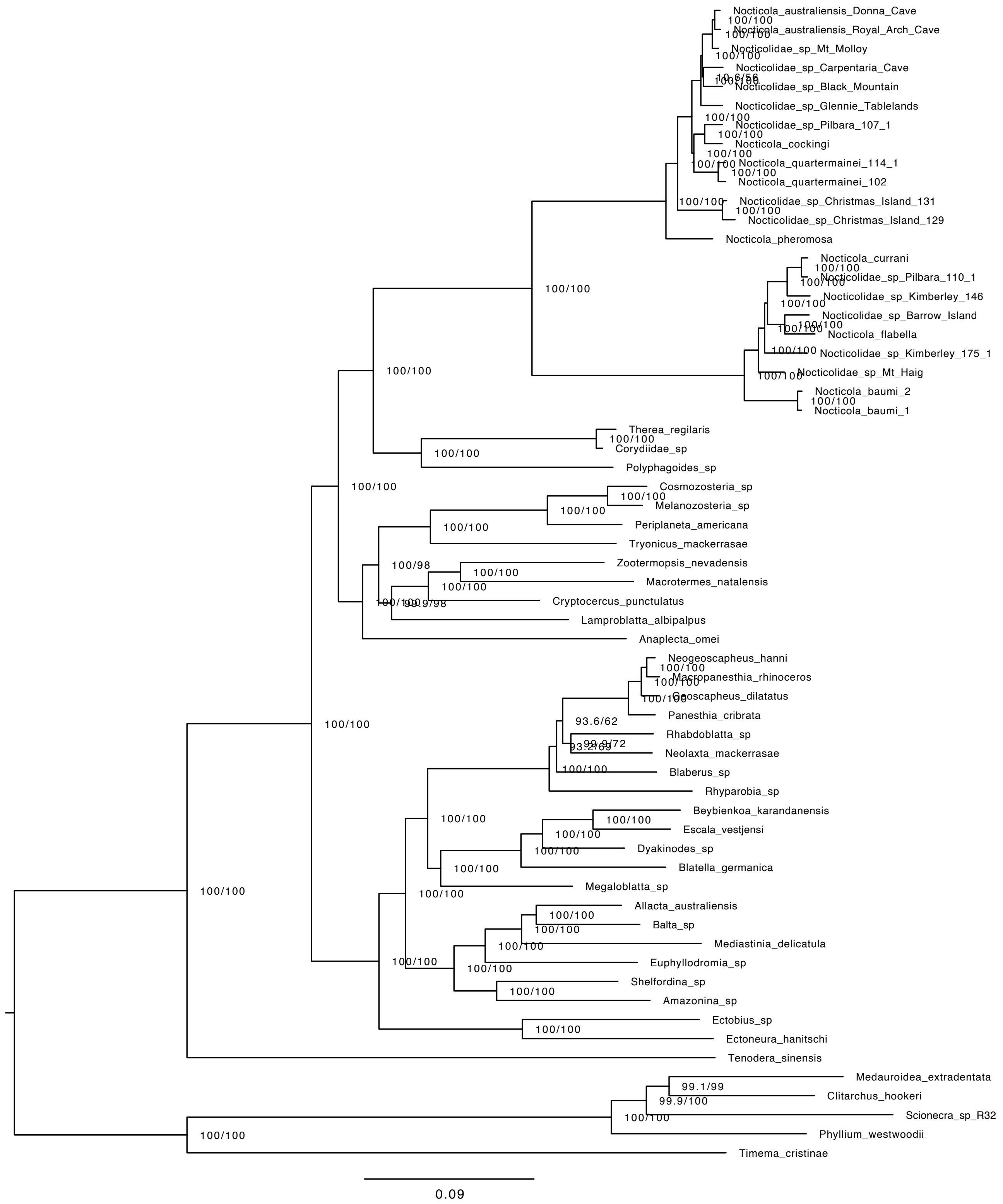


Supplementary Figure S5. Phylogeny of Nocticolidae. Maximum-likelihood tree estimated using concatenated UCE loci. Node values indicate node support as SH-like approximate likelihood-ratio test (SH-alrt) and ultrafast bootstrap (UFb) values. Scale bar represents 0.09 substitutions per site.

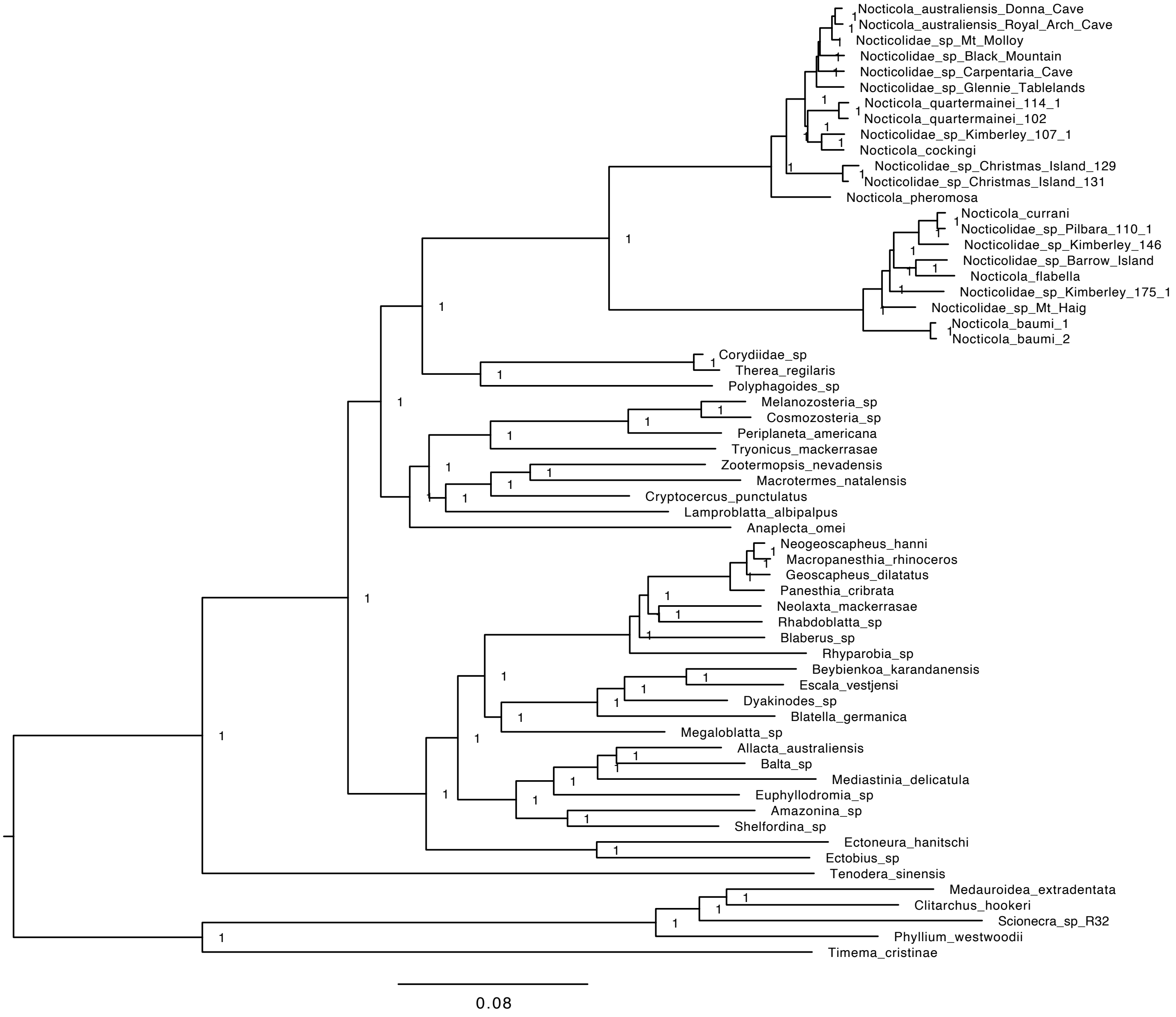


Supplementary Figure S6. Bayesian inference tree of Blattodea inferred using concatenated UCE loci. Scale bar indicates 0.08 substitutions per site. Node values represent posterior probabilities. Infered topologies were the same across the four parallel mcmc simulations.

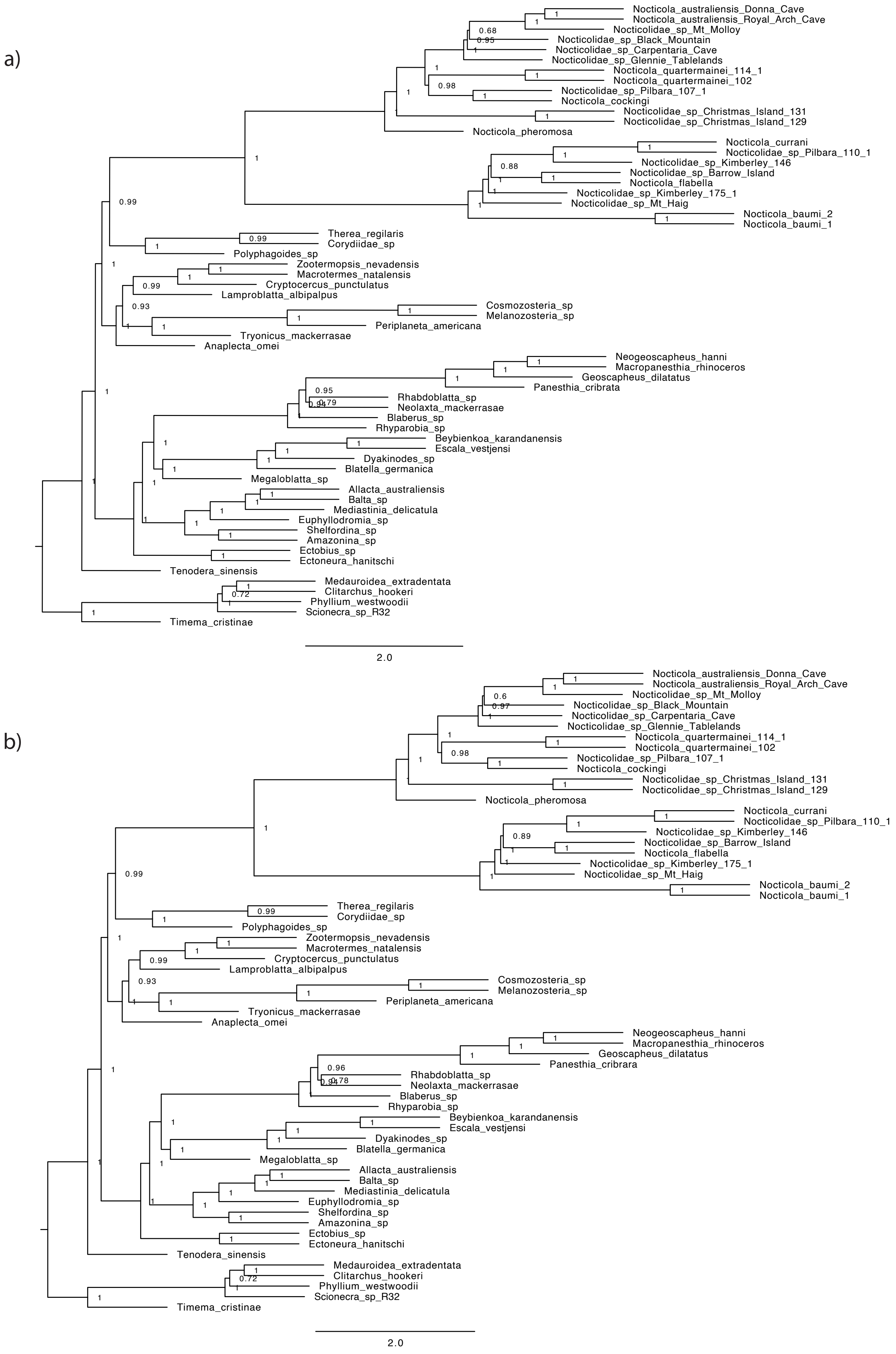


Supplementary Figure S7. Phylogenetic trees inferred using a summary coalescent approach in ASTRAL based on the UCE data set. We produced trees without (a) and with (b) branches in the gene trees contracted if they had bootstrap support lower than 10%. Node values represent local posterior probability. Scale bar represents 2.0 coalescent time units.

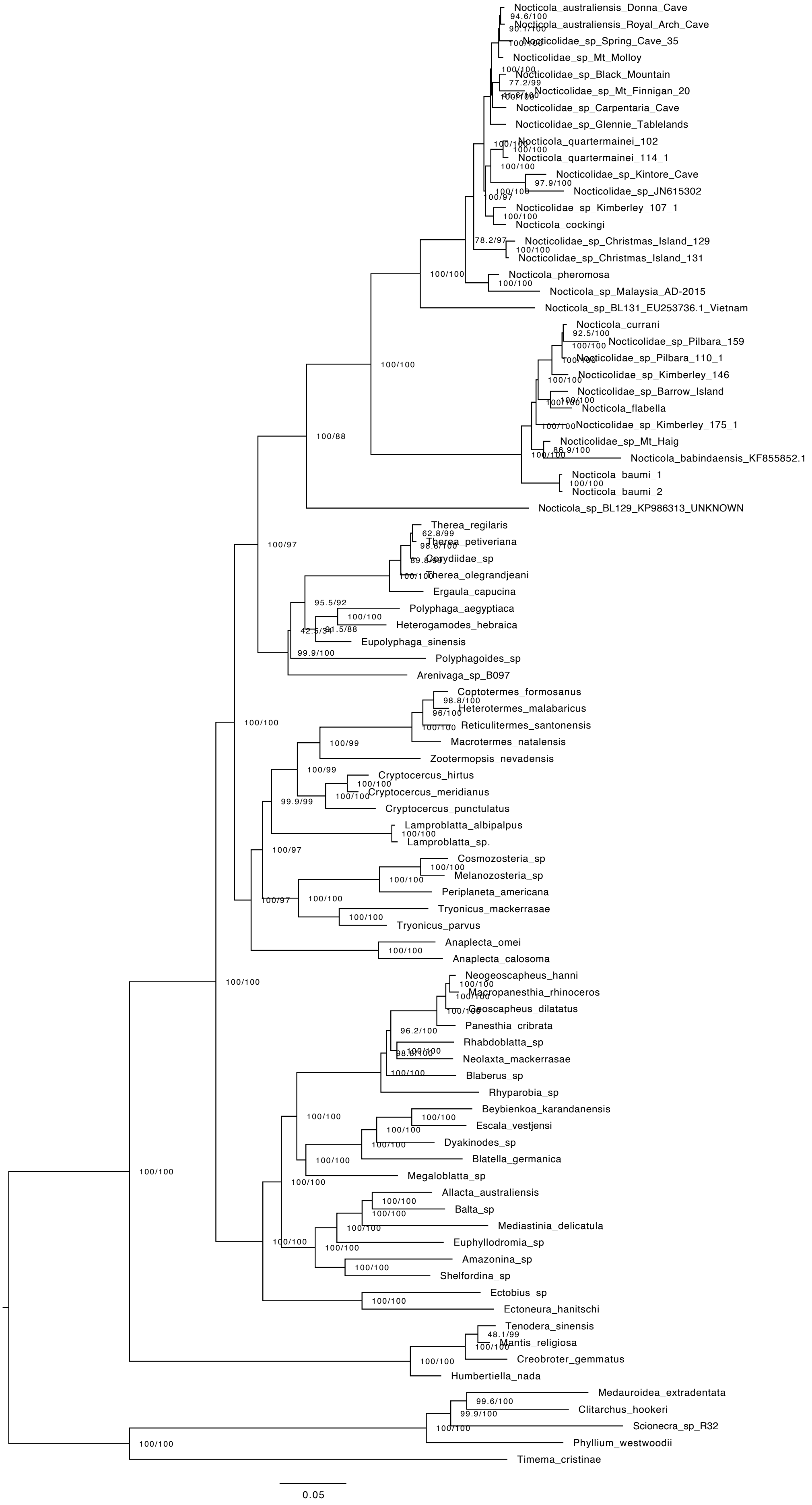


Supplementary Figure S8. Phylogeny of Nocticolidae. Maximum-likelihood tree estimated using concatenated UCE loci, the second codon positions of 13 mitochondrial protein coding genes and the mitochondrial gene encoding 16S rRNA. Node support values indicate the SH-like approximate likelihood-ratio test (SH-alrt) and ultrafast bootstrap (UFb) values. Scale bar represents 0.09 substitutions per site.

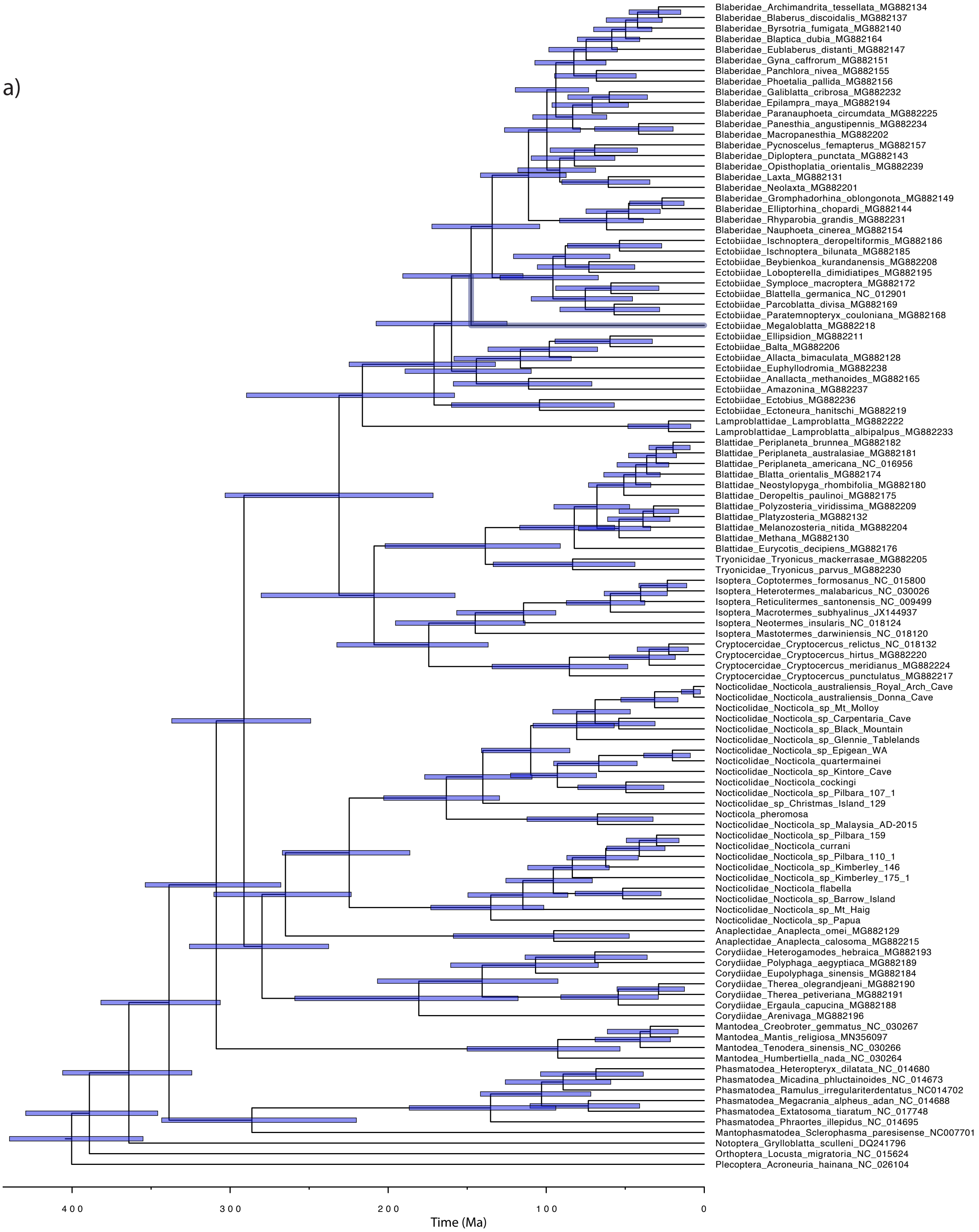


Supplementary Figure S9. Evolutionary timescale of Blattodea, with Mantodea and Phasmatodea as outgroup taxa. Divergence times were estimated based on the second codon positions of 13 mitochondrial protein coding genes, using a Bayesian phylogenomic approach in MCMCtree. Topology was constrained based on the maximum likelihood analysis of mtPCG. Analyses used an independent lognormal relaxed clock (a) and an autocorrelated relaxed clock (b).

b)

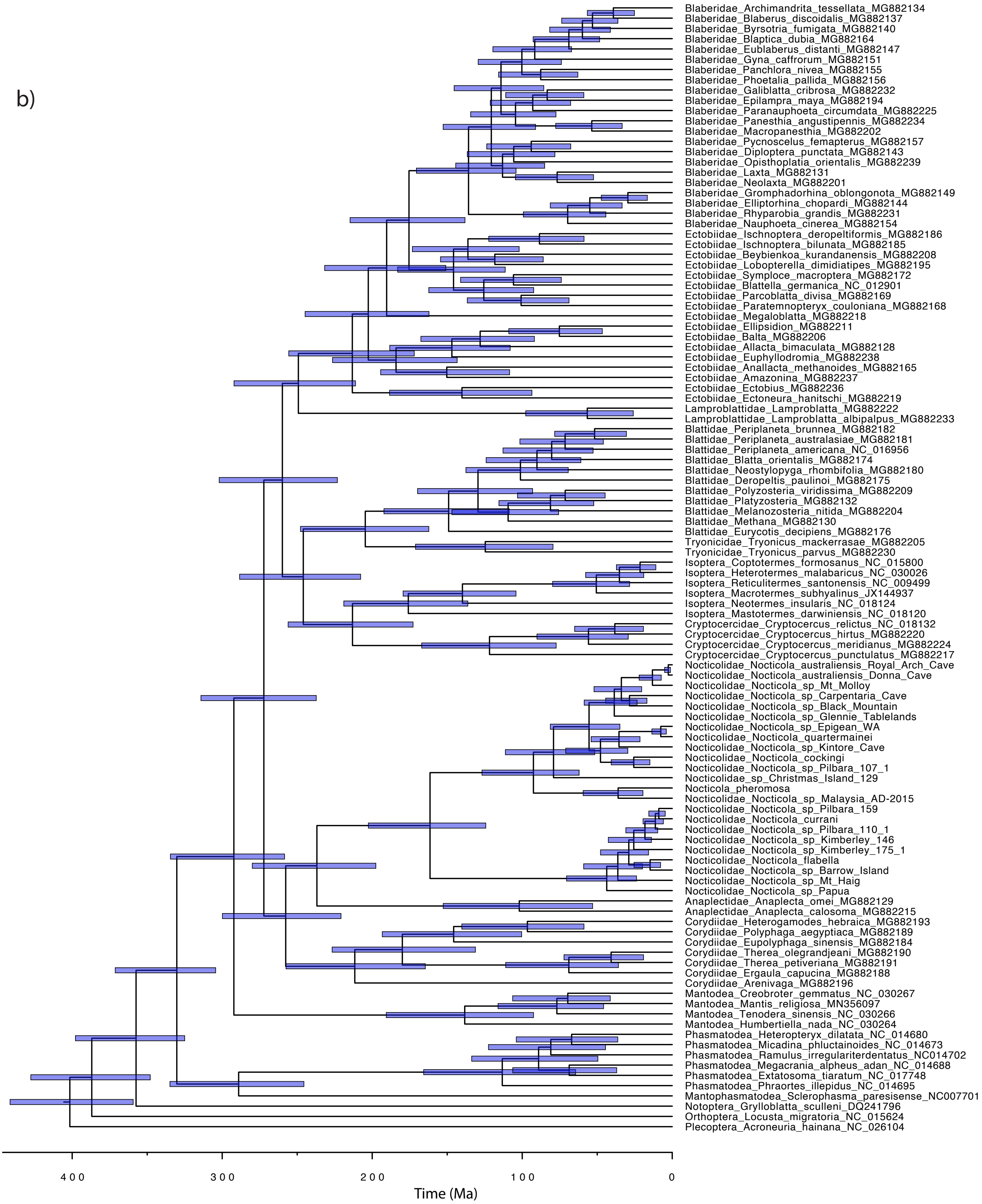


Supplementary Figure S9 continued.

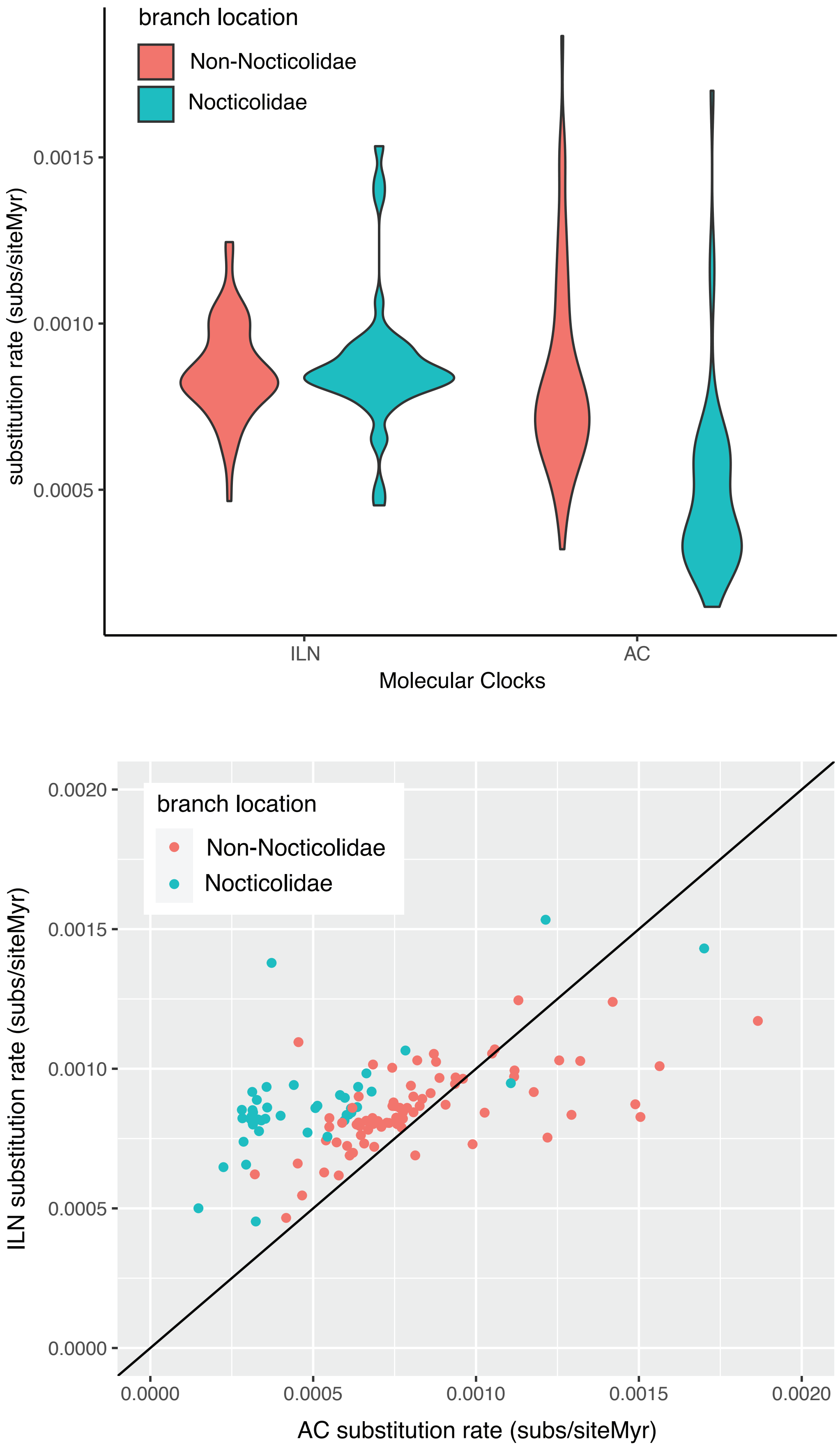


Supplementary Figure S10. Estimates of mitochondrial substitution rates in Nocticolidae compared with other families in Blattodea (cockroaches). a) Violin plots represent the mean substitution rate (substitutions per site per million years) for all terminal and internal branches within Nocticolidae (including stem) and without Nocticolidae (non-Nocticolidae). Rates were estimated using concatenated UCE loci using an independent lognormal relaxed clock (ILN) and autocorrelated relaxed clock (AC). b) comparison of substitution rates inferred for each branch using the ILN and AC models.

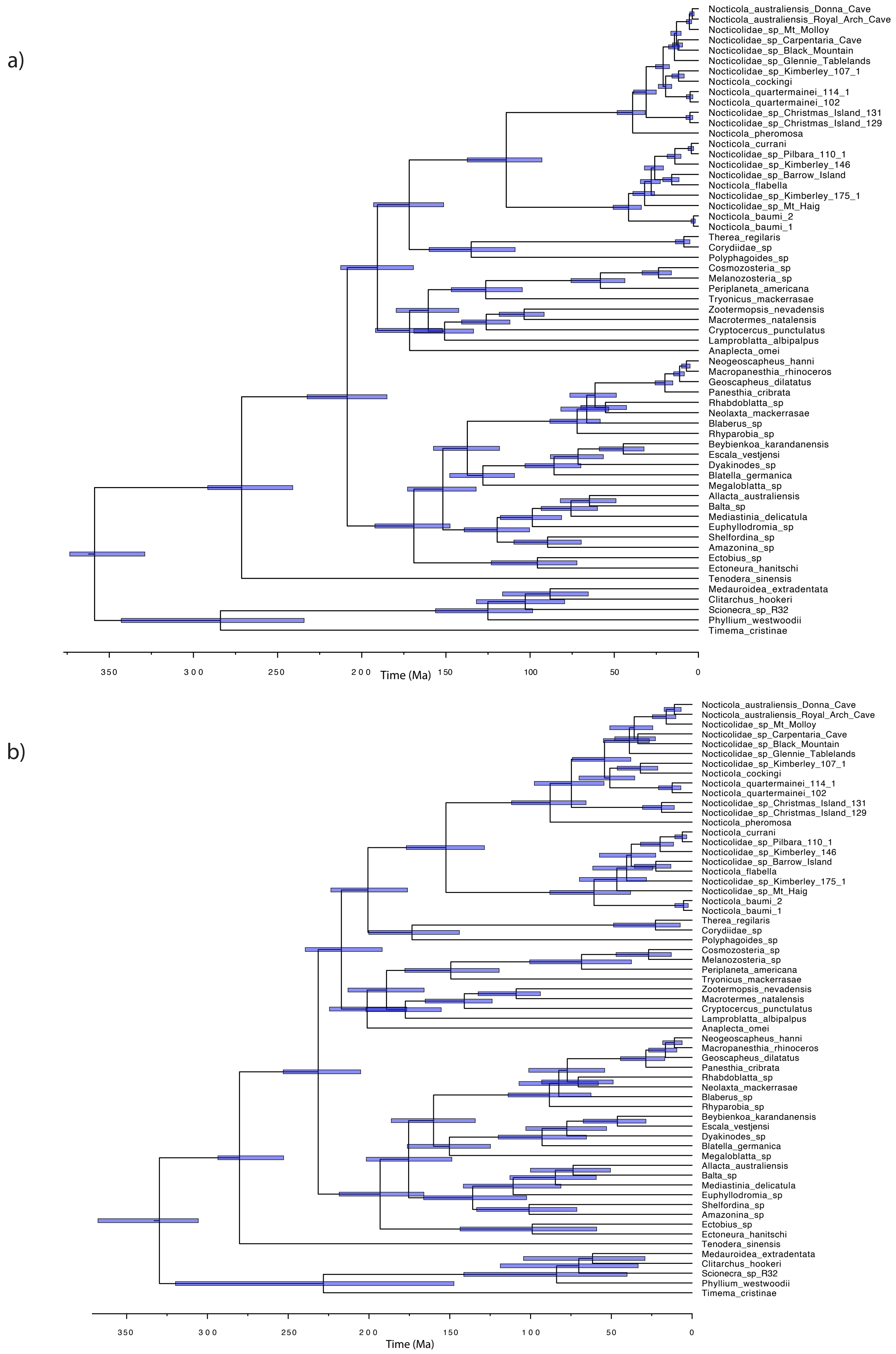


Supplementary Figure S11. Evolutionary timescale of Blattodea, with Mantodea and Phasmatodea as outgroup taxa. Divergence times were estimated based on UCE loci, using a Bayesian phylogenomic approach in MCMCtree. Topology was constrained based on the maximum likelihood analysis of UCE loci. Analyses were completed using an independent lognormal relaxed clock (a) and an autocorrelated relaxed clock (b).

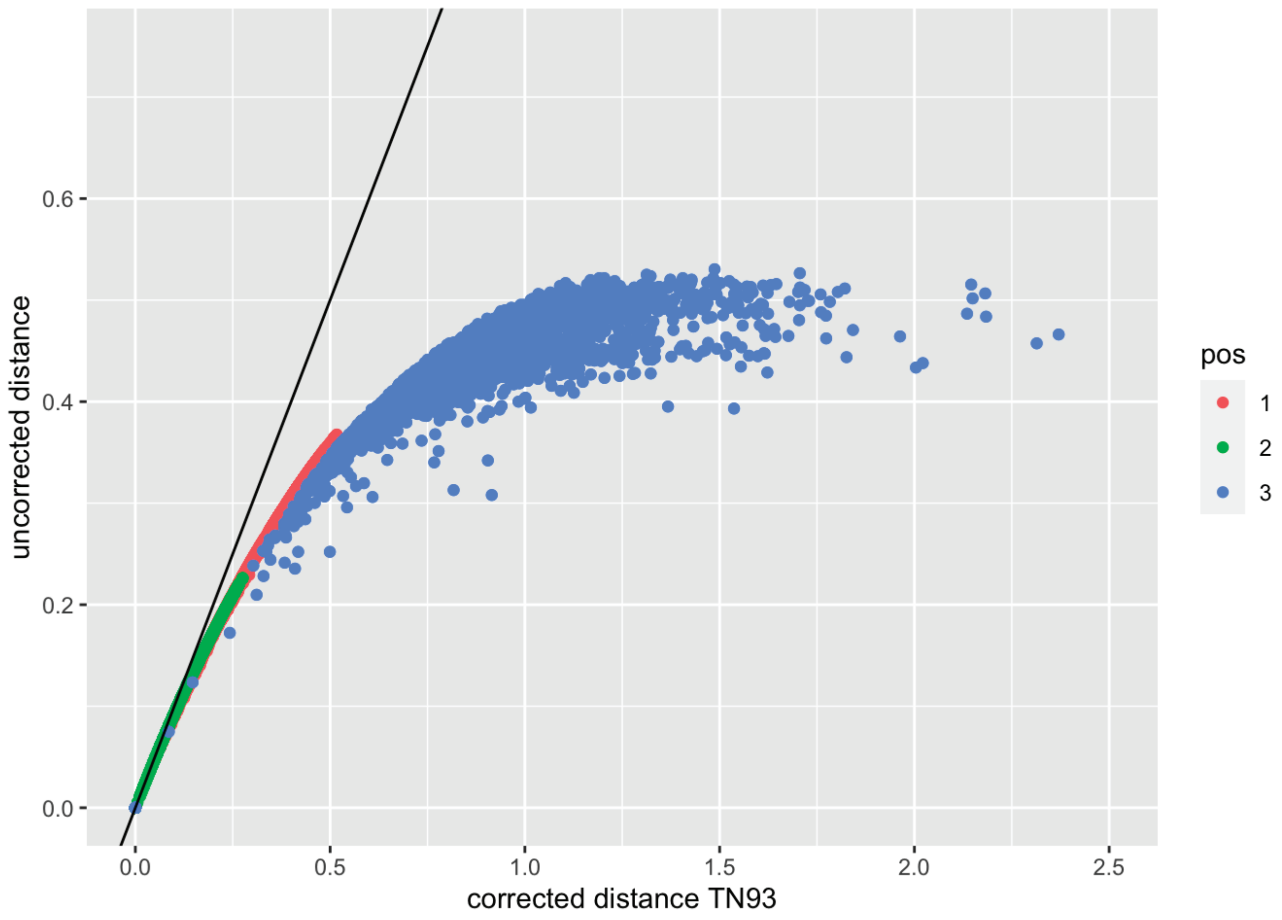


Supplementary Figure S12. Genetic distances between taxa based on the 1st, 2nd and 3rd codon positions of 13 mitochondrial protein coding genes. Uncorrected distances are plotted against distances corrected using the TN93 model. Note that a large number of corrected distances were NA between taxa based on 3rd codon positions. Black line indicates y=x.

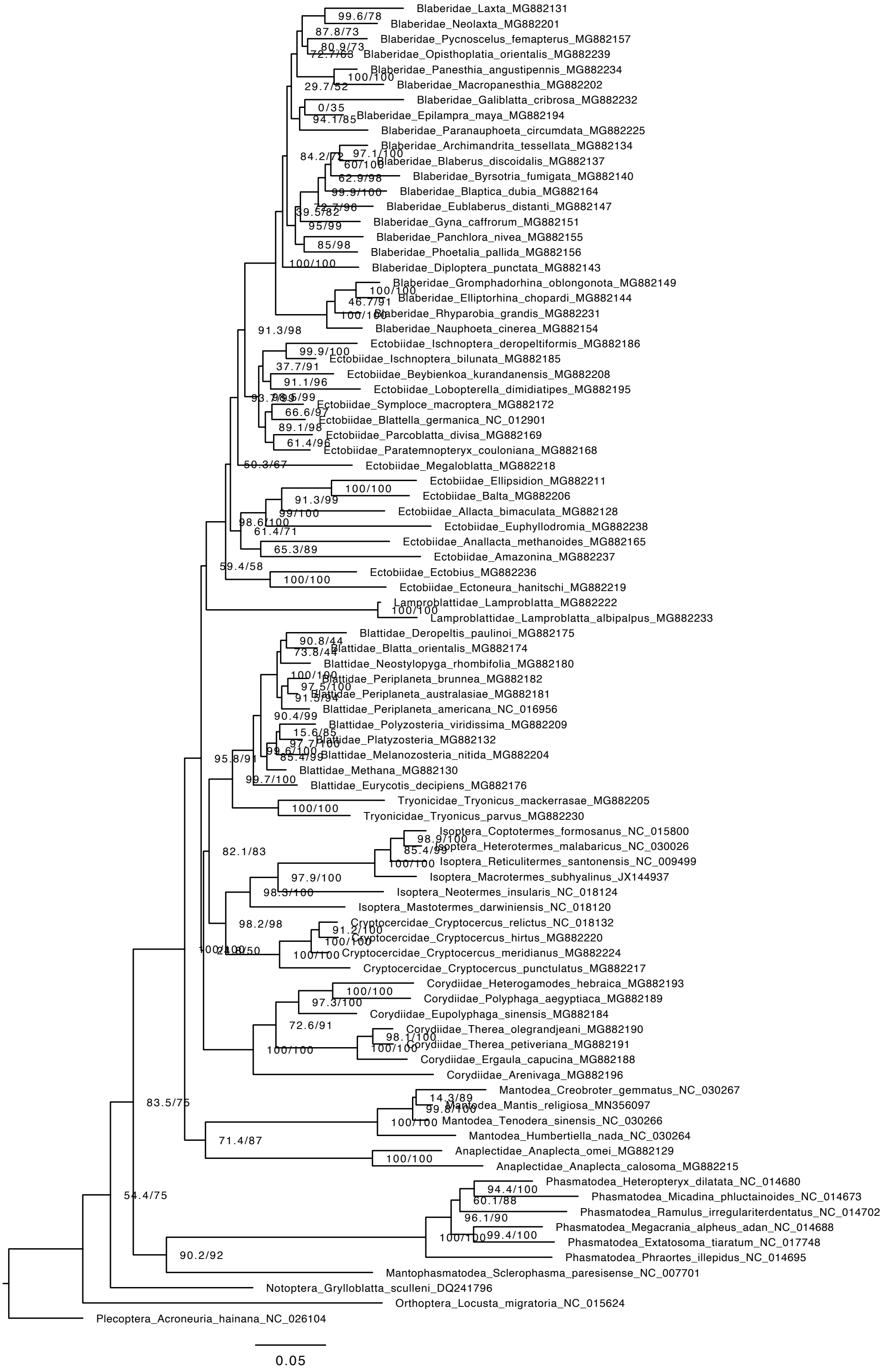


Supplementary Figure S13. Maximum-likelihood tree of Blattodea after removing representatives of Nocticolidae inferred using the second codon sites of the 13 mitochondrial protein-coding genes. Scale bar indicates 0.05 substitutions per site. Node values represent SH-like approximate likelihood-ratio test (SH-alrt) and ultrafast bootstrap (UFb) values.

a)

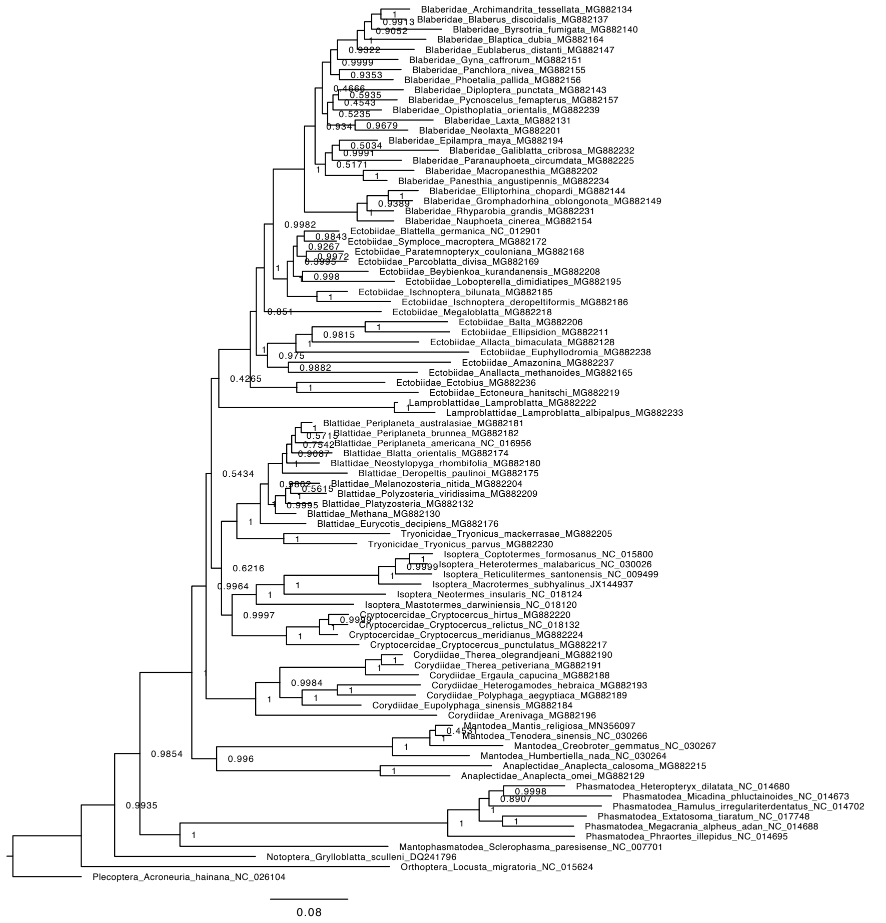


b)

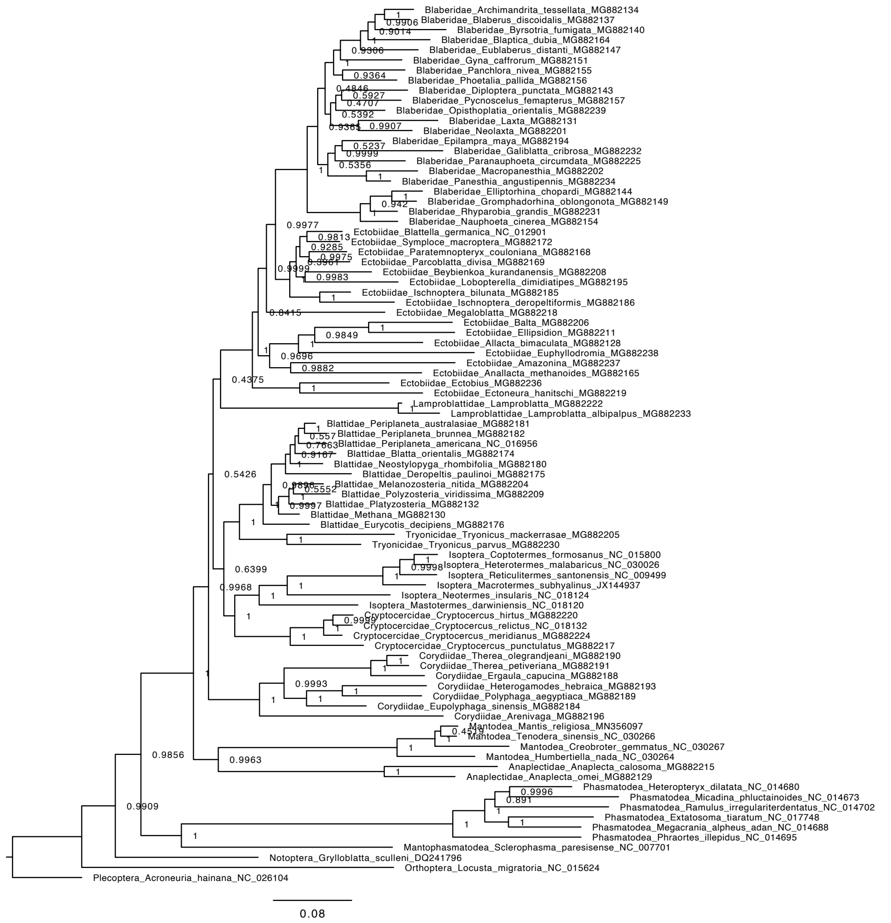


Supplementary Figure S14. Bayesian inference tree of Blattodea after removing representatives of Nocticolidae inferred using the second codon sites of the 13 mitochondrial protein-coding genes. Scale bar indicates 0.08 substitutions per site. Node values represent posterior probabilities. a-d indicate the differing topologies inferred from the four parallel mcmc simulations.

c)

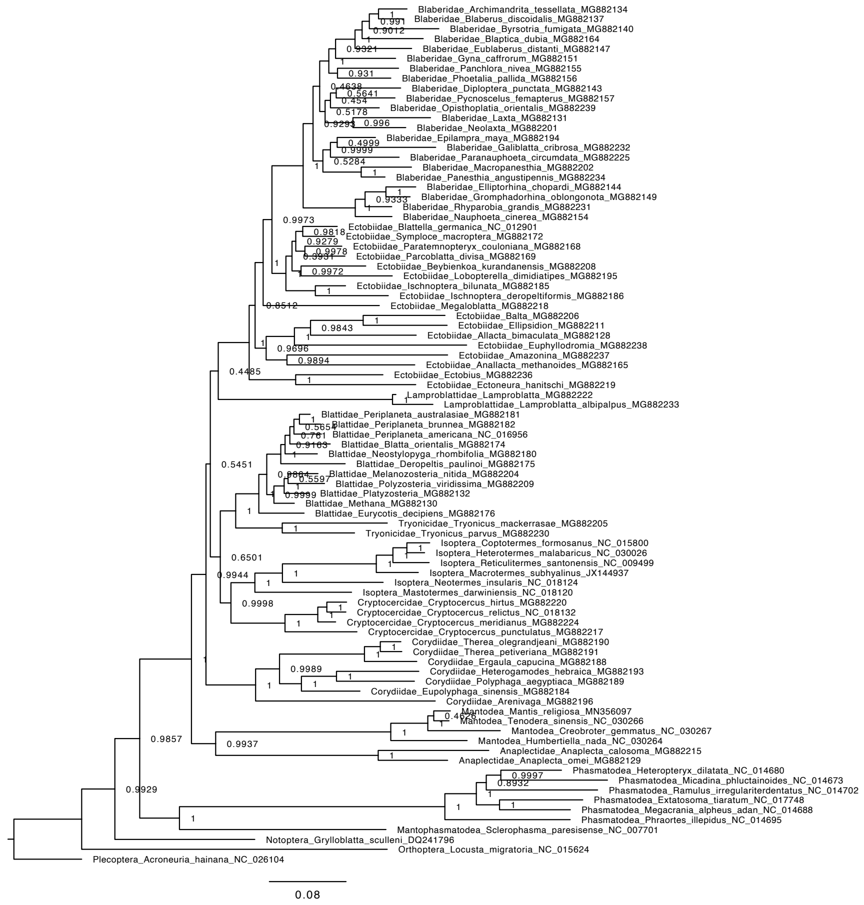


d)

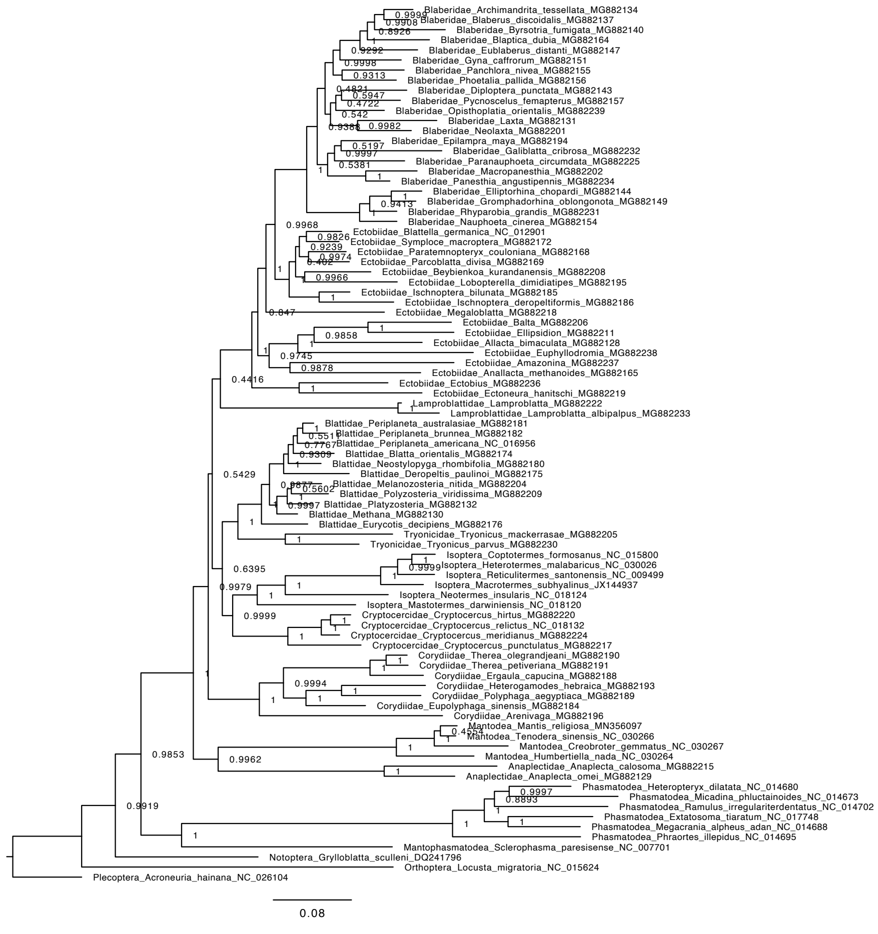


Supplementary Figure S14 continued.

a)

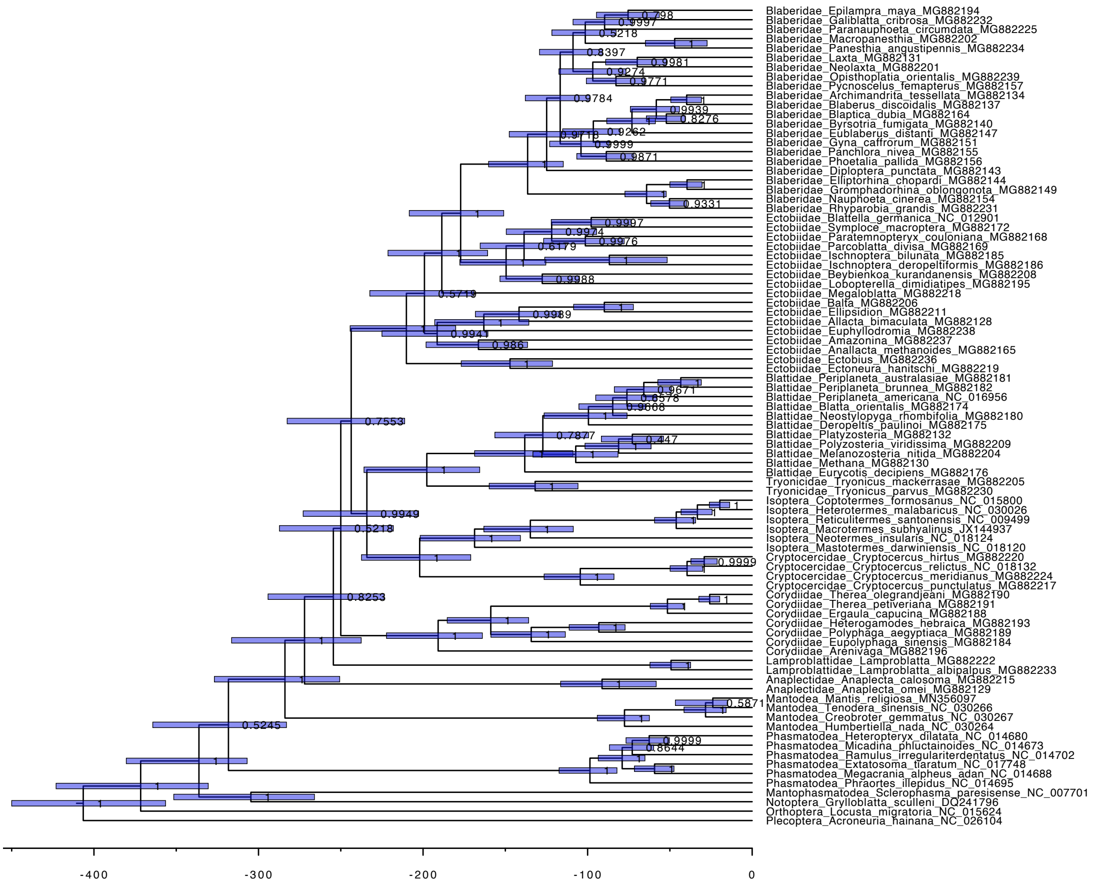


b)

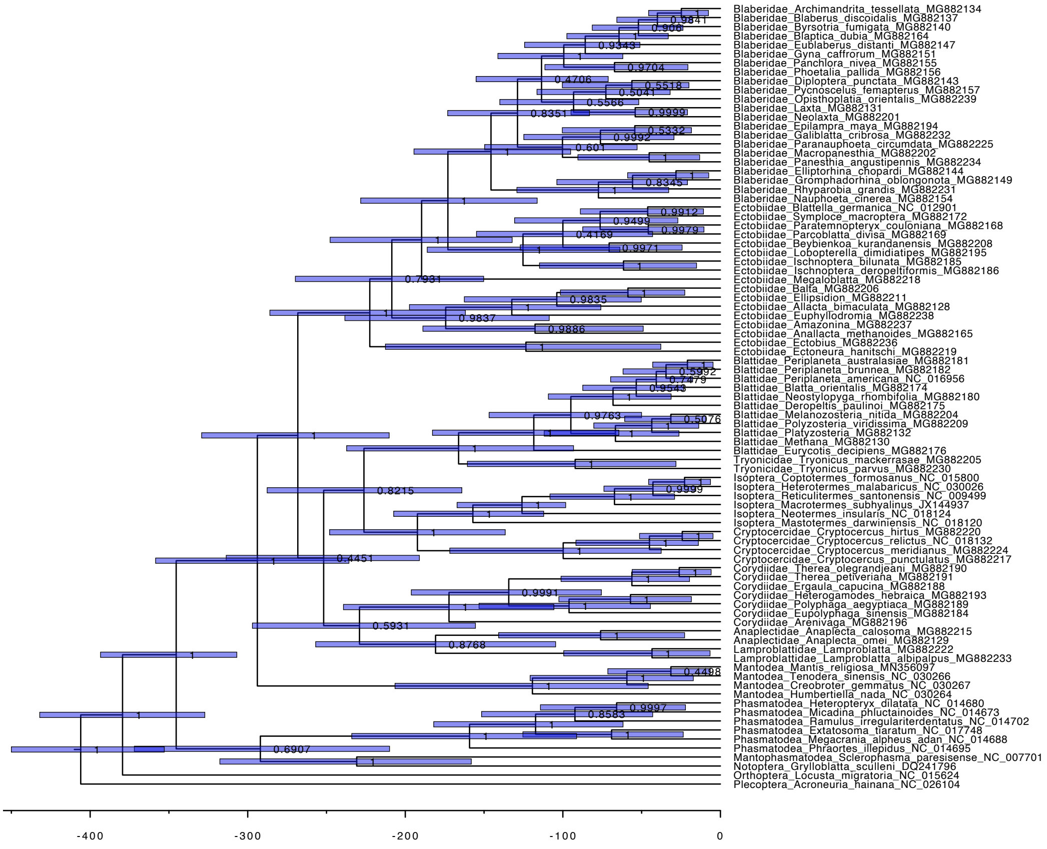


Supplementary Figure S15. Evolutionary timescale of Blattodea inferred in BEAST using the second codon sites of the 13 mitochondrial protein-coding genes. Molecular clock models used include the random local clock (a), uncorrelated exponential relaxed clock (b), uncorrelated lognormal relaxed clock (c) and the strict clock (d). Node values represent posterior probabilities and node bars indicate 95% CI.

c)

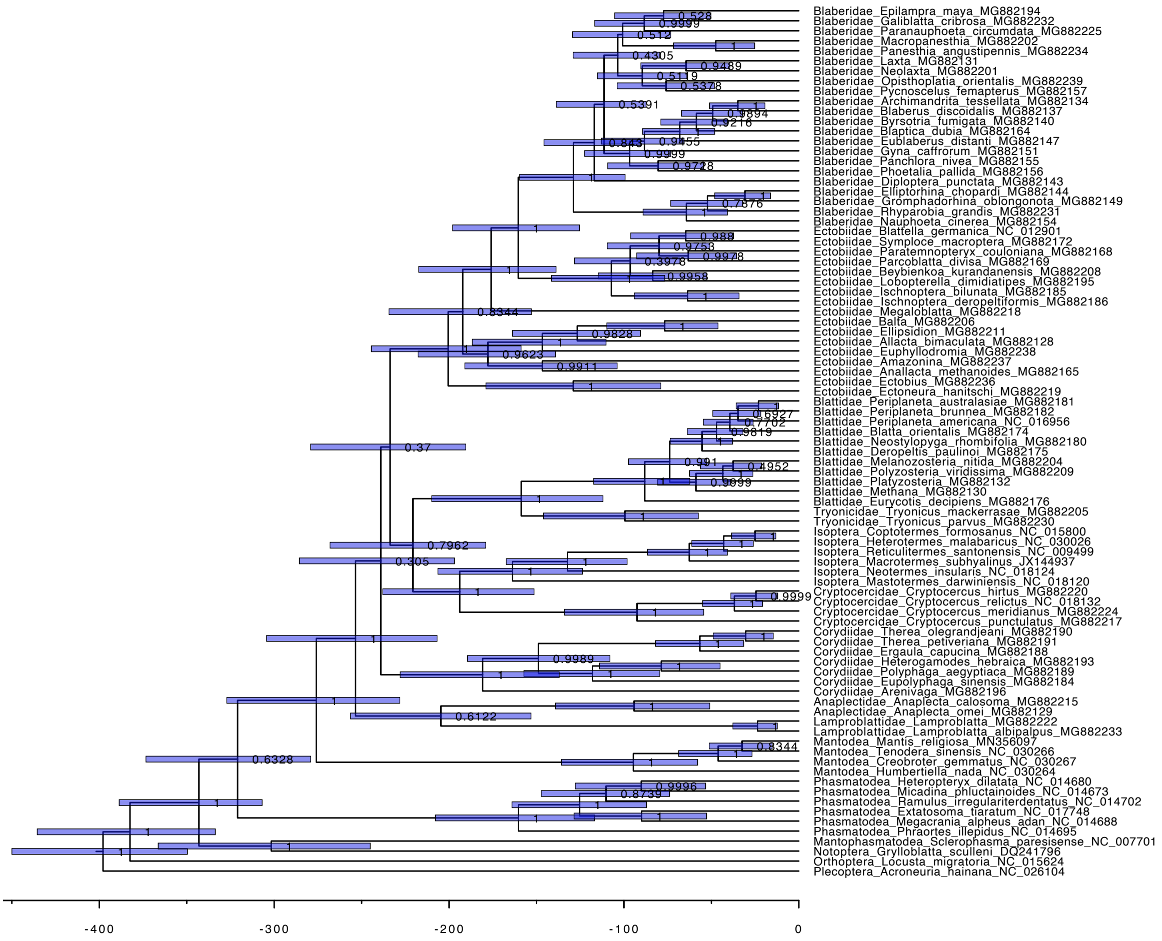


d)

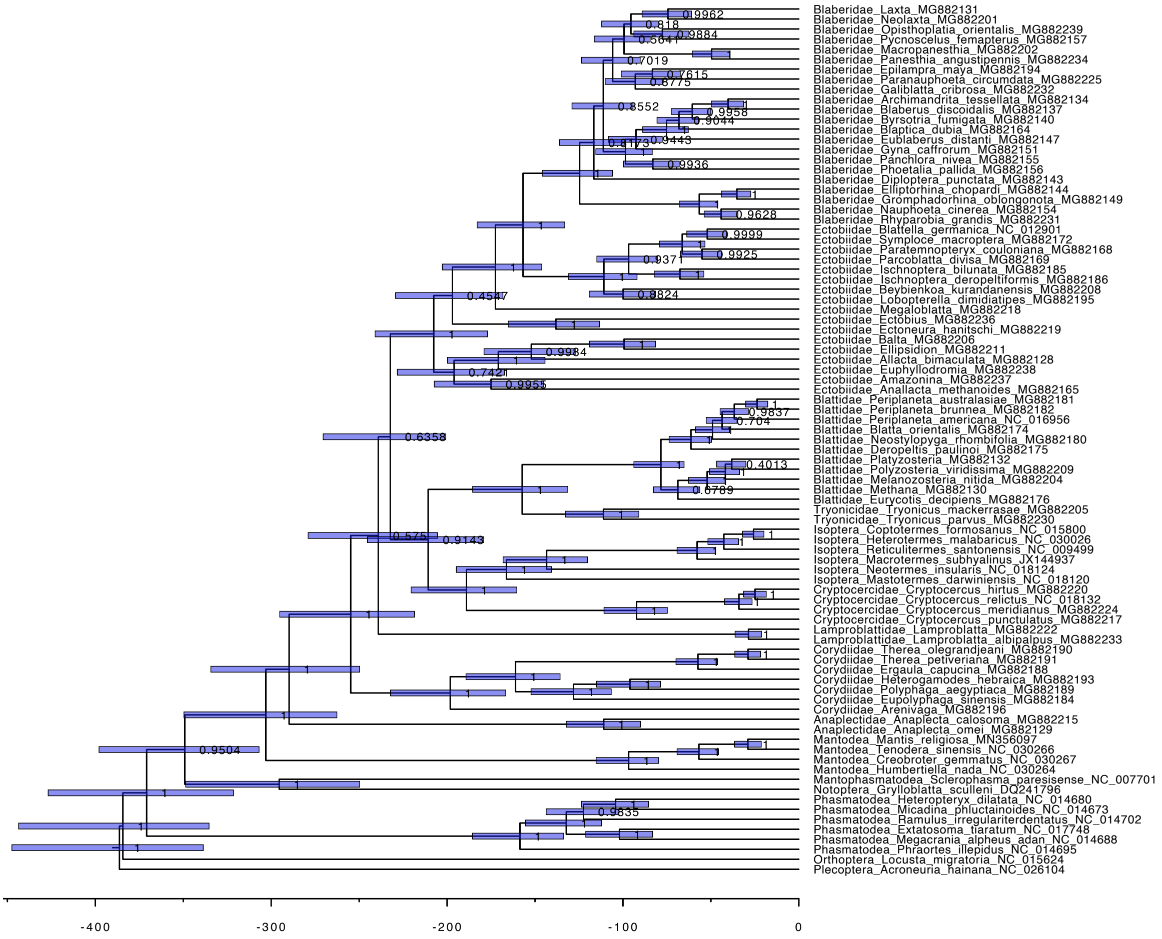


Supplementary Figure S15 continued.

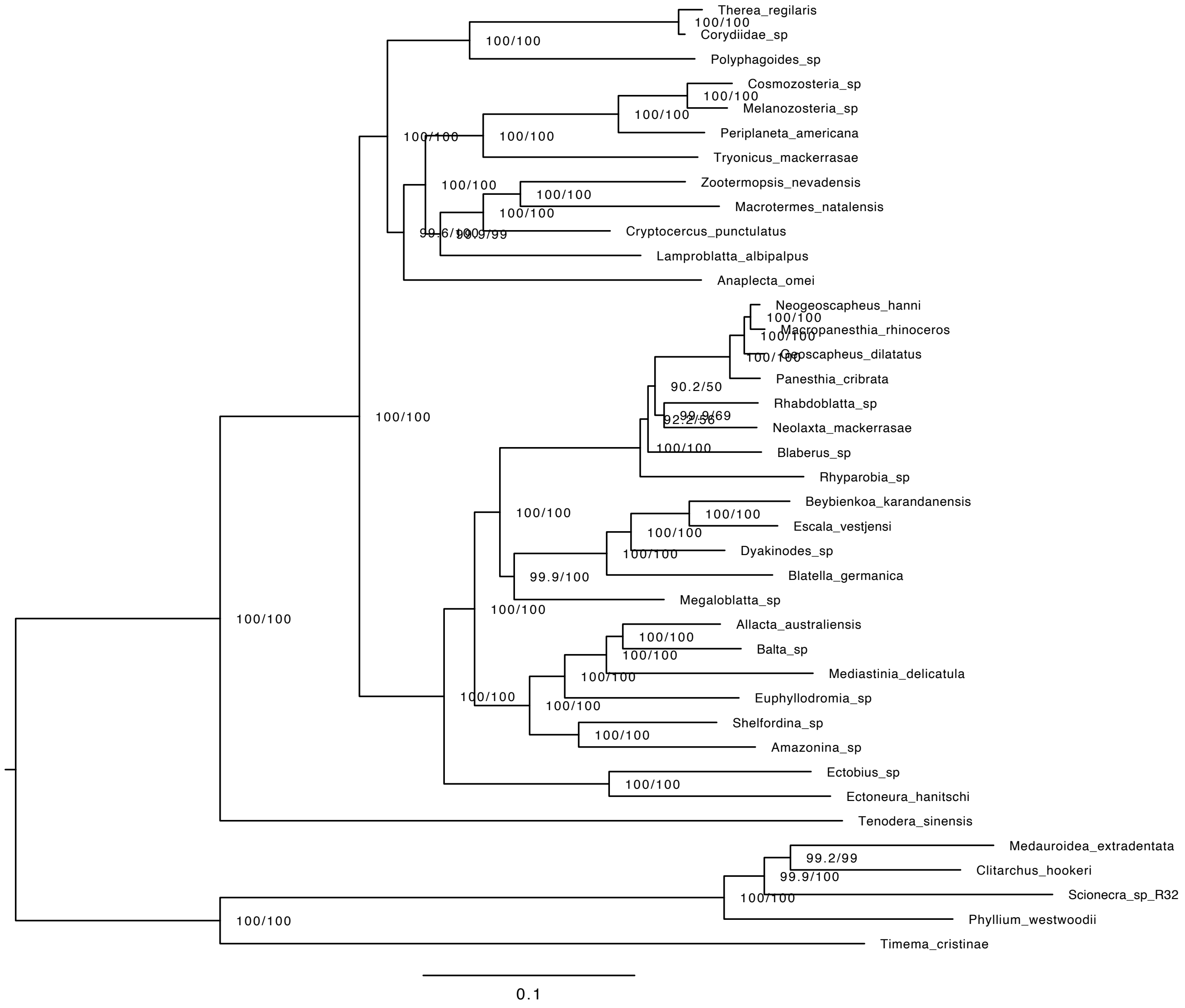


Supplementary Figure S16. Phylogeny of Blattodea after removing representatives of Nocticolidae. Maximum-likelihood tree estimated using concatenated UCE loci. Node values indicate node support as SH-like approximate likelihood-ratio test (SH-alrt) and ultrafast bootstrap (UFb) values. Scale bar represents 0.1 substitutions per site.

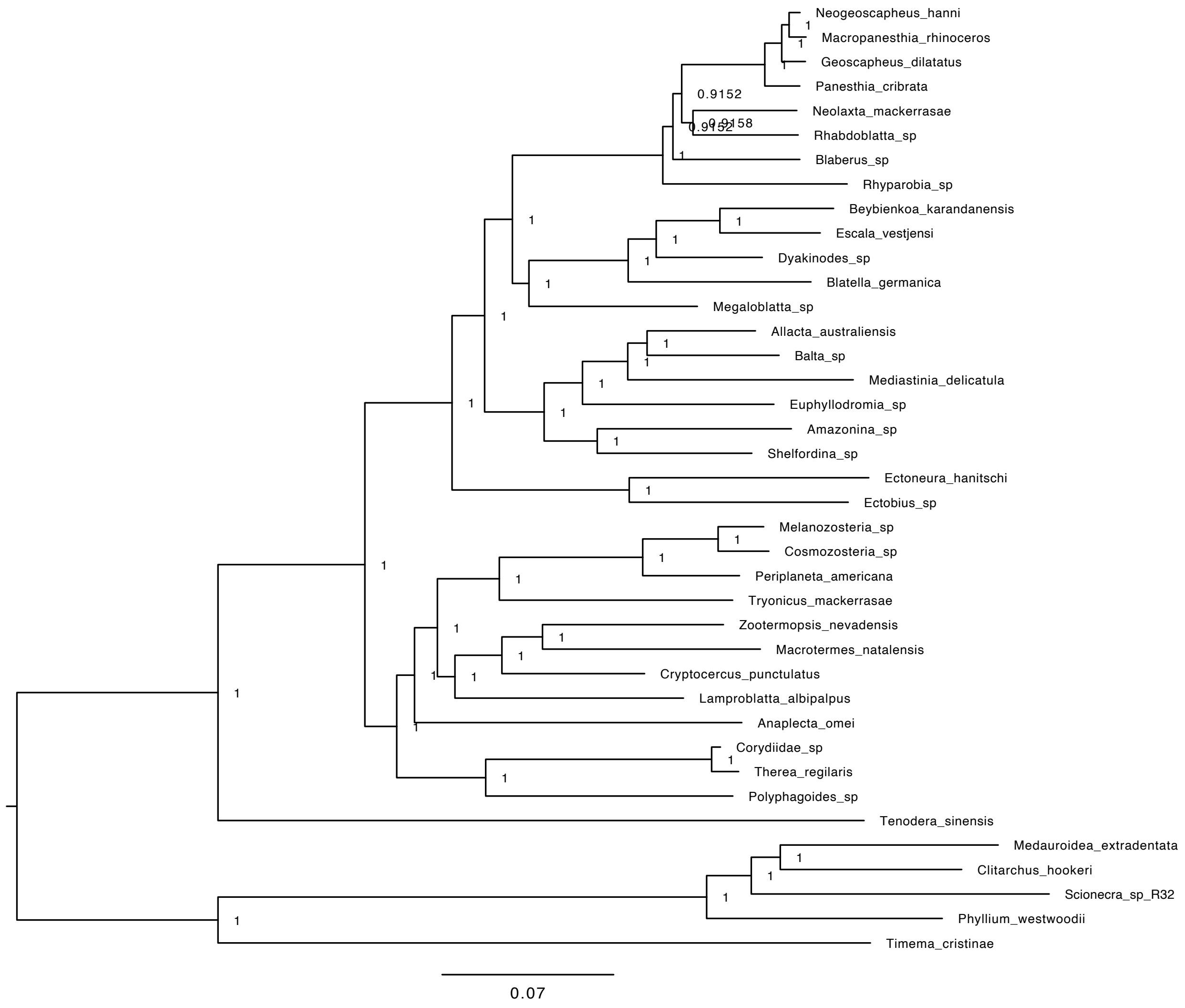


Supplementary Figure S17. Bayesian inference tree of Blattodea after removing representatives of Nocticolidae inferred using concatenated UCE loci. Scale bar indicates 0.07 substitutions per site. Node values represent posterior probabilities. Infered topologies were the same across the four parallel mcmc simulations.

a)

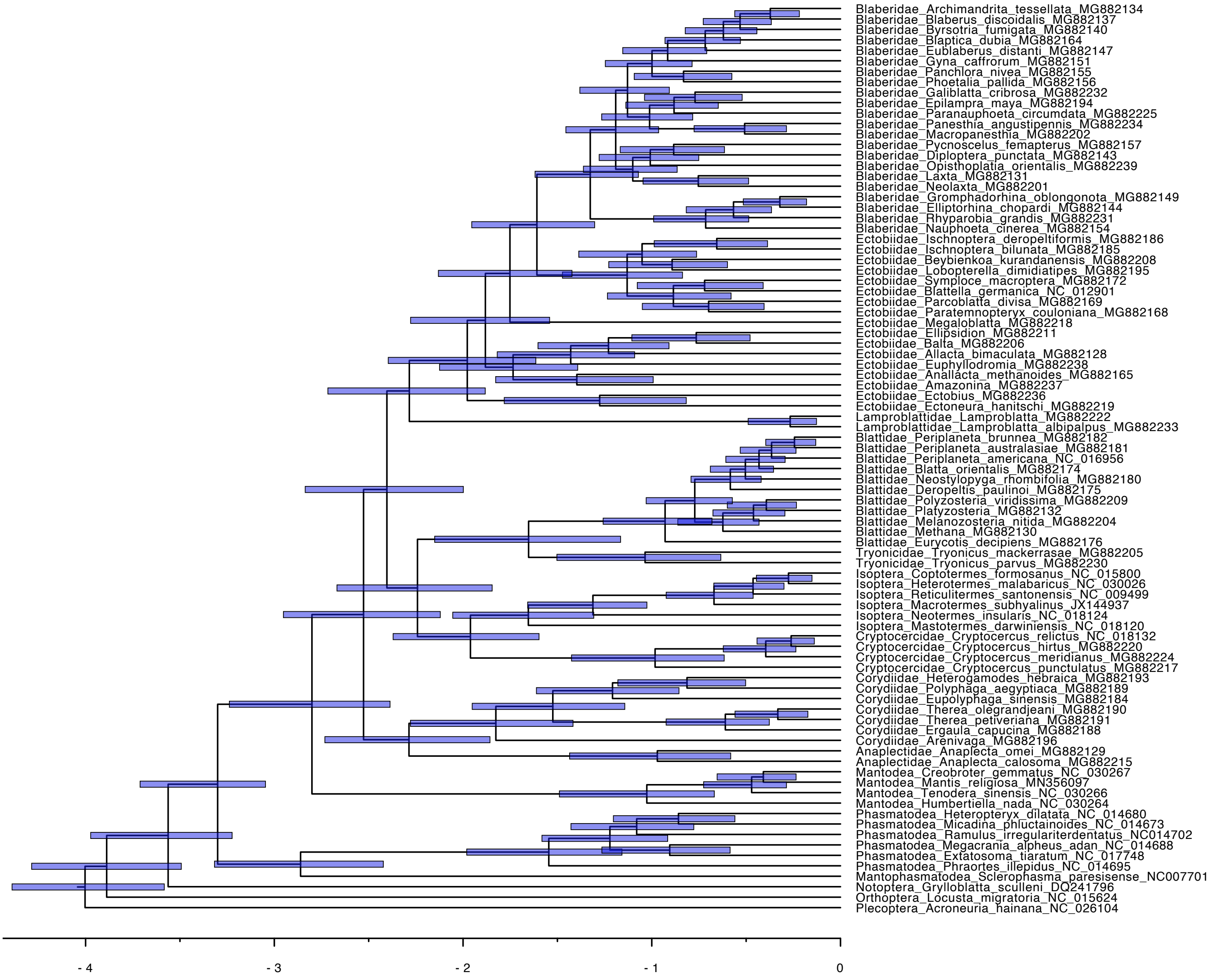


Supplementary Figure S18. Evolutionary timescale of Blattodea after removing representatives of Nocticolidae, with Mantodea and Phasmatodea as outgroup taxa. Divergence times were estimated based on the second codon positions of 13 mitochondrial protein coding genes, using a Bayesian phylogenomic approach in MCMCtree. Topology was constrained based on the maximum likelihood analysis of mtPCG. Analyses used an independent lognormal relaxed clock (a) and an autocorrelated relaxed clock (b).

b)

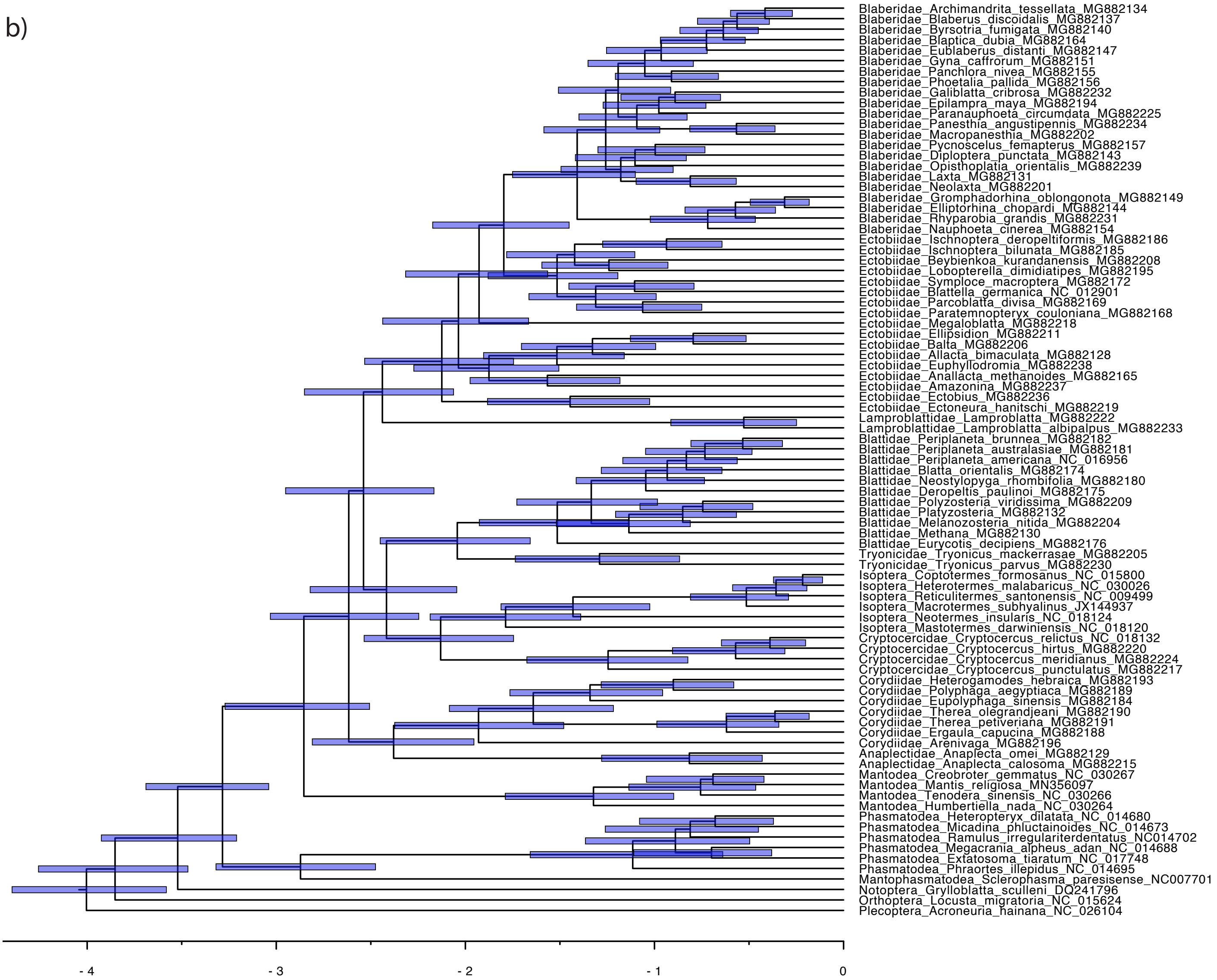


Supplementary Figure S18 continued.

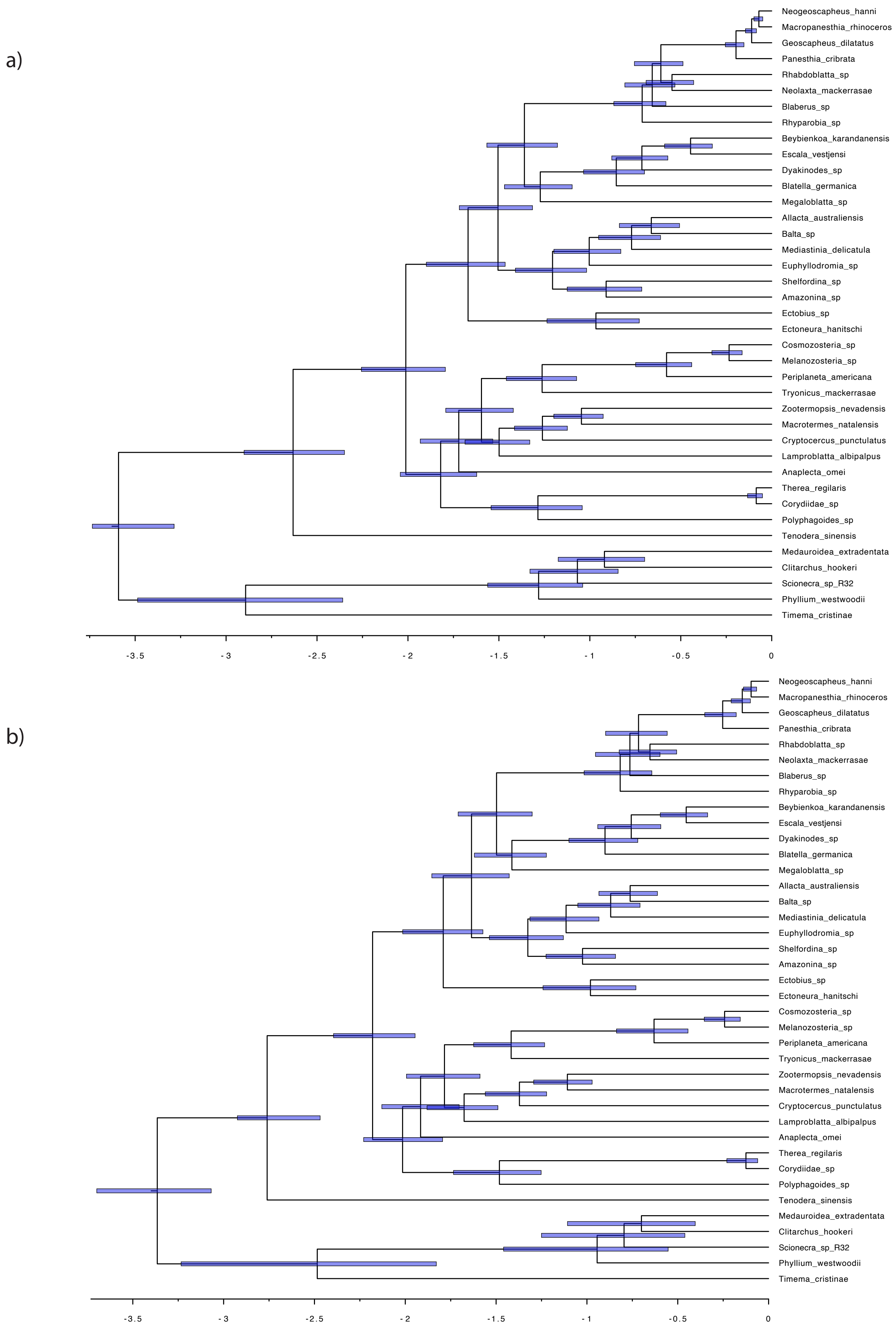


Supplementary Figure S19. Evolutionary timescale of Blattodea after removing representatives of Nocticolidae, with Mantodea and Phasmatodea as outgroup taxa. Divergence times were estimated based on UCE loci, using a Bayesian phylogenomic approach in MCMCtree. Topology was constrained based on the maximum likelihood analysis of UCE loci. Analyses were completed using an independent lognormal relaxed clock (a) and an autocorrelated relaxed clock (b).

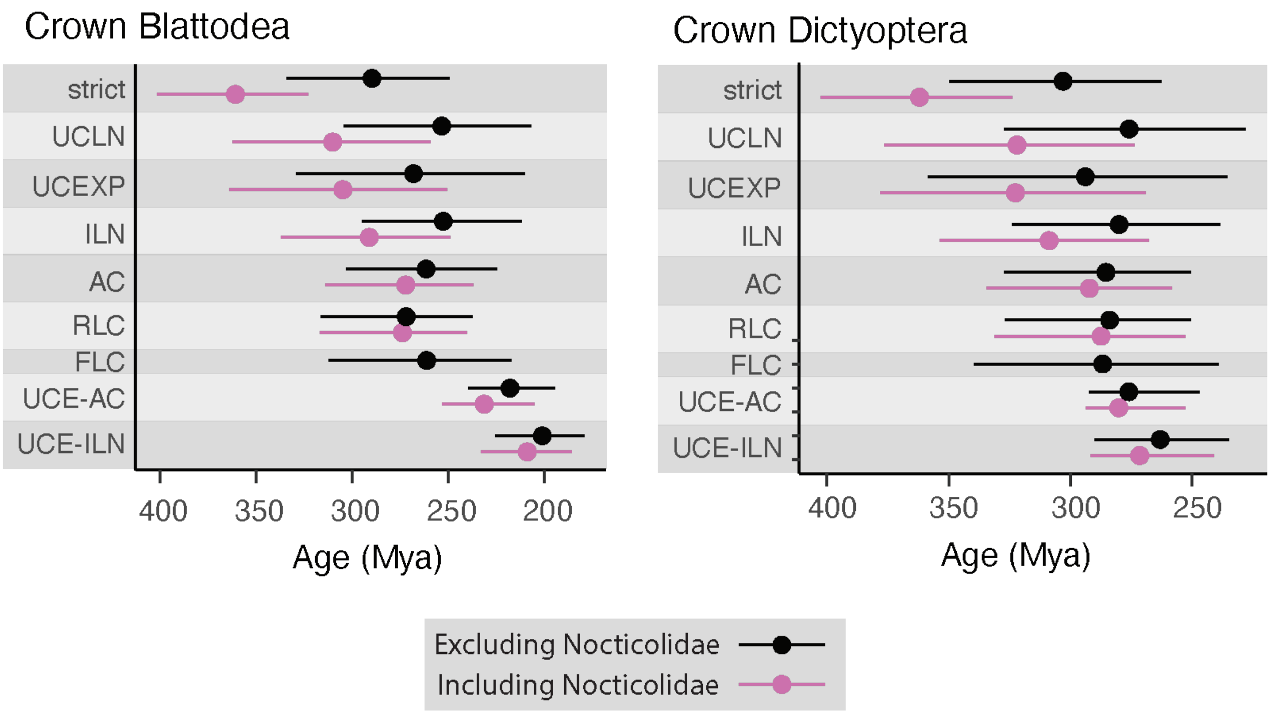


Supplementary Figure S20. Estimates of divergence times using a range of molecular-clock models when including (pink) and excluding (black) representatives of Nocticolidae from analyses. Dates are presented for crown Blattodea and crown Dictyoptera, estimated from mitochondrial protein-coding genes using a strict clock (strict), uncorrelated lognormal relaxed clock (UCLN), uncorrelated exponential relaxed clock (UCEXP), independent lognormal relaxed clock (ILN), autocorrelated relaxed clock (AC), random local clock (RLC), and flexible local clock (FLC). Date estimates for the same nodes using our UCE data set are also presented for comparison using an autocorrelated relaxed clock (UCE-AC) and independent lognormal relaxed clock (UCE-ILN). Circles indicate posterior means and bars indicate 95% credibility intervals.
