## Supplementary Material for "Dating in the Dark: Elevated Substitution Rates in Cave Cockroaches (Blattodea: Nocticolidae) Have Negative Impacts on Molecular Date Estimates"

### Extended Methods

#### Taxon Sampling

Samples of nocticolids were sought from a wide variety of locations across the known range of the family. Museum samples examined in this study were obtained from across northern Australia, Christmas Island, Singapore, and New Guinea (Supplementary Table S2). Sample age and condition varied, with the oldest sample collected in 1991. To capture the available genetic diversity in our samples, which has been found to be locally restricted in some Nocticolidae species (Trotter et al. 2017), we selected samples for DNA analysis that represented available locations.

In addition to the museum samples, we collected live individuals from field sites. During a visit to the Mungara-Chillagoe Caves National Park (MCCNP), 215 km west of Cairns, from 22 to 25 January 2020, we collected samples from Donna Cave, Royal Arch Cave, and Carpentaria Cave (just outside MCCNP). Within these caves, nocticolids were collected from under damp rocks using a small pooter. In the “Ballroom Fig” section of Royal Arch Cave, nocticolids were also collected from damp leaf litter (a small opening at the roof of this cavern appeared to act as a source of organic material). Because sample numbers were particularly small in Carpentaria Cave, slices of sweet potato were left overnight to attract additional nocticolids, which were collected the next day.

Previously, samples collected from Cutta Cutta Caves, NT, have been referred to as *N. brooksi* (eg. Djernæs et al. 2020). The description of *N. brooksi* included males, females, and nymphs collected from caves in the Kimberley, Western Australia, as well as females and nymphs collected >300 km away, from Cutta Cutta Caves, Northern Territory (Roth 1995). Females in Nocticolidae typically have relatively unspecialized morphology, and many species lack characters that enable them to be differentiated from females of other nocticolids (Trotter et al. 2017). Despite this, Roth (1995) classified specimens from these two locations as the same species despite noting that the females and nymphs from Cutta Cutta Caves were distinctly different (they lacked ommatidia, which were present in all samples, including nymphs, collected from the Kimberley). Thus, it is likely that the samples collected from Cutta Cutta Caves are a different species from *N. brooksi*. Hence, we refer to genomic samples from this location as Nocticolidae sp. (Cutta Cutta Cave).

#### DNA Extraction, Amplification, and Sequencing

DNA was extracted from a total of 113 specimens (Supplementary Table S2) using the DNeasy Blood and Tissue kit (Qiagen; Hilden, Germany), following the manufacturer’s protocol with the modifications outlined below. Due to the small size of specimens (adults ~5 mm long), whole individuals were destructively sampled for most samples, to maximize DNA yield during extractions. For unique specimens with no available paratypes, one hind leg was dissected and used for DNA extraction, such that the body could be retained for morphological analysis. Tissue was crushed using a sterile microtubule pestle and incubated with Proteinase K for 60 min at 56°C or until the tissue had completely lysed (some samples were left overnight). During the extraction process, extracts were eluted for 10 min in 100–200 µL of AE buffer. Smaller volumes of buffer were used for samples extracted from small amounts of tissue in order to maximize DNA concentration. DNA concentrations were measured using a Qubit fluorometer and a Qubit dsDNA HS Assay Kit for increased sensitivity. Samples with concentrations <0.5 ng/µL were discarded.

Polymerase chain reaction (PCR) amplification of a 440 bp fragment of the gene encoding mitochondrial 16S rRNA (*16S*) was attempted on all DNA samples using the primers LR-N-13398 (5’-CGCCTGTTTAACAAAAACAT-3'; forward) and LR-J-12961 (5’-TTTAATCCAACATCGAGG-3'; reverse) (Simon et al*.* 1994; Cognato and Vogler 2001). This marker was chosen because it has been used successfully in previous studies of cockroaches from a range of different families (Djernaes et al*.* 2012). PCR was performed in 200 µL microtubes in a solution of 25 µL containing 2 µL of DNA extract, 1 µL of each of the forward and reverse primers (10 µM), 8.5 µL of molecular grade water (G-Biosciences, Missouri, USA), and 12.5 µL of EconoTaq Plus 2x Master Mix (Lucigen, Wisconsin, USA). PCR involved an initial denaturation step (94°C for 2 min), followed by 44 cycles of denaturation (94°C for 30 s), annealing (45°C for 40 s), and extension (72°C for 1 min) (after 5 cycles, annealing temperature was increased to 48°C to limit off-target amplification for the remaining 39 cycles). The run was completed with a final extension step (72°C for 5 min). Following PCR, excess primers and nucleotides were removed using Exosap-IT (Thermo Fisher Scientific, Massachusetts, USA) following the manufacturer’s instructions and visualized on a 1% agarose gel to check product size and quality. Sanger sequencing of PCR products was performed by Macrogen (Seoul, South Korea) using primers used for amplification. Forward and reverse chromatograms were aligned, visually inspected, and manually trimmed in Geneious Prime 2020.0.5 ([www.geneious.com](http://www.geneious.com/)).

#### Mitochondrial 16S rRNA Sequence Alignment

In total, 113 nocticolid *16S* sequences were generated in this study (Supplementary Table S2). These were combined with six nocticolid sequences available on GenBank (Supplementary Table S3), as well as outgroup sequence data from 36 non-nocticolid cockroaches and termites and 5 other insects (Supplementary Table S4). We also assembled one *16S* sequence of *Nocticola* sp. (Malaysia) through mapping of transcriptome sequence data (Evangelista et al. 2019) against a nocticolid mitochondrial genome sequence (see below). All *16S* sequences were aligned in MAFFT v7.475 using the Q-INS-I McCaskill algorithm, which maximizes homology between sites by taking into account RNA secondary structures (Katoh et al., 2005; Katoh and Toh, 2008). This alignment was checked manually, and regions with poor sequence coverage or ambiguously aligned sites were removed.

#### Mitochondrial Genome and Nuclear Gene Sequence Data

We performed low-coverage high-throughput sequencing of 23 DNA samples (~5 Gb per sample; Supplementary Table S5). This approach has previously been used in studies of other cockroaches to sequence whole mitochondrial genomes and nuclear ribosomal RNA genes (Bourguignon et al. 2018; Beasley-Hall et al. 2021). A diverse range of nocticolid samples were selected for sequencing based on the phylogeny inferred from the *16S* data set and the availability of adequate DNA. Two samples from Christmas Island were included due to their low DNA concentration and the importance of sampling taxa from outside of the Australian mainland. Library construction and DNA sequencing was performed by BGI (Shenzhen, China). Specialised ‘low input’ libraries were constructed for some samples for which DNA extraction had produced a small yield.

Mitogenome assemblies were performed on all samples using the ‘map to reference’ function in Geneious Prime with the ‘Medium-Low Sensitivity/Fast’ setting. For each sample, all raw reads were mapped to the only available nocticolid mitogenome on GenBank (accession no. MG882221, Bourguignon et al. 2018). Next, for samples with high coverage (>10× coverage across >80% of the reference genome), these mapped reads were *de novo* assembled into contigs in Geneious Prime with the ‘Medium-Low Sensitivity/Fast’ setting. A contig of the appropriate size (~15 kb) was selected and IUPAC degenerate nucleotide codes were incorporated at positions where a single base was not present across >70% of assembled reads.

For some libraries (representing members of Nocticolidae B), the initial mapping step resulted in inadequate coverage and poor assemblies, most likely due to high divergence between the samples and the reference sequence. In these cases, we used NOVOplasty (Dierckxsens et al*.* 2017) to *de novo* assemble contigs from the entire raw read library, using the *16S* sequence produced by PCR for each sample as the starting ‘seed’. The output assembly from NOVOplasty was then used as the reference sequence for each sample, to which the raw reads were mapped following the method described above. Finally, the short-read archive for a previously generated transcriptome data set for *Nocticola* sp. (Malaysia) (Evangelista et al*.* 2019) was used as the starting library for the Geneious Prime ‘map to reference’ and *de novo* assembly method described above. Assembled genomes were ~15 kb and often lacked the AT-rich control region due to difficulties in assembly. All mitogenomes were annotated using web server (Donath et al*.* 2019) and reading frames were checked and adjusted manually. In total, we assembled 24 nocticolid mitogenomes. The mitogenome assembled from transcriptome data was incomplete, containing 11 of the 13 genes.

Eighty-seven outgroup taxa for phylogenetic analysis were selected from Bourguignon et al*.* (2018) to represent the genetic diversity across Blattodea (including representatives from each of the nine other families) and related insect lineages (Supplementary Table S6). We also selected taxa to maximize the specificity of the placement of relevant fossil calibrations for molecular dating. Nucleotide sequences for each of the 13 mitochondrial protein-coding genes were aligned at the amino acid level using MUSCLE through the TranslatorX web server (Abascal et al. 2010). Our final alignment of concatenated mitochondrial protein-coding genes (mtPCG) had a length of 10,590 bp, from 23 nocticolid taxa (one Christmas Island sample was dropped) and 87 outgroup taxa.

#### Tests of Sequence Saturation

To assess sequence saturation in our mtPCG data set, we analysed each of the 13 mitochondrial genes and the combined alignment using PhyloMAd (Duchêne et al. 2022). Based on entropy t-statistics, PhyloMAd detected a low risk of saturation effecting phylogenetic inference. However, the power of PhyloMAd to assess the effects of saturation using short sequence alignments like those produced for mitochondrial genomes is unknown. To investigate saturation further in our mtPCG data set, we compared uncorrected genetic distances with those corrected using the Tamura and Nei (1993) substitution model (Supplementary Fig. S12) implemented in the R package ape (Paradis and Schliep 2019). For 3rd codon positions, saturation was so high that a large proportion (35%) of corrected genetic distances between taxa were undefined, including almost all distances between two nocticolid taxa (92%) and the majority of distances between nocticolid taxa and non-nocticolid taxa (57%). Saturation was much less severe at 1st codon positions, although still present in pairwise distances including Nocticolidae (indicated by highly differing corrected and uncorrected distances), and negligible at 2nd codon positions. Hence, we removed 1st and 3rd codon positions in all analyses of mtPCG.

#### Generation of UCEs in silico

UCEs were identified using PHYLUCE v1.6.6 (Faircloth, 2016, 2017). A total of six genomes were used to design a cockroach UCE bait set. These included three publicly available genomes, *Blatella germanica* (GenBank accession no. GCA_003018175.1) (Harrison et al., 2018), *Periplaneta americana* (GCA_002939525.1) (Li et al., 2018) and the termite *Zootermopsis nevadensis* (GCF_000696155.1) (Terrapon et al., 2014), as well as three in-house genomes, *Geoscapheus dilatatus*, *Neogeoscapheus hanni*, and *Panesthia cribrata*. We used *B. germanica* as the base genome for UCE identification, due to its higher BUSCO score. We used the same bioinformatic protocol and parameters as in a previous study to generate a UCE data set for termites (Hellemans et al., 2022). We designed a preliminary bait set for a total of 116,064 loci shared between the base genome and up to four taxa. We removed UCEs with ambiguous base calls and GC-content above 70% or below 30%, as well as duplicates. The final set comprises a total of 342,038 baits targeting 32,386 UCE loci.

We extracted UCEs from a total of 61 samples, including 57 from Blattodea, one from Mantodea, and five from Phasmatodea (Supplementary Table S7). For each sample, reads were trimmed using fastp *v*0.20.1 (Chen et al. 2018), assembled with metaSPAdes *v*3.13 (Nurk, Meleshko, Korobeynikov, & Pevzner, 2017), and UCEs extracted using the PHYLUCE suite. Generated baits were aligned to assemblies at a minimum similarity threshold of 50% with phyluce_probe_run_multiple_lastzs_sqlite, and sequences with the flanking 200 bp situated at both the 5’ and 3’ ends were extracted using phyluce_probe_slice_sequence_from_genomes. Extracted sequences were filtered for duplicates and sequences matching multiple UCE loci using minimum identity threshold of 80% over 67% of bait length using phyluce_assembly_match_contigs_to_probes. Extracted loci were aligned using MAFFT and internal trimming was performed under default parameters with Gblocks (Castresana, 2000; Talavera & Castresana, 2007). Loci absent in more than 60% of taxa were filtered out. The baits and UCEs produced in the study are available from the Dryad Digital Repository: DRYAD_DOI_TO_BE_PROVIDED. UCE data obtained from this bait set will be centralized at the Cockroach UCE Database, available at: <https://github.com/sihellem/ROA-UCE-DB>.

#### Phylogenetic Analyses of 16S and mtPCG

We estimated phylogenetic relationships using each of the mitochondrial data sets (*16S*, mtPCG). Maximum likelihood (ML) analyses were performed using IQ-TREE v2 (Bui et al. 2020). The ModelFinder option within IQ-TREE v2 was used to select the best-fitting substitution models according to the Bayesian information criterion (Kalyaanamoorthy et al. 2017). For all nucleotide data sets, the best-fitting substitution model was GTR+G+I with four rate categories. Node support was assessed using the ultrafast bootstrap (UFBoot) approximation and the *SH-like* approximate likelihood-ratio test (10,000 replicates in each case) (Hoang et al. 2018; Guindon et al. 2010).

#### Phylogenetic Iinference Using Concatenated UCEs

For our data sets containing UCEs (UCE and UCE+mtPCG+*16S*), we selected the best-fit partitioning scheme by implementing the greedy algorithm of PartitionFinder to merge individual UCEs and mitochondrial genes to reduce over-parameterization (-m TESTMERGE). We allowed each partition to have a different evolutionary rate (-spp OR -p). This implements a model in which branch lengths are proportionate among subsets of the data, which has been found to result in the highest statistical support in many cases (Duchêne et al. 2020). The best-fitting substitution model was then selected for each partition using ModelFinder. Node support was assessed as per the *16S* analysis.

We performed Bayesian phylogenetic analyses of our mtPCG and UCE data sets using ExaBayes (Aberer et al. 2014). Analyses in ExaBayes using the concatenated UCE+mtPCG+*16S* did not converge and are not included in this study. Alignments were not partitioned in Bayesian analyses due to computational limitations. The GTR+G substitution model was used with a uniform prior on tree topology and an exponential prior on branch lengths. We ran Markov chain Monte Carlo simulations for 10^7^ steps, sampling every 5000 steps, in order to estimate the posterior distributions. Convergence between four independent runs and burn-in steps was assessed using the program Tracer v1.7.1 (Rambaut et al. 2018). One run was selected randomly, and the maximum-clade-credibility tree was calculated in TreeAnnotator v2.6.6 (Bouckaert et al. 2019).

#### Phylogenomic Analysis using the Summary-Coalescent Method

To account for gene-tree incongruence amongst UCE loci, we analysed the UCE data using the summary-coalescent method in ASTRAL-III (Zhang et al. 2018). This assumes that UCE loci are unlinked and that the individual gene trees are embedded in the underlying species tree. The gene tree for each UCE locus was estimated using IQ-TREE v2 (Bui et al. 2020) with substitution models selected using ModelFinder (Kalyaanamoorthy et al. 2017) and branch support assessed using UFBoot (Hoang et al. 2018). We repeated this analysis after contracting branches with less than 10% bootstrap support using Newick Utilities (Junier and Zdobnov 2010) which can improve inference of the species tree (Zhang et al. 2017). The inferred topologies were the same across both analyses (Supplementary Fig. S7). Without contracting branches, the inferred tree had a final normalized quartet score of 0.810, and this was only slightly increased after contracting branches, inferring a tree with a final normalized quartet score of 0.813.

*Testing Substitution Model Adequacy*

We assessed the adequacy of our nucleotide substitution models using PhyloMAD (Duchéne et al. 2018). We analysed the adequacy of the GTR model when analysing our mtPCG (unpartitioned) dataset using 100 simulations. This was completed for the dataset with and without the nocticolid taxa. We made our assessment based on the χ^2^ statistic, Multinomial statistic, Biochemical diversity, Consistency index and the squared Mahalanobis distance to summarise the risk across each metric. Unfortunately, each of our UCE partitions contained too much missing data to conduct the adequacy tests.

#### Estimating Evolutionary Timescales using BEAST

Evolutionary timescales were estimated for the mtPCG data sets using BEAST v2.5 (Bouckaert et al. 2019). For the mtPCG data set, only 2nd codon positions were used (3793 bases) to minimize saturation. Minimum bounds for node ages across Blattodea were chosen according to their earliest fossil representatives. We used a set of five fossils (Supplementary Table S9) conservatively selected by Evangelista et al*.* (2017, 2019) according to the criteria proposed by Parham et al. (2012), to inform the hard minimum bound of uniform priors. The recently discovered *Crenocticola burmanica* from Burmese amber was used as a sixth fossil, to provide a minimum age for stem-Nocticolidae (Li and Huang 2020). We followed previous studies in using the lack of winged insects in Rhynie chert before 412 Ma to infer a hard maximum bound for most of the fossil calibrations. An exception was the calibration based on *Archeorhinotermes rossi*, which was given a hard maximum bound of 237 Ma based on the absence of termites in the fossil record prior to 130 Ma, and their abundance in the record after 130 Ma (see Evangelista et al*.* (2019) for more information). We used a uniform prior for the age of the root with a hard maximum of 450 Ma based on estimates for the first established local terrestrial ecosystems, before which the diversification of insects is unlikely to have occurred (as used by Misof et al*.* 2014). Except for the sister groups to Nocticolidae and Dictyoptera, which are unresolved, we forced monophyly on all calibrating nodes based on relationships found previously (Bourguignon et al. 2018, Evangelista et al*.* 2019).

We used a birth-death tree prior to account for the mixture of intraspecific and interspecific sampling (Ritchie et al. 2017). Owing to the high rates of evolution among nocticolids detected in previous analyses, we investigated a range of different molecular clock models, including strict, random local, uncorrelated relaxed lognormal, uncorrelated relaxed exponential, and the flexible local clock (Drummond et al. 2006; Drummond and Suchard 2010; Fourment and Darling 2018). The flexible local clock provides a combination of the local and relaxed clocks by allowing multiple independent, uncorrelated relaxed clocks for subsets of a tree selected *a priori* (Fourment and Darling 2018). We implemented two independent uncorrelated lognormal relaxed clocks, one for Nocticolidae (including the stem branch), and one for the rest of the tree. MCMC simulations were run in duplicate for each of molecular clock models for 5×10^7^ steps, sampling every 10^3^ steps. Outputs were checked in Tracer for sufficient sampling of parameters. We generated the maximum-clade-credibility tree after removing a proportion of trees as burn-in and combining duplicate runs using LogCombiner.

To allow comparison of clock models, we estimated marginal likelihoods and their uncertainties using nested sampling with 20 particles, chain length of 3000, and convergence threshold (ε) of 10^-12^ (Maturana Russel et al. 2019). Marginal-likelihood estimates for the exponential relaxed clock had poor numerical stability, so runs were decreased to 10 particles and six replicates, choosing the highest log likelihood. We used Bayes factors to select the best-fitting clock model following the guidelines of Jeffreys (1961), which recommend that a model is strongly supported over another if the log_10_ Bayes factor is >2.3.

#### Estimating Evolutionary Timescales using MCMCtree

Evolutionary timescales were estimated for both the mtPCG and the UCE data sets using the approximate likelihood calculation in MCMCtree, part of the PAML package (Yang 2007). The analyses were performed using the topology from our maximum-likelihood analysis of each of the data sets. We used the GTR+G substitution model and ran analyses using both the independent (uncorrelated) lognormal relaxed clock and the autocorrelated relaxed clock. For mean substitution rate and the degree of rate drift, we specified gamma priors of “1 10” and “1 1”, respectively. We performed three independent MCMCtree analyses each for 5×10^7^ steps, sampling every 2500 steps, with 10% burn in. Outputs were checked in Tracer for sufficient sampling of parameters after convergence, indicated by effective sample sizes >200.

Our analysis of the mtPCG data set in MCMCtree used the same fossil calibrations as were employed in BEAST. For analysis of our UCE data set, we used the same fossils as implemented in our mtPCG analysis in BEAST with the exclusion of *Juramantophasma sinica* because the relevant node is not present in our UCE data set. Due to reduced outgroup sampling, we employed two secondary calibrations from a well resolved, transcriptomic, time-calibrated phylogeny of cockroaches and termites (Evangelista et a. 2019). The 95% credibility interval was used to infer the soft upper and lower bounds of a uniform prior on the crown age of Dictyoptera (232.2–291.2 Ma) and the soft upper bound for the root of the tree (Dictyoptera+Phasmatodea, 371 Ma) and all other fossil calibrations. An exception was the calibration based on *Archeorhinotermes rossi*, which was given a hard maximum bound of 237 Ma as was the case in our analysis of our mtPCG data set.

#### Estimating Root-to-Tip Distances

To investigate evolutionary rate variation between the families of Blattodea in our mtPCG and UCE data sets, we analysed root-to-tip distances in our maximum-likelihood trees after removing sister lineages to Blattodea. Mean root-to-tip distances were calculated for the two major clades of Nocticolidae and for each of the families of Blattodea using the R packages ape v5.5 (Paradis and Schliep 2019) and phangorn v2.7.0 (Schliep 2011).

*Analyses after removing Nocticolidae*

After removing representatives of Nocticolidae, we repeated our phylogenetic analysis in IQTREE, ExaBayes, BEAST and MCMCtree using the methods described above. The flexible local clock could not be included in analyses of the data set without Nocticolidae because without an independent relaxed clock in nocticolid lineages this model reverts to a regular relaxed clock model.

#### Metanocticola christmasensis

After inspecting the female paratype (WAM105862, previously BES 5828) of *Metanocticola christmasensis*, we have concluded that the “female” included in the description is actually a subadult. Therefore, the species requires a redescription containing a fully developed female and photos of the male. One of our samples was collected from the same cave (Jane-up Cave) as one of the paratypes included in the original description which is also closely located to the cave containing the holotype (Jedda Cave) (Roth, 1999). Therefore, it is likely that the samples from Christmas Island included here (Nocticolidae sp. (Christmas Island)) represent *Metanocticola christmasensis*, although this should be confirmed with further sampling of a male specimen. We also inferred that the Christmas Island lineages sampled here contain female cerci spines as they are present on two female specimens collected from the same location as one of our samples (Grants Well).

#### Morphology of Unsampled Nocticolidae

Based on the relationships estimated here, it is likely that members of Nocticolidae A and B represent members of at least two distinct genera. Broader sampling across Asia and Africa will potentially reveal further diversity within the family. We have identified some morphological characters that appear to align with relationships estimated from molecular data. We propose a potential synapomorphy of members of Nocticolidae A: a group of large sclerotized spines on the ventral surface of segments 4–6 of the female cerci (Roth 1988, Figure 1D; Trotter et al. 2017, Figure 5G–H). This has been observed in all available females of Nocticolidae A, and none of the observed females of Nocticolidae B. This character could provide some guidance for the placement of species without genetic data. For example, *Nocticola termitofila* (Vietnam) and *N. rohini* (=*Paraloboptera rohini*, Sri Lanka) have females with specialized spines on cerci (Silvestri 1946; Fernando 1962), suggesting a close relationship with Nocticolidae A, while *N. sinensis* (Hong Kong), *N. uenoi* (Japan), and *N.* sp. China do not (Silvestri 1946; Asahina 1974; Wang et al. 2017, Zongqing Wang personal communication), suggesting a close relationship to Nocticolidae B. Unfortunately, no females are known for the genus *Helmablatta*, but females in *Speleoblatta* which are likely to be closely related to *H. louisrothi*, do not have this morphology (Chopard 1932; Vidlicka et al. 2003, 2017).

There have been minimal collections of female Nocticolidae from Africa. This is because most studies have involved collecting flying insects in malaise traps and all nocticolid females are apterous or have severely reduced wings (Andersen and Kjaerandsen 1995). The type series of *N. jodarlingtonae* (Kenya) was collected from termite mounds and included females, but their description is basic and does not mention cerci (Roth 2003). Females of one species from Africa, *Alluaudellina cavernicola* do not contain specialised spines on cerci (Chopard 1932). This genus is likely a synonym of *Nocticola* (Chopard 1932, Roth 1988) and the lack of this morphology suggests that African Nocticolidae could be closely related to Nocticolidae Clade B.

Previous studies have found striking differences in male genitalia between closely related species in *Nocticola* (Trotter et al. 2017). However, broader inferences of relationships across the family based on morphology are limited by a historical lack of detailed drawings (more recent species descriptions contain higher-level detail, e.g., Trotter et al. 2017, Lucanas and Lit 2016). One genital character that aligns with our molecular results is the shape of L3d (left phallomere 3 dorsal), although data have not been collected for all taxa. The spear-like L3d of Nocticolidae B appears to be present in *N. gonzalezi* (Philippines) (Lucanas and Lit 2016, though labelled L2d) and the African species *N. clavata*, *N. scytala*, *N. wliensis*, and *A. cavernicola* (Andersen and Kjaerandsen 1995, Chopard 1932), although this should be confirmed by re-examination of type specimens because the drawings are old. The genitalia of *N. australiensis* and *N. babindaensis* do not align with either of these character types and require re-examination, because the drawings by Roth (1988) are basic. Unfortunately, the type species for the genus, *Nocticola simoni*, cannot be grouped into either clade due to its basic description (Bolivar 1892). Therefore, re-examination of *N. simoni* is required to identify whether Nocticolidae A or Nocticolidae B should remain as *Nocticola*, and which should be described as a new genus.

Further undescribed nocticolid species not included in this study include various taxa collected in Queensland, including from the Undara Lava Tubes (Eberhard and Howarth 2021), Walkunder Caves (Fred Stone, personal communication to J. A. Walker), Kiwi Cave (J. A. Walker personal collection), Mt Edith (collected by J. A. Walker), Mt Lewis (observed but not collected by J. A. Walker), Cammoweal (Roth 1988), and Crater Lakes National Park (Roth 1988).

Bolivar I. 1892. Voyage de M. E. Simno aux iles Philippines (mars et avril 1890). Etudes sur les Arthropodes cavernicoles de l’ile de Luzon. Orthopteres. Ann. la Société Entomol. Fr. 61:29–34.

Bolivar I. 1897. Nouvelle espèce cavernicole de la famille des blattaires. Viaggio di Leonardo Fea in Birmania e regioni vicine, LXXVIII. Annali Mus. civ. Stor. nat. “Giacomo Doria.” p. 32–36.

Bouckaert R., Vaughan T.G., Barido-Sottani J., Duchêne S., Fourment M., Gavryushkina A., Heled J., Jones G., Kühnert D., De Maio N., Matschiner M., Mendes F.K., Müller N.F., Ogilvie H.A., Du Plessis L., Popinga A., Rambaut A., Rasmussen D., Siveroni I., Suchard M.A., Wu C.H., Xie D., Zhang C., Stadler T., Drummond A.J. 2019. BEAST 2.5: An advanced software platform for Bayesian evolutionary analysis. PLoS Comput. Biol. 15:1–28.

Bourguignon T., Tang Q., Ho S.Y.W., Juna F., Wang Z., Arab D.A., Cameron S.L., Walker J., Rentz D., Evans T.A., Lo N. 2018. Transoceanic dispersal and plate tectonics shaped global cockroach distributions: Evidence from mitochondrial phylogenomics. Mol. Biol. Evol. 35:970–983.

Boussau B., Walton Z., Delgado J.A., Collantes F., Beani L., Stewart I.J., Cameron S.A., Whitfield J.B., Johnston J.S., Holland P.W.H., Bachtrog D., Kathirithamby J., Huelsenbeck J.P. 2014. Strepsiptera, phylogenomics and the long branch attraction problem. PLoS One. 9:e107709.

Brand P., Lin W., Johnson B.R. 2018. The draft genome of the invasive walking stick, Medauroidea extradendata, reveals extensive lineage-specific gene family expansions of cell wall degrading Enzymes in Phasmatodea. G3 Genes, Genomes, Genet. 8:1403–1408.

Bui Q.M., Schmidt H.A., Chernomor O., Schrempf D., Woodhams M.D., Von Haeseler A., Lanfear R. 2020. IQ-TREE 2: New Models and Efficient Methods for Phylogenetic Inference in the Genomic Era. Mol. Biol. Evol. 37:1530–1534.

Cameron S.L., Lo N., Bourguignon T., Svenson G.J., Evans T.A. 2012. A mitochondrial genome phylogeny of termites (Blattodea: Termitoidae): Robust support for interfamilial relationships and molecular synapomorphies define major clades. Mol. Phylogenet. Evol. 65:163–173.

Cameron S.L., Whiting M.F. 2007. Mitochondrial genomic comparisons of the subterranean termites from the genus Reticulitermes (Insecta: Isoptera: Rhinotermitidae). Genome. 50:188–202.

Castresana J. 2000. Selection of conserved blocks from multiple alignments for their use in phylogenetic analysis. Mol. Biol. Evol. 17:540–552.

Chen S., Zhou Y., Chen Y., Gu J. 2018. Fastp: an ultra-fast all-in-one FASTQ preprocessor. Bioinformatics. 34:i884–i890.

Chopard L. 1921. XXVII On some cavernicolous Dermaptera and Orthoptera from Assam. Rec. Indian Museum. 22:511–527.

Chopard L. 1924. On some cavernicolous Orthoptera and Dermaptera from Assam and Burma. Rec. Indian Museum. 206:383–399.

Chopard L. 1932. Un cas de microphtalmie liée à l’atrophie des ailes chez une blatte cavernicole. Livre du Centen. la Société Entomol. Fr. 485:485–496.

Chopard L. 1945. Note sur quelques Orthopteres cavernicoles de Madagascar. Rev. Fr. d’entomologie. 12:146–155.

Chopard L. 1950. Les blattes cavernicoles du genre Nocticola Bol. Eos, Tomo extraordinaire. p. 301–310.

Chopard L. 1966. Une espèce nouvelle de Nocticola provenant d’une grotte du Transvaal (Dictyoptères, Nocticolidae). Bull. la Société Entomol. Fr. 71:307–310.

Cognato A.I., Vogler A.P. 2001. Exploring data interaction and nucleotide alignment in a multiple gene analysis of Ips (Coleoptera: Scolytinae). Syst. Biol. 50:758–780.

Dierckxsens N., Mardulyn P., Smits G. 2017. NOVOPlasty: De novo assembly of organelle genomes from whole genome data. Nucleic Acids Res. 45:e18.

Djernæs M., Klass K.D., Eggleton P. 2015. Identifying possible sister groups of Cryptocercidae+Isoptera: A combined molecular and morphological phylogeny of Dictyoptera. Mol. Phylogenet. Evol. 84:284–303.

Djernæs M., Klass K.D., Picker M.D., Damgaard J. 2012. Phylogeny of cockroaches (Insecta, Dictyoptera, Blattodea), with placement of aberrant taxa and exploration of out-group sampling. Syst. Entomol. 37:65–83.

Djernæs M., Varadínova Z.K., Kotyk M., Eulitz U., Klass K.-D. 2020. Phylogeny and Life History Evolution of Blaberoidea (Blattodea). 78:29–67.

Donath A., Jühling F., Al-Arab M., Bernhart S.H., Reinhardt F., Stadler P.F., Middendorf M., Bernt M. 2019. Improved annotation of protein-coding genes boundaries in metazoan mitochondrial genomes. Nucleic Acids Res. 47:10543–10552.

Drummond A.J., Ho S.Y.W., Phillips M.J., Rambaut A. 2006. Relaxed phylogenetics and dating with confidence. PLoS Biol. 4:e88.

Drummond A.J., Suchard M.A. 2010. Bayesian random local clocks, or one rate to rule them all. BMC Biol. 8:114.

Duchêne D.A., Mather N., Van Der Wal C., Ho S.Y.W. 2022. Excluding Loci with Substitution Saturation Improves Inferences from Phylogenomic Data. Syst. Biol. 71:676–689.

Duchêne D.A., Tong K.J., Foster C.S.P., Duchêne S., Lanfear R., Ho S.Y.W. 2020. Linking branch lengths across sets of loci provides the highest statistical support for phylogenetic inference. Mol. Biol. Evol. 37:1202–1210.

Eberhard S.M., Howarth F.G. 2021. Undara Lava Cave Fauna in Tropical Queensland with an Annotated List of Australian Subterranean Biodiversity Hotspots. Diversity. 13:1–25.

Evangelista D.A., Djernæs M., Kohli M.K. 2017. Fossil calibrations for the cockroach phylogeny (Insecta, Dictyoptera, Blattodea), comments on the use of wings for their identification, and a redescription of the oldest Blaberidae. Palaeontol. Electron. 20.

Evangelista D.A., Wipfler B., Béthoux O., Donath A., Fujita M., Kohli M.K., Legendre F., Liu S., Machida R., Misof B., Peters R.S., Podsiadlowski L., Rust J., Schuette K., Tollenaar W., Ware J.L., Wappler T., Zhou X., Meusemann K., Simon S. 2019. An integrative phylogenomic approach illuminates the evolutionary history of cockroaches and termites (Blattodea). Proc. R. Soc. B Biol. Sci. 286:20182076.

Faircloth B.C. 2016. PHYLUCE is a software package for the analysis of conserved genomic loci. Bioinformatics. 32:786–788.

Faircloth B.C. 2017. Identifying conserved genomic elements and designing universal bait sets to enrich them. Methods Ecol. Evol. 8:1103–1112.

Fernando W. 1962. The gastric caeca and proventriculus in the Blattidae. Ceylon J. Sci. Biol. Sci. 4:88–95.

Fourment M., Darling A.E. 2018. Local and relaxed clocks: The best of both worlds. PeerJ. 2018:e5140.

Graverley F.H. 1910. XXIX. Alluaudella himalayensis, a new species of degenerate cockroach. Rec. Indian Museum. 3:307–311.

Graverley F.H. 1920. The female of the cockroach Alluaudella. 19:17–18.

Guindon S., Dufayard J.F., Lefort V., Anisimova M., Hordijk W., Gascuel O. 2010. New algorithms and methods to estimate maximum-likelihood phylogenies: Assessing the performance of PhyML 3.0. Syst. Biol. 59:307–321.

Harrison M.C., Jongepier E., Robertson H.M., Arning N., Bitard-Feildel T., Chao H., Childers C.P., Dinh H., Doddapaneni H., Dugan S., Gowin J., Greiner C., Han Y., Hu H., Hughes D.S.T., Huylmans A.K., Kemena C., Kremer L.P.M., Lee S.L., Lopez-Ezquerra A., Mallet L., Monroy-Kuhn J.M., Moser A., Murali S.C., Muzny D.M., Otani S., Piulachs M.D., Poelchau M., Qu J., Schaub F., Wada-Katsumata A., Worley K.C., Xie Q., Ylla G., Poulsen M., Gibbs R.A., Schal C., Richards S., Belles X., Korb J., Bornberg-Bauer E. 2018. Hemimetabolous genomes reveal molecular basis of termite eusociality. Nat. Ecol. Evol. 2:557–566.

Hellemans S., Wang M., Hasegawa N., Šobotník J., Scheffrahn R.H., Bourguignon T. 2022. Using ultraconserved elements to reconstruct the termite tree of life. Mol. Phylogenet. Evol. 173:107520.

Hoang D.T., Chernomor O., Von Haeseler A., Minh B.Q., Vinh L.S. 2018. UFBoot2: Improving the ultrafast bootstrap approximation. Mol. Biol. Evol. 35:518–522.

Huang D.Y., Nel A., Zompro O., Waller A. 2008. Mantophasmatodea now in the Jurassic. Naturwissenschaften. 95:947–952.

Huang M., Wang Y., Liu X., Li W., Kang Z., Wang K., Li X., Yang D. 2015. The complete mitochondrial genome and its remarkable secondary structure for a stonefly Acroneuria hainana Wu (Insecta: Plecoptera, Perlidae). Gene. 557:52–60.

Jarzembowski E.A. 1981. An early Cretaceous termite from southern England (Isoptera: Hodotermitidae). Syst. Entomol. 6:91–96.

Jeffreys H. 1961. The Theory of Probability. Oxford: Oxford University Press.

Jia Y.Y., Zhang L.P., Xu X.D., Dai X.Y., Yu D.N., Storey K.B., Zhang J.Y. 2019. The complete mitochondrial genome of Mantis religiosa (Mantodea: Mantidae) from Canada and its phylogeny. Mitochondrial DNA Part B Resour. 4:3797–3799.

Junier T., Zdobnov E.M. 2010. The Newick utilities: high-throughput phylogenetic tree processing in the UNIX shell. Bioinformatics. 26:1669–1670.

Kalyaanamoorthy S., Minh B.Q., Wong T.K.F., Von Haeseler A., Jermiin L.S. 2017. ModelFinder: Fast model selection for accurate phylogenetic estimates. Nat. Methods. 14:587–589.

Karny. 1924. Beiträge zur Malayischen Orthopteren fauna. Treubia. 5:12–19.

Katoh K., Kuma K.I., Toh H., Miyata T. 2005. MAFFT version 5: Improvement in accuracy of multiple sequence alignment. Nucleic Acids Res. 33:511–518.

Katoh K., Standley D.M. 2013. MAFFT multiple sequence alignment software version 7: improvements in performance and usability. Mol. Biol. Evol. 30:772–780.

Katoh K., Toh H. 2008. Recent developments in the MAFFT multiple sequence alignment program. Brief. Bioinform. 9:286–298.

Kômoto N., Yukuhiro K., Ueda K., Tomita S. 2011. Exploring the molecular phylogeny of phasmids with whole mitochondrial genome sequences. Mol. Phylogenet. Evol. 58:43–52.

Krishna K., Grimaldi D.A. 2003. The First Cretaceous Rhinotermitidae (Isoptera): A New Species, Genus, and Subfamily in Burmese Amber. Am. Museum Novit. 3390:1–10.

Legendre F., Nel A., Svenson G.J., Robillard T., Pellens R., Grandcolas P. 2015. Phylogeny of dictyoptera: Dating the origin of cockroaches, praying mantises and termites with molecular data and controlled fossil evidence. PLoS One. 10:1–27.

Legendre F., Whiting M.F., Bordereau C., Cancello E.M., Evans T.A., Grandcolas P. 2008. The phylogeny of termites (Dictyoptera: Isoptera) based on mitochondrial and nuclear markers: Implications for the evolution of the worker and pseudergate castes, and foraging behaviors. Mol. Phylogenet. Evol. 48:615–627.

Li S., Zhu S., Jia Q., Yuan D., Ren C., Li K., Liu S., Cui Y., Zhao H., Cao Y., Fang G., Li D., Zhao X., Zhang J., Yue Q., Fan Y., Yu X., Feng Q., Zhan S. 2018. The genomic and functional landscapes of developmental plasticity in the American cockroach. Nat. Commun. 9:1008.

Li X.R., Huang D. 2020. A new mid-Cretaceous cockroach of stem Nocticolidae and reestimating the age of Corydioidea (Dictyoptera: Blattodea). Cretac. Res. 106:104202.

Liu X.W., Zhu W.B., Dai L., Wang H.Q. 2017. Cockroaches of Southeastern China. Zhengzhou: Henan Science and Technology Press.

Lucañas C.C., Bláha M., Rahmadi C., Patoka J. 2021. The first Nocticola Bolivar 1892 (Blattodea: Nocticolidae) from New Guinea. Zootaxa. 5082:294–300.

Lucañas C.C., Lit I.L. 2016. Cockroaches (Insecta, Blattodea) from caves of Polillo Island (Philippines), with description of a new species. Subterr. Biol. 19:51–64.

Maturana Russel P., Brewer B.J., Klaere S., Bouckaert R.R. 2019. Model Selection and Parameter Inference in Phylogenetics Using Nested Sampling. Syst. Biol. 68:219–233.

Misof B., Liu S., Meusemann K., Peters R.S., Donath A., Mayer C., Frandsen P.B., Ware J., Flouri T., Beutel R.G., Niehuis O., Petersen M., Izquierdo-Carrasco F., Wappler T., Rust J., Aberer A.J., Aspöck U., Aspöck H., Bartel D., Blanke A., Berger S., Böhm A., Buckley T.R., Calcott B., Chen J., Friedrich F., Fukui M., Fujita M., Greve C., Grobe P., Gu S., Huang Y., Jermiin L.S., Kawahara A.Y., Krogmann L., Kubiak M., Lanfear R., Letsch H., Li Y., Li Z., Li J., Lu H., Machida R., Mashimo Y., Kapli P., McKenna D.D., Meng G., Nakagaki Y., Navarrete-Heredia J.L., Ott M., Ou Y., Pass G., Podsiadlowski L., Pohl H., Von Reumont B.M., Schütte K., Sekiya K., Shimizu S., Slipinski A., Stamatakis A., Song W., Su X., Szucsich N.U., Tan M., Tan X., Tang M., Tang J., Timelthaler G., Tomizuka S., Trautwein M., Tong X., Uchifune T., Walzl M.G., Wiegmann B.M., Wilbrandt J., Wipfler B., Wong T.K.F., Wu Q., Wu G., Xie Y., Yang S., Yang Q., Yeates D.K., Yoshizawa K., Zhang Q., Zhang R., Zhang W., Zhang Y., Zhao J., Zhou C., Zhou L., Ziesmann T., Zou S., Li Y., Xu X., Zhang Y., Yang H., Wang J., Wang J., Kjer K.M., Zhou X. 2014. Phylogenomics resolves the timing and pattern of insect evolution. Science. 346:763–767.

Nurk S., Meleshko D., Korobeynikov A., Pevzner P.A. 2017. metaSPAdes: a new versatile metagenomic assembler. Genome Res. 27:824–834.

Paradis E., Schliep K. 2019. Ape 5.0: An environment for modern phylogenetics and evolutionary analyses in R. Bioinformatics. 35:526–528.

Parham J.F., Donoghue P.C.J., Bell C.J., Calway T.D., Head J.J., Holroyd P.A., Inoue J.G., Irmis R.B., Joyce W.G., Ksepka D.T., Patané J.S.L., Smith N.D., Tarver J.E., Van Tuinen M., Yang Z., Angielczyk K.D., Greenwood J.M., Hipsley C.A., Jacobs L., Makovicky P.J., Müller J., Smith K.T., Theodor J.M., Warnock R.C.M., Benton M.J. 2012. Best practices for justifying fossil calibrations. Syst. Biol. 61:346–359.

Pellens R., D’Haese C.A., Bellés X., Piulachs M.D., Legendre F., Wheeler W.C., Grandcolas P. 2007. The evolutionary transition from subsocial to eusocial behaviour in Dictyoptera: Phylogenetic evidence for modification of the “shift-in-dependent-care” hypothesis with a new subsocial cockroach. Mol. Phylogenet. Evol. 43:616–626.

Piton E. 1940. Paléontologie du gisement éocène de Menat 1235 (Puy-de-Dôme) (flore et faune). Мémoires la Société d’Нistoire Nat. d’Auvergпe. 1:1–303.

Rambaut A., Drummond A.J., Xie D., Baele G., Suchard M.A. 2018. Posterior summarization in Bayesian phylogenetics using Tracer 1.7. Syst. Biol. 67:901–904.

Ritchie A.M., Lo N., Ho S.Y.W. 2017. The impact of the tree prior on molecular dating of data sets containing a mixture of inter- and intraspecies sampling. Syst. Biol. 66:413–425.

Roth L.M. 1988. Some cavernicolous and epigean cockroaches with six new species, and a discussion of the Nocticolidae (Dictyoptera: Blattaria). Rev. suisse Zool. 95:297–321.

Roth L.M. 1991. A new cave-dwelling cockroach from Western Australia (Blattaria: Nocticolidae). Rec. West. Aust. Museum. 15:17–21.

Roth L.M. 1995. New Species and Records of Cockroaches From Western Australia (Blattaria). Rec. West. Aust. Museum. 17:153–161.

Roth L.M. 1999. New cockroach species, redescriptions, and records, mostly from Australia, and a description of Metanocticola christmasensis gen. nov., sp. nov., from Christmas Island (Blattaria). Rec. West. Aust. Museum. 19:327–364.

Roth L.M. 2003. Some Cockroaches from Africa and Islands of the Indian Ocean , with Descriptions of Three New Species (Blattaria). Trans. Am. Entomol. Soc. 129:163–182.

Roth L.M., McGavin G.C. 1994. Two new species of nocticolidae (Dictyoptera: Blattaria) and a rediagnosis of the cavernicolous genus spelaeoblatta bolivar. J. Nat. Hist. 28:1319–1326.

Schliep K.P. 2011. phangorn: Phylogenetic analysis in R. Bioinformatics. 27:592–593.

Shelford R. 1908. XXVI. Some new Genera and Species of Blattidae, with Notes on the Form and the Pronotum in the Subfamily Perisphaeriinae. Ann. Mag. Nat. Hist. 8:157–177.

Shelford R. 1910. XIII. A new Cavernicolous Cockroach. Ann. Mag. Nat. Hist. 8:114–116.

Silvestri F. 1946. Prima nota su alcuni termitofili dell’Indocina. Boll. del Lab. di Entomol. Afraria Filippo. 6:313–330.

Silvestri F. 1947. Seconda nota su alcuni termitofili dell’Indocina con una appendice sul Macrotermes Barneyi Light. Boll. Lab. Ent. agr. 7:13–40.

Simon C., Frati F., Beckenbach A., Crespi B., Liu H., Flook P. 1994. Evolution, weighting, and phylogenetic utility of mitochondrial gene sequences and a compilation of conserved polymerase chain reaction primers. Ann. Entomol. Soc. Am. 87:651–701.

Song H., Moulton M.J., Whiting M.F. 2014. Rampant nuclear insertion of mtDNA across diverse lineages with in Orthoptera (Insecta). PLoS One. 9:41–43.

Strand E. 1928. Miscellanea nomenclatoria zoological et Paleontologica, I-II. Arch. für Naturgeschichte. 92:30–75.

Talavera G., Castresana J. 2007. Improvement of phylogenies after removing divergent and ambiguously aligned blocks from protein sequence alignments. Syst. Biol. 56:564–577.

Tamura K., Nei M. 1993. Estimation of the number of nucleotide substitutions in the control region of mitochondrial DNA in humans and chimpanzees. Mol. Biol. Evol. 10:512–526.

Terrapon N., Li C., Robertson H.M., Ji L., Meng X., Booth W., Chen Z., Childers C.P., Glastad K.M., Gokhale K., Gowin J., Gronenberg W., Hermansen R.A., Hu H., Hunt B.G., Huylmans A.K., Khalil S.M.S., Mitchell R.D., Munoz-Torres M.C., Mustard J.A., Pan H., Reese J.T., Scharf M.E., Sun F., Vogel H., Xiao J., Yang W., Yang Z., Yang Z., Zhou J., Zhu J., Brent C.S., Elsik C.G., Goodisman M.A.D., Liberles D.A., Roe R.M., Vargo E.L., Vilcinskas A., Wang J., Bornberg-Bauer E., Korb J., Zhang G., Liebig J. 2014. Molecular traces of alternative social organization in a termite genome. Nat. Commun. 5:3636.

Tokuda G., Isagawa H., Sugio K. 2012. The complete mitogenome of the Formosan termite, Coptotermes formosanus Shiraki. Insectes Soc. 59:17–24.

Trotter A.J., McRae J.M., Main D.C., Finston T.L. 2017. Speciation in fractured rock landforms: Towards understanding the diversity of subterranean cockroaches (Dictyoptera: Nocticolidae: Nocticola) in Western Australia. Zootaxa. 4250:143–170.

Vidlička L., Vršanský P., Kúdelová T., Kúdela M., Deharveng L., Hain M. 2017. New genus and species of cavernicolous cockroach (Blattaria, Nocticolidae) from Vietnam. Zootaxa. 4232:361–375.

Vidlička Ľ., Vršanský P., Shcherbakov D.E. 2003. Two new troglobitic cockroach species of the genus spelaeoblatta (blattaria: Nocticolidae) from North Thailand. J. Nat. Hist. 37:107–114.

Wang Z., Shi Y., Qiu Z., Che Y., Lo N. 2017. Reconstructing the phylogeny of Blattodea: Robust support for interfamilial relationships and major clades. Sci. Rep. 7:1–8.

Xiao B., Chen A.H., Zhang Y.Y., Jiang G.F., Hu C.C., Zhu C.D. 2012. Complete mitochondrial genomes of two cockroaches, Blattella germanica and Periplaneta americana, and the phylogenetic position of termites. Curr. Genet. 58:65–77.

Yang Z. 2007. PAML 4: Phylogenetic analysis by maximum likelihood. Mol. Biol. Evol. 24:1586–1591.

Ye F., Lan X.E., Zhu W.B., You P. 2016. Mitochondrial genomes of praying mantises (Dictyoptera, Mantodea): Rearrangement, duplication, and reassignment of tRNA genes. Sci. Rep. 6:1–9.

Zhang C., Rabiee M., Sayyari E., Mirarab S. 2018. ASTRAL-III: Polynomial time species tree reconstruction from partially resolved gene trees. BMC Bioinformatics. 19:15–30.

Zhang C., Sayyari E., Mirarab S. 2017. ASTRAL-III: Increased Scalability and Impacts of Contracting Low Support Branches. In: Meidanis J., Nakhleh L., editors. Lecture Notes in Computer Science. Springer. p. 53–75.

Zhang Z., Schneider J.W., Hong Y. 2013. The most ancient roach (Blattodea): A new genus and species from the earliest Late Carboniferous (Namurian) of China, with a discussion of the phylomorphogeny of early blattids. J. Syst. Palaeontol. 11:27–40.
